# Proximity labeling reveals cell cycle–specific NEK2 interactions and a regulatory axis controlling NUSAP1 stability

**DOI:** 10.64898/2026.01.25.701545

**Authors:** Enes Cicek, Selahattin Can Ozcan, Beste Kanevetci, Batuhan Mert Kalkan, Nazli Ezgi Ozkan, Nurhan Ozlu, Ceyda Acilan

## Abstract

NEK2 is a cell cycle–regulated kinase best known for its role in centrosome separation, yet the phase-specific organization of its interaction network has remained unclear. Here, we combine a doxycycline-inducible TurboID system with mass spectrometry to generate a cell cycle–resolved NEK2 interactome in synchronized U2OS cells. Using generalized additive models (GAMs), we identified different enrichment trajectories of the NEK2 interacting proteins across G1/S, late S, and G2/M, linking NEK2 to chromosome and spindle regulation, RNA–ribonucleoprotein processes, vesicle/lysosome compartments, and ubiquitin-associated pathways. Targeted validations (streptavidin pull-down, co-immunoprecipitation, and immunofluorescence) confirmed the interaction and binding for selected partners. Focusing on NUSAP1, NEK2 induction led to rapid loss of NUSAP1 protein without changes in mRNA levels, and this decrease was blocked by the proteasome inhibitor MG-132. Consistently, NUSAP1 exhibited slower decay in cycloheximide chase assays and reduced ubiquitination in NEK2 knockout cells, indicating NEK2-dependent proteasomal turnover. Global proteomic analysis of NEK2-deficient cells revealed widespread remodeling of protein abundance, including increased NUSAP1 and decreased KIF2C, accompanied by coordinated changes in pathways governing mitotic progression, microtubule organization, and ubiquitin-mediated protein turnover. Together, these findings provide a dynamic map of the NEK2 interactome across the cell cycle and uncover a NEK2–NUSAP1 degradation pathway, offering a framework to study how kinase interactomes are remodeled by cell cycle progression.

## 1. Introduction

NEK2A is a serine/threonine protein kinase that localizes to centrosomes, the nucleus, and the midbody^1^. Its expression is elevated in multiple cancer types, where it contributes to tumor progression and drug resistance^2^. NEK2 phosphorylates and inhibits p53 to promote cell division^3^ and plays a well-established role in centrosome separation prior to mitosis, thereby ensuring faithful chromosome segregation. Centrosomal linker proteins such as Rootletin^4^, C-Nap1^5^, CEP68^6^, and LRRC45^7^ are among its best-characterized phosphorylation targets. NEK2 is also required for the separation of supernumerary centrosomes^8^ and its activity is negatively regulated by the Anaphase-promoting complex through a KEN box motif in its C-terminus^9^. While NEK2 protein levels are tightly regulated throughout the cell cycle^10^, the broader network of NEK2 interactions and how this interactome is dynamically remodeled across cell cycle phases remains incompletely understood, especially in cancer.

Recent advances in proximity labeling methods, such as BioID and TurboID, have enabled the capture of protein interaction networks with spatial and temporal resolution^11,12^. These studies demonstrated that interactomes of proteins can change dynamically depending on cellular context, including transitions through the cell cycle^13,14^. Unlike conventional immuno-precipitation, which often misses transient or weak interactions, these approaches provide a powerful way to study dynamic and context-dependent protein networks in living cells. Given that NEK2 activity and abundance are tightly regulated by the cell cycle, resolving its interactome in a phase-specific manner is essential to understand how its function is coordinated by cell cycle progression. This strategy not only reveals regulatory partners of NEK2 but also provides a resource to explore how kinase interactomes adapt to distinct cellular states.

Recently, using proximity labeling, we identified KIF2C as a novel NEK2 interaction partner required for the inhibition of centrosome clustering in cancer cells with centrosome amplification^15^. This discovery suggested that NEK2 may engage in additional context-dependent interactions beyond centrosome separation, potentially linking it to other regulatory pathways. To investigate this, we performed a cell cycle–resolved interactome analysis of NEK2 using TurboID proximity labeling (Fig. 1A). This approach uncovered associations with proteins involved in centrosome biology, ubiquitination pathways, and DNA damage response. We also characterized how the NEK2 interactome and function change across cell cycle phases. Among these, we identified a previously unrecognized regulatory connection to the microtubule-binding protein NUSAP1, demonstrating that NEK2 promotes its proteasomal degradation. Our study therefore provides both a comprehensive NEK2 interactome resource and mechanistic insight into how this kinase shapes cell cycle–dependent regulation of cancer-relevant proteins.

**Figure 1.**
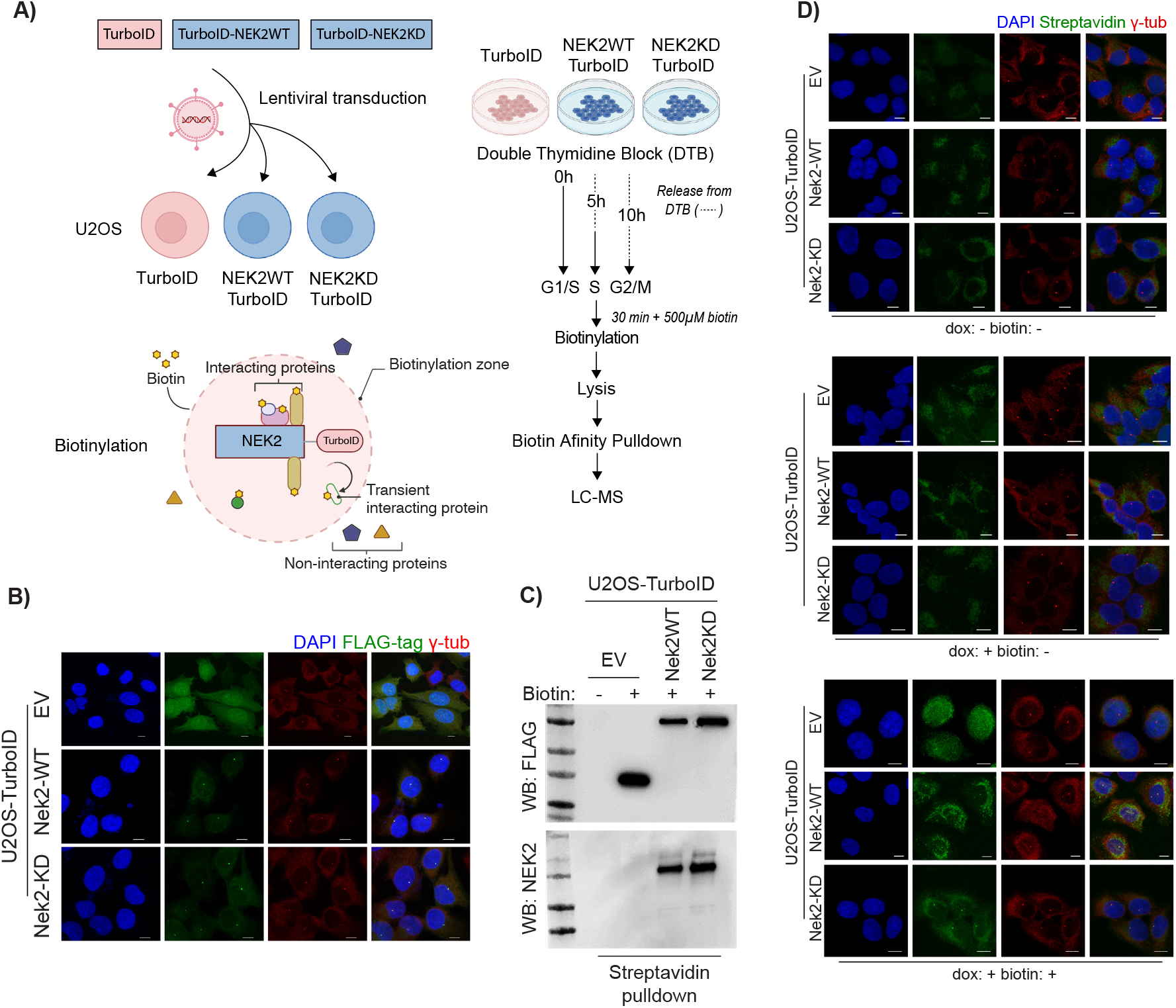
Generation and validation of the inducible TurboID–NEK2 proximity labeling system. (A) Schematic overview of the experimental design. (B) Intracellular localization of TurboID, TurboID–NEK2-WT, and TurboID–NEK2-KD proteins upon doxycycline induction. Blue: DAPI (DNA); Green: FLAG tag (TurboID-fusion proteins); Red: γ-tubulin (centrosomes). (C) Western blot analysis of FLAG and NEK2 after biotin labeling and streptavidin pull-down. (D) Localization of biotinylated proteins in U2OS–TurboID cell groups. Top: dox–/biotin–; Middle: dox+/biotin–; Bottom: dox+/biotin+. Blue: DAPI (DNA); Green: Streptavidin-488 (biotinylation); Red: γ-tubulin (centrosomes). EV: Empty vector. Scale bars show 10 μm distance.

## 2. Results

### 2.1 Generation of inducible TurboID-NEK2 proximity labeling system

We first generated doxycycline (dox)–inducible TurboID and TurboID–NEK2 constructs (wild-type, WT and kinase-dead, KD) (Fig. S1A) to examine the cell cycle–dependent changes in the NEK2 interactome. Inducible expression of the fusion proteins was confirmed by immunoblotting (Fig. S1B), and only minimal non-specific biotinylation was observed in the absence of dox and biotin treatment (Fig. S1C). Consistent with the localization of endogenous NEK2 at centrosomes throughout the cell cycle^1^, TurboID–NEK2 fusion proteins also localized to centrosomes, revealed by immunofluorescence microscopy (Fig. 1B). Upon induction of biotinylation, biotinylated proteins were efficiently enriched in streptavidin pull-down compared with whole-cell lysates (Fig. S1D). FLAG immunoblotting confirmed that enrichment occurred only after biotin treatment, and NEK2 antibodies validated the expression and self-biotinylation of the TurboID–NEK2 fusion proteins (Fig. 1C). In agreement, centrosomal biotinylation signals were detected only in dox-induced cells following biotin addition (Fig. 1D). Together, these results demonstrate the successful establishment of a dox-inducible TurboID–NEK2 proximity labeling system in U2OS cells.

Given that NEK2 is a centrosome-associated mitotic kinase^16^, its over-expression could potentially perturb centrosome function or cell cycle progression. To test this, we measured interphase centrosome distances and found no significant changes upon TurboID–NEK2 expression (Fig. S1E). Dox-induced NEK2 expression also did not cause centrosome amplification (Fig. S1F) or alter the cell cycle distribution (Fig. S1G). Thus, NEK2 over-expression did not interfere with centrosome integrity or cell cycle progression in our experimental conditions. Finally, to ensure the suitability of our system for the cell cycle–resolved experiments, we assessed the biotinylation after cell-cycle synchronization with a double thymidine block (DTB). Centrosomal biotinylation foci were observed in all phases of the cell cycle in both NEK2-WT and NEK2-KD constructs (Fig. S1H), demonstrating suitability of the experimental system.

### 2.2 Proximity labeling captures possible distinct roles of NEK2

To identify the cell cycle–dependent interactome of NEK2, we synchronized U2OS cells using a double thymidine block (DTB) and collected populations enriched in G1/S, late S, and G2/M phases (hereafter referred to as G1/S/G2 for simplicity) (Fig. 2A, Fig. S2A, Fig. S2B). Biotinylated proteins were isolated by streptavidin pull-down and analyzed by mass spectrometry. Fold-change values (log_2_FC, hereafter simplified as logFC) were calculated relative to TurboID-only control cells for each phase (Source Data Tables 1–4). Across all phases, we identified 165 proteins in NEK2-WT and 109 proteins in NEK2-KD samples that were significantly enriched (logFC > 1.5; adjusted p < 0.05) (Fig. 2B). As expected, NEK2 itself was the most enriched protein in all experimental groups (Fig. 2C). Comparison with reported NEK2 interaction partners in IntAct^17^ and BioGrid^18^ revealed 28 overlapping proteins, including CDCs, APC/C subunits, KIFC1, KIF24, ELAVL1, PSMD2, PRDX2, PDIA3, and HNRNPK (Fig. S2C; Source Data Table 5). Most of these interacting proteins displayed stable enrichment across phases (Fig. 2C), suggesting the existence of a core NEK2 interaction network that is largely preserved throughout the cell cycle. Strongly enriched groups included APC/C components (ANAPC4, ANAPC5, ANAPC16), centrosome- and microtubule-associated proteins (PCNT, KIF24, LRRC45, KIF2C, MAPRE3), and DNA repair factors such as MPG (Fig. 2C, Fig. S2D). The NEK2-KD interactome showed highly similar patterns (Fig. S2E). Gene set enrichment analysis (GSEA) of combined datasets highlighted ubiquitin-mediated proteolysis as the top pathway (Fig. 2D), which was further supported by KEGG analysis of the top 50 interacting proteins (Fig. S2F; Source Data Table 6).

**Table 1.**
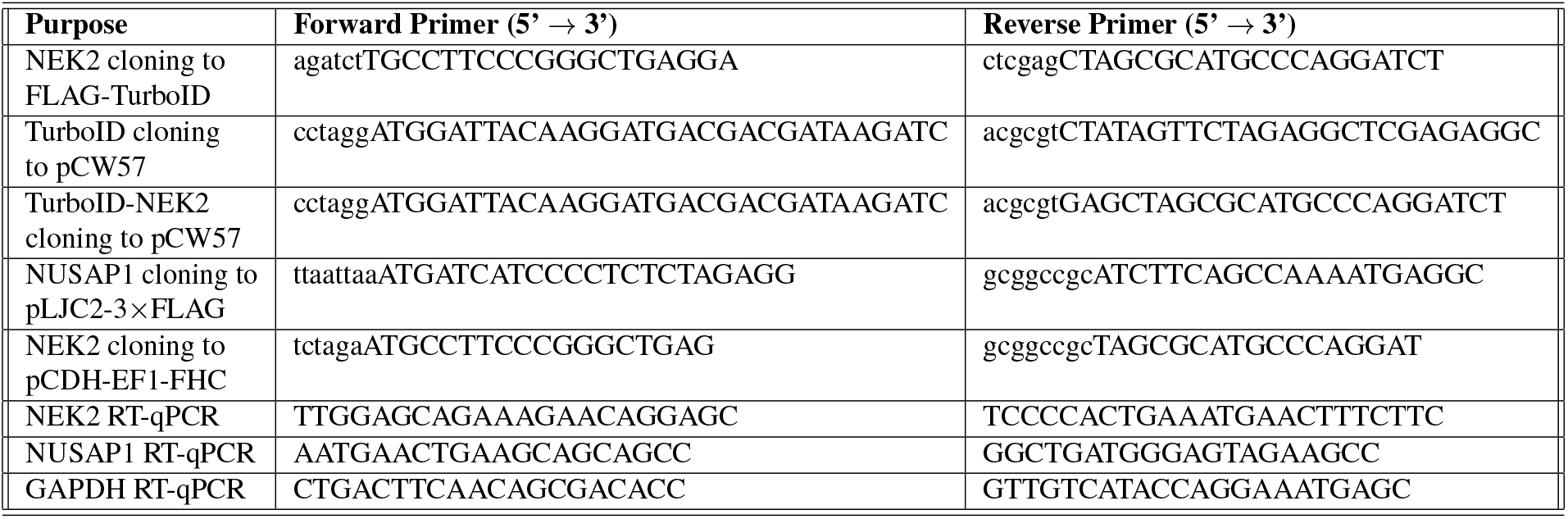
Supplementary Table - 1: Primer sequences used in this study.

**Figure 2.**
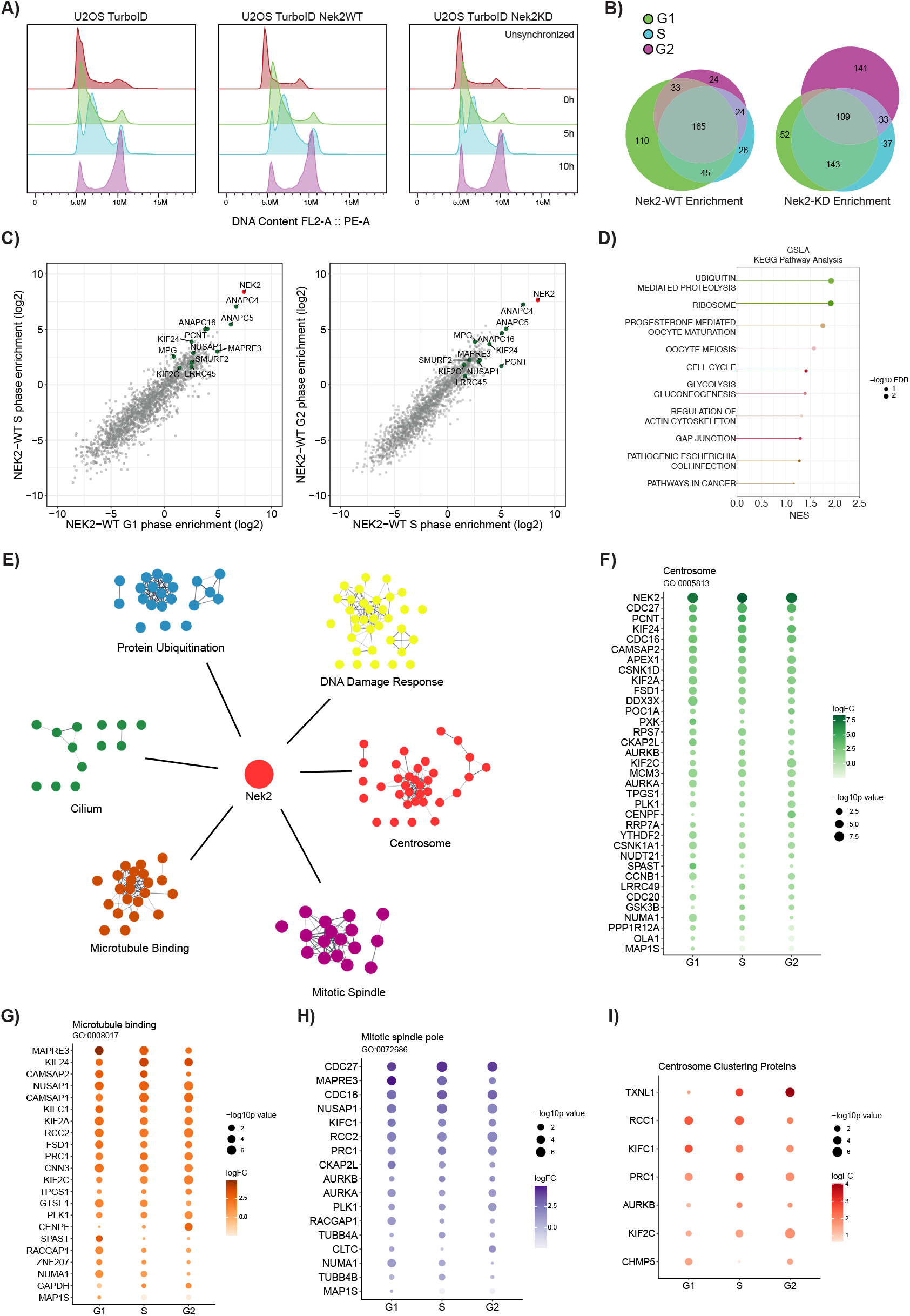
Proximity interaction analysis of NEK2 reveals roles in diverse cellular functions. (A) Cell cycle synchronization by double thymidine block (DTB) and DNA content distributions after 5h and 10h release. (B) Enriched proteins in different cell cycles in NEK2-WT and NEK2-KD proximity labeling experiments. (C) Distribution of logFC enrichment in the NEK2-WT interactome across G1, S, and G2 phases. Left side plot shows G1-S comparison, while right side plot shows S-G2 comparison. (D) Gene set enrichment analysis (GSEA-KEGG) of ranked NEK2 interaction partners. (E–I) Distribution of NEK2-WT biotinylation hits across functional categories. (E) Relationship patterns of NEK2 interaction partners across six selected GO categories. (F–I) Cell cycle–dependent enrichment of proteins associated with centrosomes (F), microtubule binding (G), mitotic spindle pole localization (H), and centrosome clustering (I).

Many of the strongest hits corresponded to previously established NEK2 interaction partners. For instance, CDC20 and APC/C proteins mediate NEK2 degradation prior to anaphase onset^10^; LRRC45, a centrosome linker protein, is phosphory-lated by NEK2 to regulate centrosomal tethering^7^; and phosphorylation of KIF24 by NEK2 is required for its microtubule-depolymerizing activity, which suppresses ciliogenesis^19^. We also previously identified KIF2C as a NEK2 binding partner essential for multipolar spindle formation in centrosome-amplified cells^15^, further validating the robustness of the current experimental approach.

To further evaluate the NEK2 interactome, we categorized proteins according to Gene Ontology (GO) functional and localization terms. Six GO categories were selected to group proteins by well-defined functions or localizations (Fig. 2E). Analysis of centrosome-associated proteins identified NEK2, CDC27, PCNT, KIF24, and CDC16 as top hits (Fig. 2F). While many showed stable enrichment across phases, several phase-specific differences was observed: (i) PCNT and CAMSAP2 were less enriched in G2, (ii) KIF2C and CENPF were more enriched in G2, and (iii) although overall enrichment was lower, NUMA1, PPP1R12A, OLA1, and MAP1S were more enriched in G1 (Fig. 2F). Among microtubule-binding and spindle-pole proteins, MAPRE3 was the most significant hit, with its highest enrichment observed in G1 (Fig. 2G, H). Proteins associated with cilia included

CFAP20, CSNK1A1, CSNK1D, and POC1A (Fig. S2G). As expected, APC/C components dominated ubiquitination-related terms, although other E3 ligases such as SMURF2 and TRIM3, and the E1 enzyme UBA1, were also prominent (Fig. S2H). Unexpectedly, multiple DNA damage response proteins including SIRT7, RAD9A, PARP2, and MPG were enriched in the NEK2 interactome, showing a dispersed enrichment trend through cell cycle phases (Fig. S2I). Finally, by integrating prior centrosome clustering studies^20,21^, we curated a gene list of centrosome clustering-related proteins (Source data table 7) and observed NEK2 interactions with TXNL1, RCC1, KIFC1, PRC1, AURKB, KIF2C, and CHMP5 (Fig. 2I). Among these, TXNL1 and KIF2C were preferentially enriched in G2 samples, suggesting a possible role in cell division related processes as NEK2-mediated multipolar spindle formation. Analysis of NEK2-KD enrichment yielded consistent results with minor differences (Fig. S3A–G).

### 2.3 Cell cycle phase specific hits suggest different functions through cell cycle

To better resolve the cell cycle–specific NEK2 interactome, we compared differential logFC enrichment patterns in TurboID–NEK2-WT samples (Fig. S4A, S4C). After filtering out proteins that were not significantly enriched (n = 1369) or enriched across all phases (n = 178), we obtained a dataset of 210 proteins with distinct enrichment patterns (Fig. S4B, S4C). Applying a combined p-value filter to identify the confident shared hits (see Methods) yielded 113 proteins that are classified as either unique or shared interaction partners (All phases shared: 178, G1 unique: 52, S unique: 12, G2 unique: 8, G1–S shared: 16, G1–G2 shared: 15, S–G2 shared: 10) (Fig. S4D, Fig. 3A). Their enrichment distributions showed characteristic patterns (ranging from two-to eightfold) in their representative phases compared with others, validating that the filtered set represents proteins with clear phase-specific enrichment (Fig. 3B). Because many hits were enriched in G1, we next examined their subcellular distribution, which revealed proteins linked to focal adhesions, ribonucleoprotein granules, chromosomes, and spindle structures (Fig. S4E).

**Figure 3.**
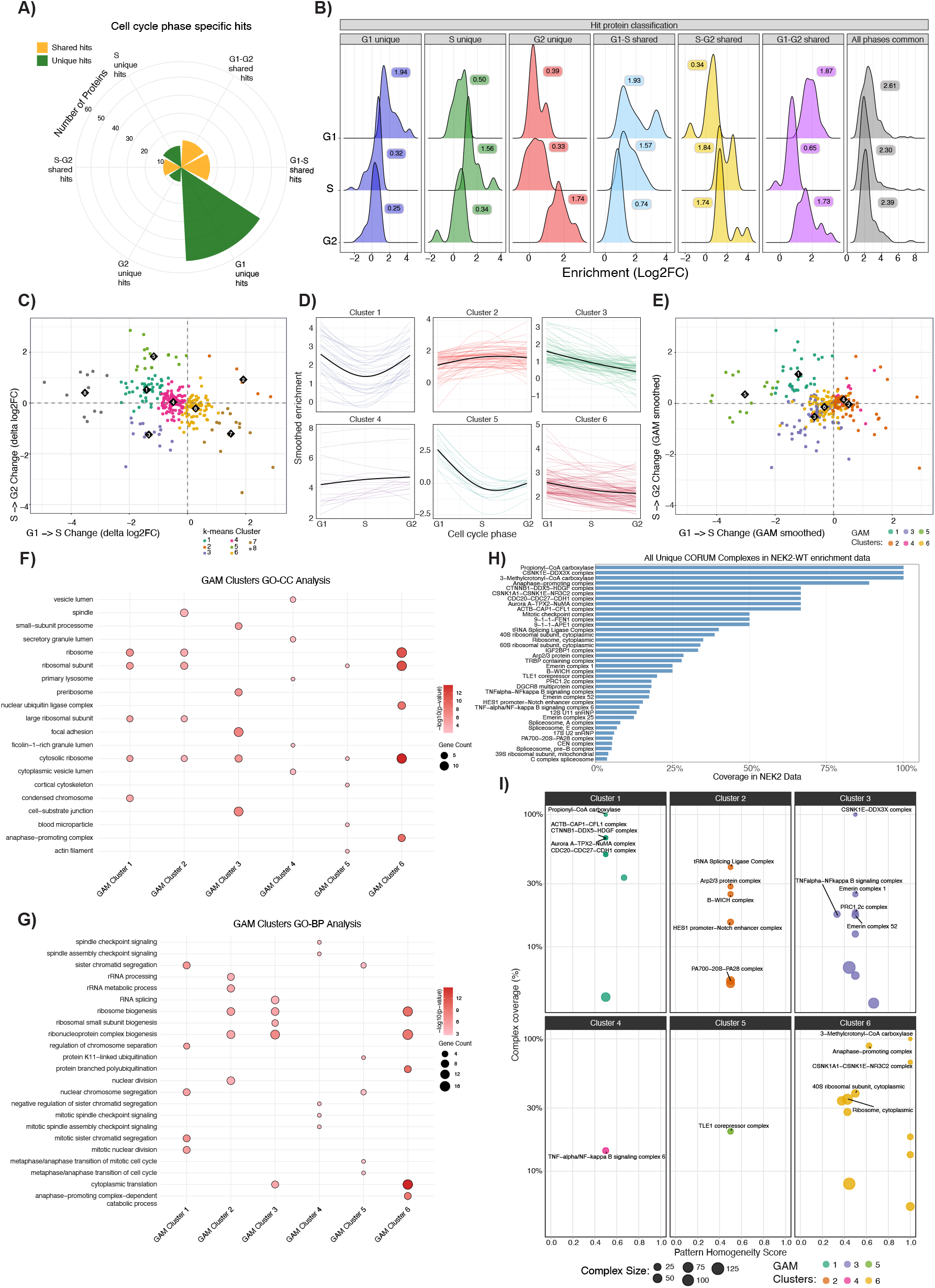
Cell cycle–specific functional interactions of NEK2. (A) Distribution of cell cycle–specific confident hits across different categories. (B) logFC enrichment of unique and shared hits in proximity labeling experiments. Mean enrichment of each group was shown on plots. (C) k-means clusters plotted according to changes in logFC enrichment from G1 to S and from S to G2 phases. (D) Protein enrichment dynamics across the cell cycle in GAM clusters. Colored lines represent GAM smoothed logFC values for individual proteins, and black lines indicate the mean trajectory. (E) GAM clusters plotted by changes in logFC enrichment from G1 to S and from S to G2 phases. (F–G) Functional enrichment analyses of GAM clusters: (F) GO-CC analysis and (G) GO-BP analysis. (H) Coverage of detected protein complexes in the NEK2 interactome. (I) Pattern homogeneity of protein complexes across GAM clusters.

To evaluate how the function and localization of NEK2 interaction partners vary across phases, we performed a GO-CC analysis to unique and shared hits. As expected, ribosomal proteins were enriched among shared hits, whereas mitotic spindle microtubule–associated proteins were enriched in G2-unique hits (Fig. S5A). Unexpectedly, G1-unique hits showed enrichment of kinetochore- and spindle-associated proteins, indicating that NEK2 already engages division-related partners before mitosis. G1–S shared hits were enriched for proteins associated with condensed chromosomes, nuclear specks, and U2-spliceosome complex, which is consistent with reports showing that NEK2 regulates alternative splicing through SRSF1^22^, even though SRSF1 itself was not detected in our dataset.

Motivated by these findings, we next analyzed how different groups of NEK2 interaction partners are regulated across cell cycle phases. To this aim, we first performed k-means clustering using both shared and phase-specific confident hits (see Methods). This approach revealed eight distinct clusters with clearly different enrichment dynamics across cell cycle phases. For example, Cluster 2 represented proteins with progressively higher enrichment from G1 to S and from S to G2, suggesting that their association with NEK2 increases steadily through the cell cycle (Fig. 3C). In contrast, Cluster 5 included proteins with lower enrichment in S compared with G1, but higher enrichment in G2 compared with S (Fig. 3C). When enrichment categories were evaluated across clusters, we observed a non-uniform distribution (Fig. S5B). For instance, Cluster 8 consisted exclusively of G1-unique proteins whose enrichment decreased in S relative to G1 but increased again in G2 relative to S (Fig. 3C). However, GO enrichment analysis of k-means clusters did not reveal strongly distinct functional patterns (data not shown here, but added to accompanying Zenodo folder^23^), suggesting that this approach was less sensitive in resolving biologically meaningful changes.

To overcome this limitation and more sensitively capture the dynamics of interaction changes between phases, we applied a generalized additive model (GAM) to NEK2 interactome dataset^24–26^ (see Methods). Unlike k-means clustering, which groups proteins by their absolute enrichment levels, the GAM fits a smoothed curve to the relative changes across the categorical phases, reducing bias toward any single phase and emphasizing temporal trends (Fig. 3D). This analysis identified six distinct categories of enrichment dynamics (Fig. 3D).

When enrichment categories were compared with k-means analysis, we observed that GAM produced a more uniform distribution of enrichment categories across different clusters (Fig. S5C). This reflects the fundamental differences between the two approaches: k-means groups proteins based on their initial ΔlogFC values and tends to be biased toward specific categories, whereas GAM smoothening emphasizes changes across time and provides a less biased view of enrichment dynamics. Plotting the clusters in phase-change space (G1-to-S versus S-to-G2) revealed three general dynamics in six clusters: some clusters dipped in S phase before recovering in G2, others gained enrichment in S phase with variable changes in G2, and a final group showed gradual loss across the entire cell cycle (Fig. 3D, 3E). To define their functional relevance, we next performed GO enrichment analyses, which revealed the following cluster-specific associations (Fig. 3F, G, Fig. S6):

- **Cluster 1** – Enrichment values increased from S to G2, consistent with late cell cycle functions. GO-CC terms highlighted condensed chromosomes and ribosomal subunits, while GO-BP terms included sister chromatid segregation and nuclear division. These findings indicate that proteins in the NEK2 interactome within this cluster mainly function in chromosome organization. Key proteins such as CDC16, CDC20, NUMA1, CENPF, RAD9A, RCC1, and CHMP5 were associated with cell division and mitosis-related terms (Fig. S6A), supporting a strong connection between NEK2 and regulators of mitotic progression.
- **Cluster 2** – Proteins in this cluster showed progressively higher enrichment in S phase compared with G1, with enrichment remaining similar in G2. GO-CC analysis identified spindle- and APC/C-associated proteins as well as ribosomal subunits, while GO-BP terms included ribosome biogenesis, rRNA metabolism, and spindle checkpoint signaling. NEK2 itself was located in this cluster. KIF24, MAPRE3, FSD1, KIF2A, and GTSE1 were among the proteins localized to microtubules and spindle poles, pointing to association of NEK2 with spindle-related structures (Fig. S6B). In addition, ACTR2, MUS81, SIRT7, KIF2A, and ANAPC7 were linked to nuclear division and chromosome segregation processes (Fig. S6B). Interaction of NEK2 with partners involved in spindle pole organization, checkpoint signaling, and chromosome segregation increased in S phase and were maintained through G2.
- **Cluster 3** – Proteins in Cluster 3 were most enriched in G1, with enrichment gradually declining as cells progressed through S and G2 phases. GO-CC analysis mapped these proteins to focal adhesions and cell–substrate junctions, while GO-BP terms highlighted RNA splicing and ribonucleoprotein biogenesis. Notably, DDX3X, TXNDC12, PDIA3, SEC61B, DNAJC10, and TBL2 were enriched in ER stress–related processes, and UBA1, SRP9, and SEC61B showed ER localization (Fig. S6C). In parallel, LUC7L2, LUC7L3, BUD31, and HNRNPK were associated with RNA splicing functions (Fig. S6C). This pattern suggests that NEK2 partners in Cluster 3 engage with RNA splicing machinery and may also contribute to ER stress–related pathways, pointing to an expanded interphase role for NEK2 in RNA splicing, protein folding and stress response mechanisms.
- **Cluster 4** – Proteins in Cluster 4 followed a pattern similar to Cluster 2, with enrichment increasing in S phase compared to G1 and in many cases continuing to rise in G2. GO-CC terms highlighted vesicle lumen and lysosomal compartments, while GO-BP processes included spindle checkpoint signaling and regulation of chromatid segregation. CPNE3, ANXA2, SRP14, and EEF2 were associated with lysosomes, vesicles, and secretory granules, whereas BUB3 and ZNF207 mapped to mitosis-related functions (Fig. S6D). Although this cluster contained fewer proteins and enrichment signals than others, it suggests that NEK2 may influence vesicle-mediated trafficking and lysosomal activity while simultaneously engaging with checkpoint regulators.
- **Cluster 5** – Proteins in Cluster 5 showed reduced enrichment in S phase followed by partial recovery in G2, a trend reminiscent of Cluster 1. GO-CC analysis identified associations with actin filaments, chromosomal regions, and chromosome segregation complexes, while GO-BP terms included the metaphase–anaphase transition, sister chromatid segregation, and K11-linked ubiquitination. Although this cluster contained relatively few proteins and enrichment signals, the combination of cytoskeletal and ubiquitination pathways points to a specialized role for NEK2 in these cellular processes.
- **Cluster 6** – Proteins in Cluster 6 showed a steady decline in enrichment in the NEK2 interactome from G1 to G2. GO-CC analysis highlighted ribosomes, APC/C subunits, and nuclear ubiquitin ligase complexes, while GO-BP terms included translation, ribosome biogenesis, and APC/C-dependent proteolysis. Interestingly, although the APC/C complex is known to mediate NEK2 degradation after the metaphase–anaphase transition, our analysis revealed that APC/C components remain associated with NEK2 throughout the entire cell cycle (Fig. S6E). This suggests that NEK2 maintains a continuous, possibly regulatory, interaction with APC/C beyond its canonical degradation window.

Taken together, these results indicate that NEK2 engages distinct sets of partners across the cell cycle. The six GAM clusters revealed enrichment patterns linked to chromosome organization, spindle regulation, adhesion complexes, RNA metabolism, vesicle and lysosomal compartments, cytoskeletal networks, and ubiquitin-mediated proteolysis. This integrated analysis highlights NEK2 as a kinase with multiple, spatially distinct roles, whose interactome is dynamically reorganized across cell cycle progression. Together, these analyses show that GAM resolves biologically meaningful, phase-specific patterns of kinase interactome dynamics.

### 2.4 NEK2 engages with protein complexes in temporally coordinated patterns

To determine whether NEK2 engages with entire protein complexes in a coordinated manner, we analyzed the NEK2-WT interactome dataset against the CORUM database of mammalian protein complexes^27^. This revealed 38 known complexes with at least two interacting subunits, exhibiting coverage ranging from 15% to 100% (Fig 3H). Strikingly, 26 complexes (68%) showed coordinated temporal interaction patterns, defined by a pattern homogeneity score *≥*0.5, indicating that most subunits within these complexes follow similar enrichment dynamics with NEK2 across the cell cycle.

Mapping these complexes to their dominant GAM clusters revealed distinct functional partitioning (Fig. 3I). Cluster 1 was enriched for mitotic complexes, including the Aurora-A–TPX2–NuMA complex (coverage = 67%) and the CDC20–CDC27–CDH1 complex (67%), consistent with its role in chromosome organization. Cluster 2 featured the tRNA splicing ligase complex (40%), aligning with RNA metabolism functions. Cluster 3 showed the CSNK1E–DDX3X complex (100%), supporting roles in both RNA processing and cell adhesion. Most notably, Cluster 6 contained the anaphase-promoting complex (APC/C) with high coverage (89%) but moderate homogeneity (0.63). This suggests that while multiple APC/C subunits interact with NEK2 throughout the cell cycle, they do so with varying kinetics.

This complex-level analysis demonstrates that NEK2 engages with functionally coherent protein modules in temporally coordinated patterns. The high pattern homogeneity observed for key complexes supports the biological relevance of our GAM-derived clusters and provides mechanistic insight into how NEK2 orchestrates distinct cellular processes across cell cycle progression.

### 2.5 NEK2 proximity labeling successfully identifies true interacting proteins

After analyzing the mass spectrometry dataset in detail, we selected several candidate proteins for biological validation. First, we confirmed the enrichment of two previously established NEK2 interaction partners, LRRC45 and KIF2C, both of which were enriched in TurboID–NEK2 streptavidin pull-down samples compared with TurboID-only control lysates (Fig. S7A).

Although the enrichment of LRRC45 and KIF2C was evident, some protein signal was also detectable in control samples, consistent with the high levels of non-specific biotinylation often observed in TurboID-only cells (Fig. S1C, S1E). To address this directly, we established an ELISA-based assay to quantify the total biotinylated protein content across conditions, providing an orthogonal normalization strategy. This analysis revealed that TurboID-only cells contained approximately five fold more biotinylated proteins than TurboID–NEK2 cells (Fig. S7B). Normalizing Western blot inputs to total biotinylated protein ratios, analogous to the internal normalization used in mass spectrometry analyses, allowed us to rigorously validate the enrichment of MAPRE3, NUSAP1, and MPG in the NEK2 interactome, in addition to LRRC45 and KIF2C (Fig. 4A). We next evaluated cell cycle–dependent enrichment of target proteins using DTB synchronization and observed increased biotinylation of NUSAP1 and MPG in G2/M-synchronized cells. This finding is consistent with our proteomics data, which showed increased enrichment of MPG during the S and G2 phases (Fig. 4B, Fig. S4A, Fig. S4B).

**Figure 4.**
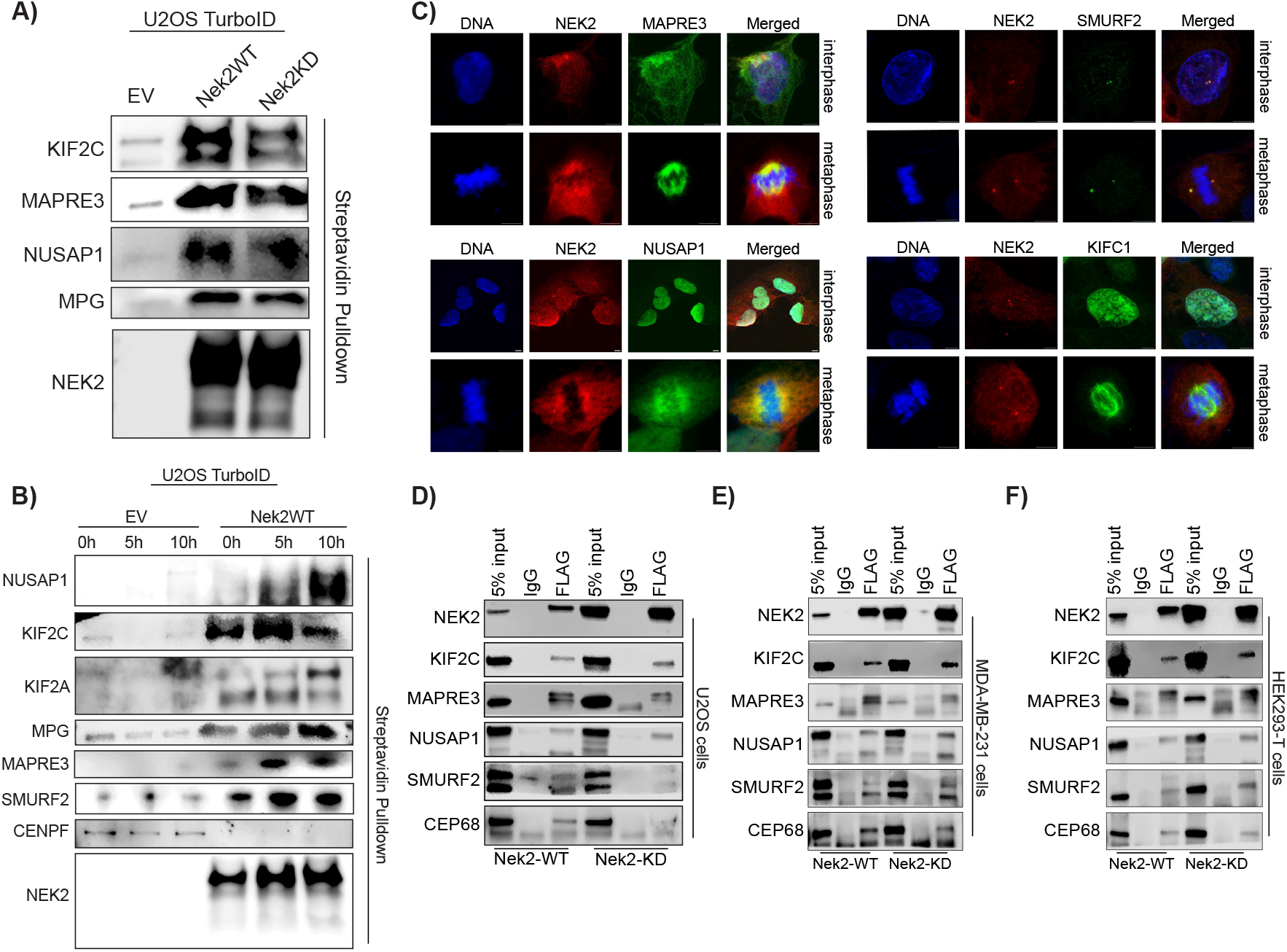
Targeted pull-down experiments validate mass spectrometry enrichment results for selected proteins. (A) Western blotting analysis of KIF2C, MAPRE3, NUSAP1, and MPG in unsynchronized samples. (B) Western blotting analysis of NUSAP1, KIF2C, KIF2A, MPG, MAPRE3, and SMURF2 in synchronized samples. (C) Immunofluorescence images showing co-localization of NEK2 with MAPRE3, NUSAP1, SMURF2, and KIFC1. (D–F) Co-immunoprecipitation (co-IP) validation of NEK2 interaction partners in (D) U2OS, (E) MDA-MB-231, and (F) HEK293T cells. CEP68 was used as a positive control for NEK2 interaction. EV: Empty vector. IgG: Immunoglobulin G. Scale bars show 5 μm distance.

To examine the spatial association of these proteins with NEK2, we performed immunofluorescence staining with specific antibodies. NUSAP1, MAPRE3, SMURF2, KIFC1, and MARK3 co-localized with NEK2 at centrosomes, spindle structures, or the nucleus (Fig. 4C, Fig. S7C). Given their close proximity to NEK2 in both biotin labeling and immunofluorescence imaging, we next tested whether these interactions were direct. Overexpression of FLAG-tagged NEK2-WT and NEK2-KD constructs (Fig. S7D), followed by co-immunoprecipitation, confirmed that NEK2 directly binds and interacts with MAPRE3, NUSAP1, and SMURF2. Importantly, these interactions were conserved across multiple cell lines, including U2OS, MDA-MB-231, and HEK293T cells (Fig. 4D–F).

Finally, we examined the cell cycle–regulated changes in endogenous protein levels in synchronized cells. NUSAP1 and NUMA1 levels increased from G1 to G2/M, KIF2C remained stable throughout the cell cycle, and NEK2 decreased after the G2 to M transition, consistent with regulation by the anaphase-promoting complex (Fig. S7E). In contrast, proximity-labeling proteomics revealed that NUMA1 and NUSAP1 maintained similar enrichment patterns across all cell cycle phases, whereas KIF2C showed stronger enrichment in G2. Thus, while NUMA1 and NUSAP1 protein levels rise toward mitosis but remain consistently associated with NEK2, KIF2C protein levels stay constant, yet its interaction with NEK2 becomes more pronounced in G2. This indicates that understanding the regulation of endogenous protein abundances is also important to consider when interpreting the cell cycle phase resolved interactome data.

### 2.6 NEK2 regulates the proteasomal degradation of NUSAP1

To evaluate the functional relationship between NEK2 and NUSAP1, we first examined how changes in NEK2 levels affect NUSAP1 abundance. In NEK2-KO cells, NUSAP1 protein levels were increased in two independent knockout clones (Fig. 5A). To determine whether this effect reflects a chronic adaptation to NEK2 loss or a direct response, we next reduced NEK2 expression by siRNA for 48 hours. Consistent with the knockout results, NUSAP1 levels also increased following short-term NEK2 depletion (Fig. 5A). Supporting this, doxycycline-inducible over-expression of NEK2 led to a rapid reduction in NUSAP1 protein levels within 30–60 minutes (Fig. 5B), without altering NUSAP1 mRNA expression (Fig. S7F). Interestingly, NUSAP1 abundance was partially recovered upon prolonged NEK2 induction, consistent with a fast and dynamic regulation of protein stability (Fig. 5B). To exclude potential confounding effects of doxycycline treatment itself, we treated parental U2OS cells lacking doxycycline-inducible expression cassettes with doxycycline for 1–2 h and observed no change in NUSAP1 protein levels (Fig. S7G). Together, these results indicate that NEK2 regulates NUSAP1 abundance at the protein level in a rapid and dynamic manner, independent of changes in NUSAP1 transcription.

**Figure 5.**
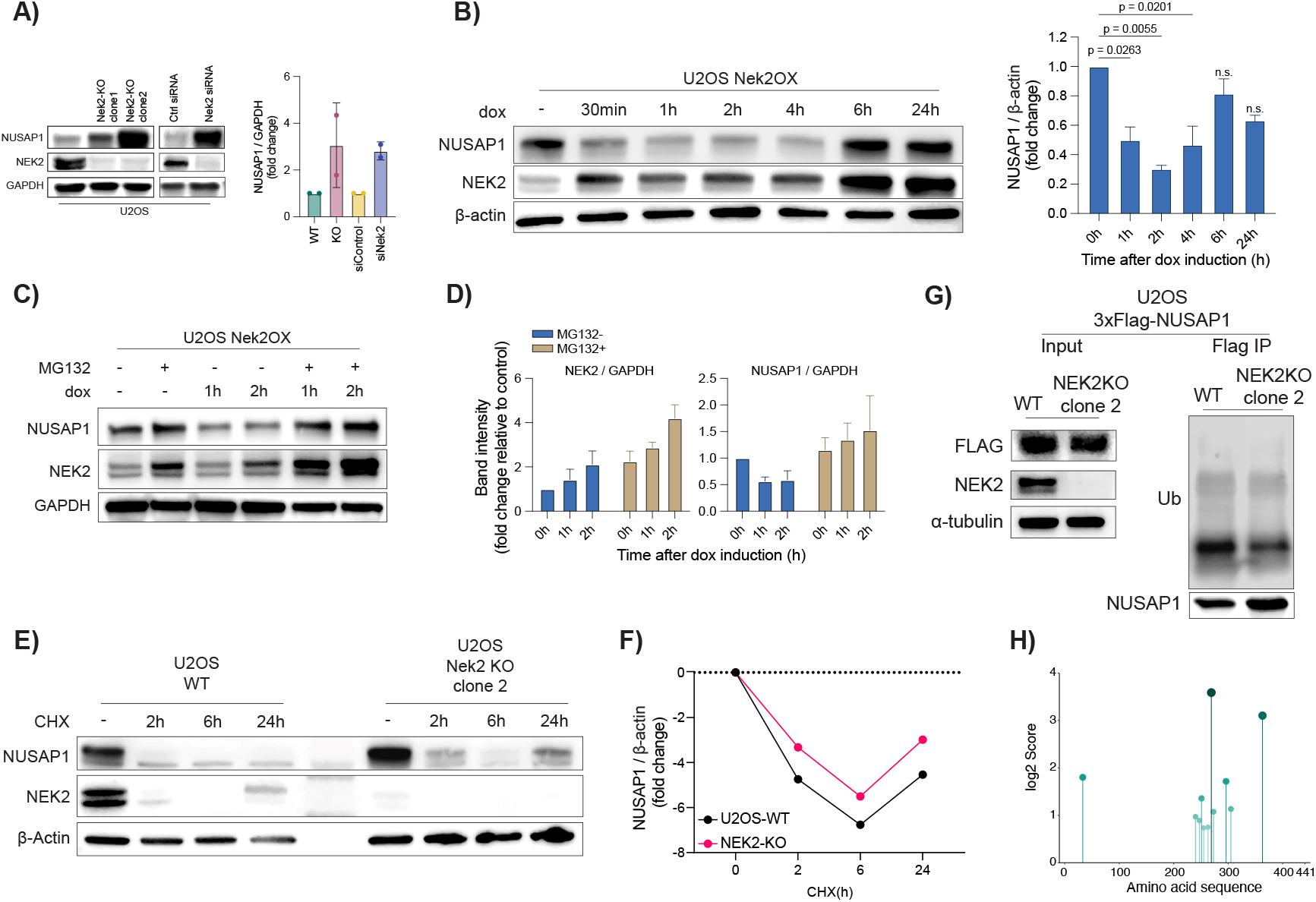
NEK2 regulates proteasomal degradation of NUSAP1. (A) Western blot analysis of NUSAP1 in NEK2-KO and NEK2 knockdown cells. (B) NEK2 overexpression decreases NUSAP1 levels. Left: Western blot results; Right: quantification of Western blot signals. Experiments were performed in two independent replicates; graphs show mean ± SD. Statistical significance was calculated using one-way ANOVA. p values are indicated on the graph. (C) NEK2-associated reduction of NUSAP1 is dependent on proteasome activity. (D) Quantification of panel C. Experiments were conducted in three independent replicates; graphs show mean ± SD. (E) Inhibition of translation with cycloheximide results in higher residual NUSAP1 levels in NEK2-KO cells compared with WT. (F) Quantification of panel E. (G) NEK2-KO results in reduced ubiquitination of NUSAP1. Left: Western blot showing FLAG–NUSAP1 expression in WT and NEK2-KO cells. Right: ubiquitin blot of FLAG pull-down samples. Note that although NUSAP1 protein levels were higher in NEK2-KO cells after pull-down, ubiquitination was decreased. (H) Prediction of putative NEK2-associated phosphorylation sites within the NUSAP1 protein sequence.

Building on these observations, we next tested whether the rapid, protein-level regulation of NUSAP1 by NEK2 reflects proteasome-mediated degradation. To this end, cells were treated with MG132, a well-established proteasome inhibitor. As observed previously (Fig. 5B), doxycycline-induced expression of NEK2 for 1–2 hours markedly reduced NUSAP1 protein levels; however, this reduction was completely abolished in MG132-treated samples (Fig. 5C, 5D), indicating that proteasomal activity is required for NEK2-dependent NUSAP1 down-regulation. To further assess whether NEK2 influences NUSAP1 protein stability, we inhibited *de novo* protein synthesis using cycloheximide (CHX) and monitored NUSAP1 levels over time in U2OS-WT and NEK2-KO cells (Fig. 5E). In WT cells, CHX treatment resulted in a progressive loss of NUSAP1 protein, whereas in NEK2-KO cells, NUSAP1 degradation was markedly attenuated, with residual protein persisting for longer periods despite translational blockade (Fig. 5E, Fig. 5F). Together, these results demonstrate that NEK2 promotes NUSAP1 turnover by accelerating its proteasome-dependent degradation.

To directly test whether NEK2 regulates the ubiquitination of NUSAP1, we generated a lentiviral 3×FLAG-NUSAP1 construct and over-expressed it in both U2OS-WT and NEK2-KO cells. FLAG immunoblotting confirmed comparable expression levels across conditions (Fig. 5G). We then performed FLAG immunoprecipitation of exogenous NUSAP1 and probed for K48-linked ubiquitination. Compared with WT cells, NUSAP1 immunoprecipitates from NEK2-KO cells displayed markedly reduced K48-linked ubiquitin signals (Fig. 5G). Furthermore, using a Nek2 phosphorylation consensus sequence (see Methods), we predicted putative phosphorylation sites on NUSAP1 and identified two high-confidence serine residues, S269 and S363 (Fig. 5H), suggesting that NEK2 may target and phosphorylate these residues to mark NUSAP1 for ubiquitination. Together, these results show that NEK2 promotes NUSAP1 ubiquitination and thereby regulates its stability through the proteasomal degradation pathway.

### 2.7 NEK2 regulates intracellular levels of interacting proteins

After identifying the negative regulation of endogenous NUSAP1 levels by NEK2, we next examined how other proteins in the NEK2 interactome are influenced by NEK2. To this end, we performed a global proteomics analysis comparing protein abundances in NEK2-KO versus U2OS wild-type cells. This analysis revealed widespread changes, with numerous proteins differentially regulated in the absence of NEK2 (Fig. S8A, S8B). Cross-referencing the results with previously reported interaction partners of NEK2 in the BioGRID and IntAct databases, together with our TurboID-derived hits, identified 68 significantly (logFC > 0.5 & adjusted p < 0.05) down-regulated and 92 up-regulated proteins in NEK2-KO compared with WT cells (Fig. 6A). Consistent with our earlier findings, KIF2C was down-regulated and NUSAP1 was up-regulated. Among the top up-regulated proteins were CRKL, HNRNPF, NPM1, and LGALS1, while FXR1, LTN1, and RPS13 were among the most down-regulated. GO enrichment analysis of up-regulated proteins highlighted processes such as nuclear export and mRNA splicing, whereas down-regulated proteins were enriched in mitotic nuclear division, chromosome segregation, microtubule organization, and ubiquitin-mediated protein turnover (Fig. S8C). These results demonstrate that NEK2 not only regulates individual substrates such as NUSAP1 but also broadly shapes intracellular protein levels.

**Figure 6.**
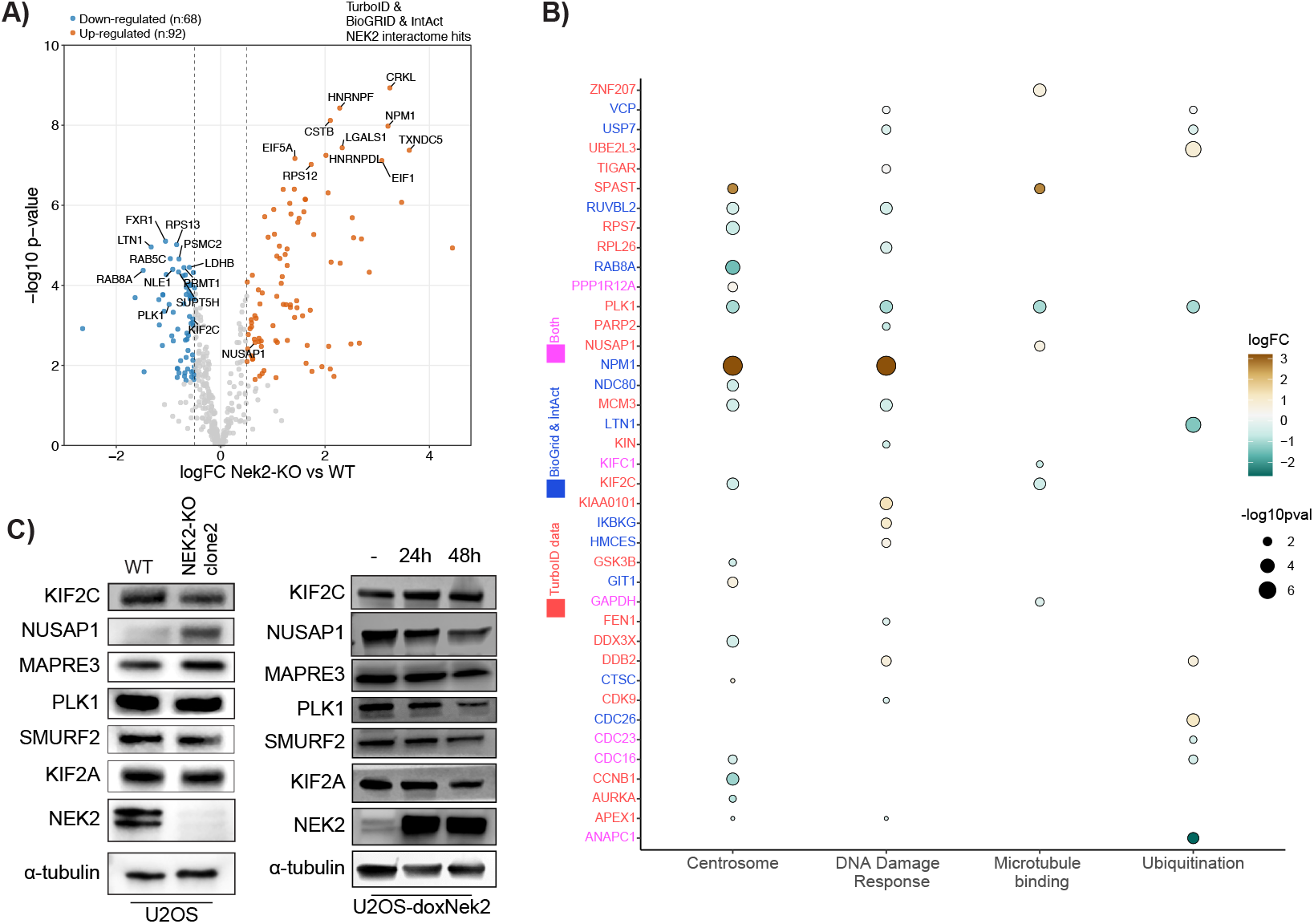
NEK2 knockout broadly affects levels of NEK2 interaction partners. (A) Volcano plot showing protein abundance changes upon NEK2 knockout. The dataset was filtered for previously reported (BioGRID & IntAct) and TurboID-identified NEK2 interaction partners. (B) Protein abundance changes in selected groups of proteins associated with representative GO terms. (C) Western blot analysis showing the effects of NEK2 knockout and doxycycline-induced overexpression (24 h and 48 h) on protein levels of selected NEK2 interaction partners.

To better resolve the impact on specific interaction partners, we filtered the dataset for proteins localizing to centrosomes or involved in DNA damage response, microtubule binding, and ubiquitination; GO-terms that together capture central functions of NEK2. Within this subset, the most enriched protein in NEK2-KO samples was NPM1, while ANAPC1, an APC/C subunit associated with ubiquitination, was among the most depleted (Fig. 6B).

We then directly examined how NEK2 level changes affect the abundance of selected interaction partners. In agreement with the global proteomics results, NEK2 knockout was associated with reduced KIF2C levels and increased NUSAP1 levels (Fig. 6C). Conversely, doxycycline-induced NEK2 overexpression for 24 and 48 hours decreased NUSAP1 levels and increased KIF2C levels (Fig. 6C). In contrast, MAPRE3, PLK1, and SMURF2 remained largely unchanged regardless of NEK2 expression. Together, these findings validate KIF2C and NUSAP1 as direct regulatory targets of NEK2, while suggesting that not all interacting proteins are equally sensitive to changes in NEK2 abundance.

## 3 Discussion

NEK2A has long been recognized as a centrosome-associated kinase critical for centrosome separation and spindle organization^4,5,7^. Here, we extend this classical view by providing a cell cycle–resolved map of the NEK2 interactome and by functionally linking NEK2 to the proteasome-dependent regulation of the spindle-associated factor NUSAP1. Through proximity labeling combined with mass spectrometry, we identified both previously validated partners, such as APC/C components and KIF24, and novel interacting proteins, including NUSAP1, MAPRE3, MPG, and SMURF2. These findings broaden the functional scope of NEK2, positioning it as a coordinator of multiple pathways across the cell cycle.

Our study identifies NUSAP1 as a direct substrate regulated by NEK2. NUSAP1 is a microtubule-binding factor essential for spindle assembly and chromosome segregation, and its overexpression has been linked to aggressive tumor phenotypes^28–30^. Notably, NUSAP1 also modulates the microtubule-depolymerizing activity of KIF2C, another protein identified in the NEK2 interactome^31^. We observed that NEK2 overexpression reduced NUSAP1 levels but increased KIF2C levels, suggesting a linked regulatory axis. Since KIF2C is required for NEK2-driven multipolar metaphase formation^15^, NUSAP1 down-regulation may cooperate with increased KIF2C abundance to promote this phenotype.

Interestingly, recent research has also characterized NEK2 as a regulator of proteasomal degradation of OLA1^32^. Although the enrichment of OLA1 was relatively low in our dataset, we detected it in the NEK2 interactome (Fig. 2F). In that study, NEK2 was shown to phosphorylate OLA1 and promote its polyubiquitination by Aurora A, which was identified for the first time as a polyubiquitinating factor. In our work, we could not resolve the precise mechanism underlying NUSAP1 ubiquitination, but it is conceivable that Aurora A may also act downstream of NEK2 phosphorylation. Together, these observations suggest that phosphorylation by NEK2 may act more generally as a trigger for protein degradation, since several established NEK2 substrates including centrosomal linker proteins, as well as OLA1 and NUSAP1, are phosphorylated and subsequently directed to proteolysis.

The interactome analysis also revealed enrichment of several DNA damage response factors (e.g., PARP2, SIRT7, MPG), pointing to additional testable connections between NEK2 and genome maintenance. While NEK2 has long been linked to chromosomal instability in cancer^2,3^, our results suggest a broader regulatory interface with DNA repair machinery. Similarly, the presence of ubiquitin ligases and APC/C subunits is consistent with prior reports that levels of NEK2 are tightly controlled by ubiquitin-mediated turnover^9,10^, and our dataset indicates that APC/C subunits remain proximal to NEK2 across the cell cycle, consistent with sustained regulatory contact.

Proximity labeling further reinforced the connection of NEK2 to centrosome clustering. We previously reported that NEK2 interacts with KIF2C to promote multipolar spindle formation in cells with centrosome amplification^15^. Here, we extend this observation by identifying additional clustering-related proteins, including TXNL1, RCC1, and CHMP5. Notably, enrichment of KIF2C interactions increased in G2 relative to G1 and S, and similar patterns were seen for TXNL1 in NEK2-WT interactome and MYCBP2 and ANLN in NEK2-KD interactome. These findings suggest that regulation of centrosome clustering by NEK2 may extend beyond KIF2C to include additional interacting proteins such as TXNL1, MYCBP2, and ANLN.

From a cancer biology perspective, our findings emphasize NEK2 as a dual regulator of centrosome organization and protein stability. Over-expression of NEK2 has been correlated with poor prognosis, drug resistance, and enhanced proliferation in tumors^1,2^. By combining proximity labeling with cell cycle resolution and GAM-based trajectory analysis, we provide a dynamic resource that maps when and where NEK2 engages functional modules—including chromosome organization, spindle regulation, adhesion and RNA metabolism, vesicle/lysosome pathways, and ubiquitin systems. The NEK2–NUSAP1 axis described here highlights a tractable regulatory node that could be exploited in contexts of chromosomal instability.

Methodologically, we refined proximity labeling in three ways. First, we used a doxycycline-inducible TurboID system in dialyzed FBS, minimizing the background biotinylation^33^. Second, we included a matched-expression TurboID-only control group, in line with recommendations that matched bait and control expression levels are essential to reduce false positives and negatives in interactome studies^34^. Third, we added an orthogonal normalization step to validation experiments by quantifying the total biotinylated protein levels via ELISA, which improved validation of the enrichments by Western blot. Together, these refinements increase the robustness and reproducibility of proximity labeling datasets.

Limitations of the study include our reliance on U2OS cells, which may not capture cell type–specific differences in NEK2 interaction networks or regulatory outputs, and the fact that TurboID proximity labeling identifies spatially proximal proteins without distinguishing direct phosphorylation targets from indirect or transient associations. In addition, while our data establish that NEK2 regulates NUSAP1 abundance through proteasome-dependent mechanisms, we did not define the precise molecular determinants underlying this process. Specifically, functional mapping of NEK2-dependent phosphorylation sites on NUSAP1, identification of the E3 ubiquitin ligase(s) responsible for NUSAP1 ubiquitination, and comprehensive profiling of ubiquitin chain linkages were beyond the scope of this study. Future studies integrating phosphoproteomics, degron mapping, targeted perturbation of candidate ubiquitin ligases, and complementary interaction-mapping approaches will be required to fully delineate how NEK2 kinase activity is coupled to proteostasis regulation across the cell cycle.

In conclusion, this study provides the first cell cycle–resolved map of the NEK2 interactome and identifies a previously unrecognized regulatory pathway by which NEK2 promotes proteasome-dependent degradation of NUSAP1. Together, these results expand the known functions of NEK2 and establish a resource for future studies aimed at dissecting how kinase interactomes are dynamically regulated across the cell cycle.

## 4 Methods

### 4.1 Cell culture and cell treatments

U2OS (ATCC, CRL3455), MDA-MB-231 (ATCC, HTB-26) and HEK293-T (ATCC, CRL-3216) cells were maintained in DMEM (Sigma, D6429) supplemented with 10% FBS (Biowest, S1520) and 1% penicillin-streptomycin at 37°C in 5% CO_2_ and were obtained from ATCC. All cell lines were tested monthly for mycoplasma contamination. TurboID and TurboID-NEK2 expressing U2OS cells were cultured in tetracycline-free FBS (Biowest, S181T) in routine culturing. 72 hours before performing proximity biotinylation experiments, medium was changed to dialyzed FBS (Biowest, S181D) to reduce any unwanted background biotinylation. Dox-inducible NEK2-WT expressing U2OS cells and NEK2-KO U2OS cells used in the study were generated and described previously^8,15^.

In targeted experiments, cells were treated with 2.5 μM MG132 (Sigma, M7449) for 1 or 2 hours and 10 μM cycloheximide (CHX; MedChem Express, HY-12320) for 2 to 24 hours.

### 4.2 Cloning and lentivirus generation

The TurboID-FLAG construct used in this study was a generous gift from Feng-Qian Li and Ken-Ichi Takemaru (Addgene, 124646). This plasmid was first linearized with the restriction enzymes BglII (NEB, R0144S) and XhoI (NEB, R0146S). NEK2-WT and NEK2-KD sequences^15^ were PCR-amplified from template DNA using Phusion high-fidelity DNA Polymerase (NEB, M0530S) with primers designed to include flanking BglII and XhoI sites. The PCR products were digested with the same enzymes and ligated into the prepared backbone using T4 DNA ligase (NEB, M0202S), generating TurboID–NEK2-WT and TurboID–NEK2-KD constructs. To generate doxycycline-inducible expression vectors, the pCW57-RFP-T2A-MCS plasmid (Addgene, 78933) was modified. The backbone was linearized with NheI (NEB, R3131S) and MluI (NEB, R3198S) to excise the RFP-T2A sequence. TurboID, TurboID–NEK2-WT, and TurboID–NEK2-KD coding sequences were PCR-amplified with primers introducing AvrII (NEB, R0174S) and MluI sites, since the NEK2 sequence contains an internal NheI site. These PCR products were digested with AvrII and MluI and cloned into the prepared backbone, yielding the pCW57-FLAG-TurboID and pCW57-FLAG-TurboID-NEK2 expression vectors.

For NUSAP1 cloning, the pLJC2-GPT2-3×FLAG plasmid (Addgene, 163447) was used as the starting backbone. The plasmid was digested with PacI (NEB, R0547S) and NotI (NEB, R3189S) to excise the GPT2 insert. The full-length NUSAP1 coding sequence was PCR-amplified from a U2OS cDNA library, digested with PacI and NotI, and ligated into the prepared backbone to generate the pLJC2-NUSAP1-3*×*FLAG construct.

The FLAG-tagged NEK2 overexpression system used for co-immunoprecipitation experiments was generated by cloning NEK2-WT and NEK2-KD coding sequences^15^ into the pCDH-EF1-FHC plasmid (Addgene, 64874). Inserts were PCR-amplified, digested with XbaI (NEB, R0145S) and NotI, and ligated into the backbone using T4 DNA ligase (NEB, M0202S). All constructs generated in this study were verified by Sanger sequencing prior to experimental use. The complete list of cloning primers is provided in Supplementary Table 1.

Lentiviral particles were produced in HEK293T cells using psPAX2 (Addgene, 12260) and VSV.G (Addgene, 14888) packaging plasmids. Target cells were infected at MOI 3 in the presence of 8 μg/mL polybrene and selected with 3 μg/mL puromycin (Sigma-Aldrich, P7255) for 3 days.

### 4.3 Cell cycle synchronization and flow cytometry

Cells were synchronized using a double thymidine block. Briefly, cells were treated with 2.5 mM thymidine (Sigma-Aldrich, T9250) for 18 hours, released by washing twice with PBS and once with full growth medium, and then cultured in fresh medium for 8 hours. A second block was applied with 2.5 mM thymidine for 18 hours. Cells were collected either immediately after the second block (0 h; G1/S sample), or released into fresh medium for 5 hours (late S sample) or 10 hours (G2/M sample). Time points were determined based on pilot experiments spanning 12 hours after release. For western blot validations, an M phase synchronization was also used. Cells were treated with 5 μM S-Trytyl-L-cysteine (STLC; MedChem Express, HY-W011102) for 16 hours, and mitotic cells were collected with a mitotic shake.

Cell cycle distribution was assessed by propidium iodide (PI; Sigma-Aldrich, P 4170) staining. Cell pellets were fixed with ethanol, treated with 50 μg/mL RNase A (Thermo, EN0531), stained with 100 μg/mL PI solution, washed with PBS, and analyzed on a flow cytometer (Cytoflex, Beckman Coulter). Gating strategy included FSC/SSC to select cells, SSC-A/SSC-H to exclude doublets, and PI intensity was plotted as histograms for the final gated populations. Unsynchronized populations were used as the control group in all flow cytometry experiments.

### 4.4 Biotinylation and Streptavidin pull-downs

Cells were maintained in dialyzed FBS for at least 72 hours before any biotinylation experiment. Expression of the constructs was induced by 1 μg/ml doxycycline, and biotinylation was performed with 500 μM biotin (Sigma Aldrich, B4639) for 30 minutes. To stop biotinylation, cell plates were washed three times with ice-cold PBS, with incubation on ice between washes. After washing, cells were trypsinized, and pellets were collected by centrifugation followed by two additional PBS washes.

Cells were lysed in RIPA buffer containing 0.1% NP-40 (Thermo, PI28324), 0.05% sodium deoxycholate (Thermo, 89904), 0.01% SDS (Bio Rad, 1610418), 15 mM NaCl (Isolab, 7647-14-5), 5 mM Tris-Cl-pH 8.0 (Thermo, 17926), protease inhibitor cocktail (Sigma-Aldrich, 11873580001), phosphatase inhibitor cocktail (Sigma-Aldrich, 4906845001), and PMSF (Sigma-Aldrich, P7626). Lysates were centrifuged at 15,000 *×* g for 15 min to remove debris, and protein concentrations were determined by BCA assay (Thermo, 10678484). Equal amounts of protein from each sample were incubated with streptavidin resin beads (Thermo, 88817) overnight at 4°C. Beads were subsequently washed four times with lysis buffer, and enriched proteins were processed for either western blotting or mass spectrometry analysis.

### 4.5 LC-MS/MS analysis

Proteins precipitated with streptavidin beads were subjected to on-bead tryptic digestion. Briefly, beads were washed with 8M urea solution, and proteins were reduced with 100 mM dithiothreitol (DTT) and alkylated with 100 mM iodoacetamide. Following alkylation, beads were washed with 50 mM ammonium bicarbonate and incubated with trypsin overnight at 37°C. The resulting peptide mixtures were collected and desalted using C18 STAGE tips.

Peptides were analyzed by C18 nanoflow reversed-phase liquid chromatography (nLC) using a Dionex Ultimate 3000 Nano Liquid Chromatography system (Thermo Scientific) coupled to a Q Exactive Orbitrap HF mass spectrometer (Thermo Scientific). Separation was performed on a 100 μm i.d. × 23 cm C18 column using an 80 min linear gradient of 5–25%, 25–40%, and 40–95% acetonitrile in 0.1% formic acid, at a flow rate of 300 nL/min with a total run time of 100 min. MS1 scans were acquired in the Orbitrap at a resolution of 70,000, with a mass range of 400–1500 m/z and maximum injection time of 32 ms. Data-dependent acquisition (DDA) was used, selecting the top 15 most intense ions for fragmentation. MS2 spectra were acquired by collision-induced dissociation (CID) at a resolution of 17,500, AGC target of 1e6, normalized collision energy of 26, and maximum injection time of 85 ms. The experimental setup and parameters were adapted from Ozkan et al^35^.

To analyze global proteomic changes associated with NEK2 deficiency, U2OS-WT cells and U2OS-NEK2-KO clone 2 cells were harvested by trypsinization, lysed in RIPA buffer as described in the Western blotting section, and subjected to LC–MS/MS analysis as outlined above. Proteomic analyses were performed using two independent biological replicates, each with three technical replicates, both for Turbo ID experiments and global proteomics.

### 4.6 Proteomics data analysis

Raw MS data were processed with MaxQuant^36^. Carbamidomethylation of cysteine was set as a fixed modification, while N-terminal acetylation and methionine oxidation were specified as variable modifications, with a maximum of five modifications allowed per peptide. Match-between-runs was enabled with a 20-minute matching time window. Label-free quantification (LFQ) was performed using classic normalization and a minimum LFQ ratio count of two, based on both unique and razor peptides. Trypsin was set as the protease, and searches were conducted against the human UniProt proteome FASTA file (UP000005640_9606). Intensity-based absolute quantification (iBAQ) was also enabled with loading normalization applied.

The resulting MaxQuant protein groups file was processed using the Cassiopeia R pipeline^37^. Contaminants were removed, and data processing included a razor + unique peptide filter (minimum two peptides), a valid values filter (>2 in at least one group), median normalization, and imputation using the “normal” mode (downshift = 1.8, width = 0.3). Enrichment logFC and p-values generated by Cassiopeia, which runs limma (v.3.50.1), were used for downstream analyses. Proteins were considered enriched in a given phase if logFC > 1 (two-fold enrichment compared with TurboID control). After excluding proteins below this threshold, p-values were used to define confident hits: proteins enriched in two phases were considered confident if p < 0.05 in both, while proteins enriched in all three phases were considered confident if p < 0.05 in all comparisons.

Global proteomics datasets were analyzed using the same MaxQuant–Cassiopeia pipeline with the same settings, ensuring consistency between TurboID-based interactome profiling and whole-proteome comparisons.

Both the raw MaxQuant output and Cassiopeia-processed results are provided as Source Data Tables 1 and 2, and the Cassiopeia pipeline output file has been deposited in Zenodo^23^.

**Table 2.**
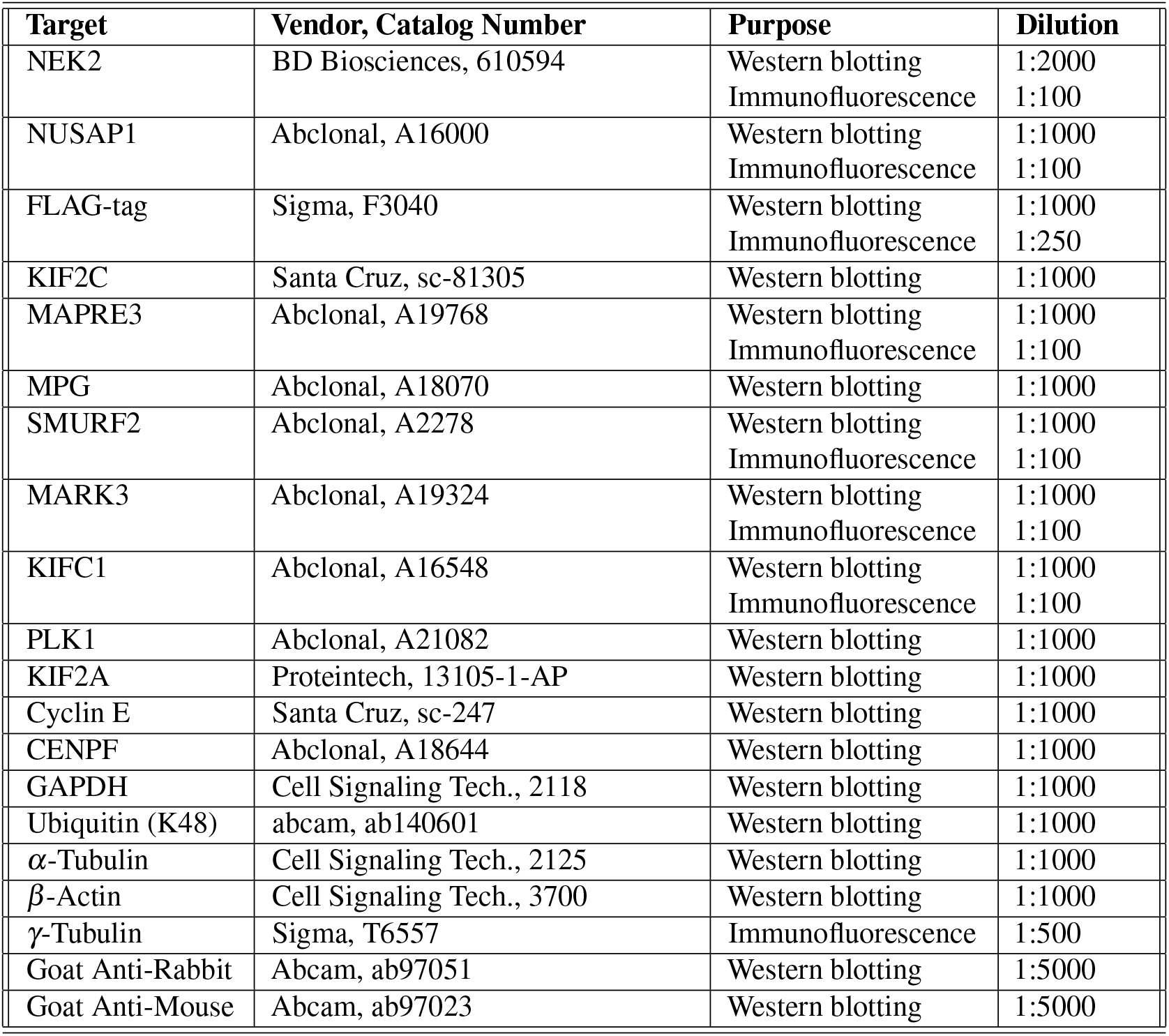
Supplementary Table - 2: Antibodies used in this study.

### 4.7 Gene set enrichment and over-representation analyses

Gene set enrichment analysis (GSEA) was performed using ranked protein lists ordered by enrichment scores using GSEA software (v 4.2.2). Gene Ontology (GO) enrichment analyses were conducted with clusterProfiler (v 4.17.0) and enrichplot (v 1.29.2) using the org.Hs.eg.db (v 3.21.0) annotation package in R (v 4.5.1).

### 4.8 k-means and generalized additive models (GAM) clustering analyses

For k-means clustering, we used phase-to-phase changes rather than raw logFC values. Specifically, we computed ΔlogFC(G1→S) = logFC_S_ −logFC_G1_ and ΔlogFC(S→G2) = logFC_G2_ − logFC_S_ for each protein, and applied k-means analysis (base R). The optimal cluster number was determined using the elbow and silhouette criteria and then fixed for downstream analyses.

To capture smoother dynamics across the G1–S–G2 transitions, generalized additive models (GAMs) were fitted to each protein with phase treated as a continuous variable with mgcv package (v 1.9.3) in R. For proteins with successful fits, model coefficients were extracted, scaled, and clustered with k-means, grouping proteins by the shape of their smoothed trajectories rather than absolute fold-change values. Predicted GAM values were then used to calculate phase-to-phase changes and generate scatter plots in transition space. Cluster assignments from both approaches were compared against original protein categories, and bar plots were generated to illustrate the composition of each cluster relative to prior classifications.

GO enrichment analysis was performed for each cluster, and results were visualized with dot plots, enrichment maps, and gene–concept networks, with the top enriched terms summarized per cluster.

### 4.9 Protein complex coordination analysis

NEK2-interacting proteins were mapped to known mammalian protein complexes using the CORUM database (v4.0)^27^. For each complex, we calculated coverage as the fraction of its total subunits detected in the NEK2 interactome. To assess whether complex subunits showed coordinated temporal interaction patterns with NEK2, we computed a pattern homogeneity score for each complex, defined as the proportion of its detected subunits that belonged to the same dominant GAM cluster (the most frequent cluster among the complex’s members). Complexes with homogeneity scores of 0.5 or higher were considered to exhibit coordinated interaction dynamics.

To address redundancy in complex definitions within the CORUM database, we implemented a similarity-based de-duplication step. Complexes were considered duplicates if their Jaccard similarity^38^ (the fraction of shared subunits relative to total unique subunits) exceeded 0.6. In such cases, only the complex with higher coverage in our dataset was retained. Complexes containing fewer than two interacting proteins were excluded from analysis. All analyses and visualizations were conducted in R (v4.2.0).

### 4.10 Direct ELISA assay for total biotinylated proteins

Enzyme-linked immunosorbent assay (direct ELISA) was performed following the protocol described by Lin (2015)^39^. Proteins were coated onto 96 well plates using carbonate–bicarbonate buffer and incubated overnight at 4°C, then denatured with 2% SDS for 30 minutes. Plates were blocked with SuperBlock Blocking Buffer (Thermo, 37515), incubated with Streptavidin–HRP (Thermo, 21140), and developed using an ECL substrate (Thermo, 32109). Signal intensity was quantified by measuring the luminescence with a plate reader (Gain: 150) (Synergy H1, Biotek).

### 4.11 Immunoprecipitation

Cells were collected by trypsinization (0.05%) and fixed with 1% paraformaldehyde (PFA; Merck, 104005.1000) for 7 min at room temperature. PFA was quenched with 1.25 mM ice-cold glycine (Isolab, 56-40-6), removed by centrifugation, and cell pellets were lysed in ice-cold IP Lysis Buffer (Thermo, 87787) supplemented with protease inhibitor (Roche, 11836170001), phosphatase inhibitor (Roche, 4906845001), and 1 mM PMSF for 45 min on ice. Protein concentrations were determined using the BCA assay (Thermo, 10678484).

For pre-clearance, lysates were incubated with Protein G magnetic beads (Thermo, 10004D) for 30 min at 4°C. Beads were magnetically removed, and the cleared lysates were incubated with either anti-FLAG antibody (Invitrogen, MA1-91878) or IgG control (Sigma, NI03) for 2 h at 4°C. Antibody–protein complexes were then captured by incubation with Protein G magnetic beads (Thermo, 88847) overnight at 4°C. Beads were washed four times with lysis buffer, and proteins were eluted by denaturation in 4*×* Laemmli buffer (Bio-Rad, 1610747) containing 10% DTT (Thermo, 20290) for 15 min at 95°C. Samples were subsequently analyzed by western blotting.

For ubiquitination assays, cells were treated with 5 μM MG132 5 hours prior to immunoprecipitation, as described before^40^.

### 4.12 Western blotting

Cells were lysed in RIPA buffer (0.1% NP-40, 0.05% sodium deoxycholate, 0.01% SDS, 15 mM NaCl, 5 mM Tris-Cl, pH 8.0) supplemented with protease inhibitors, phosphatase inhibitors, and PMSF, and incubated on ice for 15 min. Lysates were centrifuged at 15,000 × g for 15 min at 4°C to remove cell debris. Protein concentrations were determined using the Thermo BCA Protein Assay. Samples were denatured in 4 × Laemmli buffer containing 5% DTT for 12 min at 95°C and separated on 4–15% gradient SDS-PAGE gels. Proteins were transferred to PVDF membranes (Bio-Rad, 1620177), which were blocked in 5% non-fat dry milk (Bio-Rad, 1706404) in TBST for 1 h at room temperature. Membranes were incubated with primary antibodies (see Supplementary Table - 2) overnight at 4°C, followed by HRP-conjugated secondary antibodies (1:5000) for 1 h at room temperature. HRP-Streptavidin (Invitrogen, S32354, 1:5000) was used to detect biotinylation in western blots. Signals were visualized using Forte ECL reagent (Millipore, WBLUF0100) and imaged with an imaging device (Odyssey Fc, Licor).

### 4.13 Immunofluorescence microscopy

For immunofluorescence staining experiments, cells were grown on glass coverslips in 6-well plates. Following treatments, coverslips were fixed with methanol at -20°C for 5 minutes, washed three times with PBS, and blocked in blocking solution (5% BSA in 1 × PBS) for 1 hour at room temperature. After blocking, coverslips were incubated with primary antibodies overnight at 4°C, followed by secondary antibody incubation for 1 hour at room temperature. Slides were mounted with DAPI-containing mounting medium (Vectashield, H-1200-10) and imaged using a confocal microscope (DMI8, Leica). A fluorescence microscope (Axio Imager M1, Zeiss) was used for centrosome amplification counting experiments. For biotinylation stainings, Streptavidin-Alexa Fluor 488 (Invitrogen, S11223) was used. Antibodies used are listed in Supplementary Table-2.

### 4.14 RT-qPCR

RNA was extracted from cells using Nucleospin RNA (Macherey-Nagel; 740955) and cDNA was synthesized from 1 μg of total RNA using the M-MLV reverse transcriptase. SYBR green master mix (Roche; 04707516001) was used to amplify 10 ng of cDNA template. Real-time quantitative PCR was performed on a LightCycler 480 (Roche), and the relative fold change in gene expression was measured with the 2^-ΔΔCT^ method. Primers are listed in Supplementary Table-1.

### 4.15 NEK2 phosphorylation site analysis

Possible NEK2 phosphorylation site analysis was conducted using the PhosphoSitePlus database (https://www.phosphosite.org/). The kinase prediction tool was applied to identify NEK2 phosphorylation scores within NUSAP1 phosphorylation sites, and log_2_ scores were subsequently mapped and visualized along the NUSAP1 amino acid sequence.

### 4.16 Statistical analyses

Statistical analyses were performed using GraphPad Prism 9 and R (v 4.5.1). Proteomics data were analyzed with limma (v 3.50.1). For experiments requiring standard statistical approaches, one-way or two-way ANOVA tests were applied according to the experimental design, followed by appropriate multiple-comparisons & post hoc tests. A threshold of *p <* 0.05 was considered statistically significant.

## Supporting information

Uncut Western Blots

## 5 Acknowledgements

We are thankful to the KUTTAM-OMICS team for their technical assistance on LC-MS/MS analysis. The authors gratefully acknowledge the use of the services and facilities of the Koç University Research Center for Translational Medicine (KUTTAM), funded by the Presidency of Turkey, Presidency of Strategy and Budget.

## 7 Author contributions statement

Conceptualization: SCO and CA; Formal analysis: EC and SCO; Investigation: EC, SCO, BK, BMK and NEO; Methodology: SCO, EC; Visualization: SCO and EC; Project Administration: SCO, NO and CA; Funding Acquisition: SCO, CAA ; Writing - original draft: SCO, EC; Writing - review & editing: NO, CA.

## 8 Additional information

### Data availability

The mass spectrometry proteomics data have been deposited to the ProteomeXchange Consortium via the PRIDE partner repository^41^ with the dataset identifiers PXD068515 (TurboID experiments) and PXD046115 (Global proteomics). Raw and Cassiopeia processed proteomics results are shared in the Source data file, as well as in the Zenodo repository^23^.

## 9. Supplementary Tables

