## Supplemental Figures and Legends for "Proximity labeling reveals cell cycle–specific NEK2 interactions and a regulatory axis controlling NUSAP1 stability"

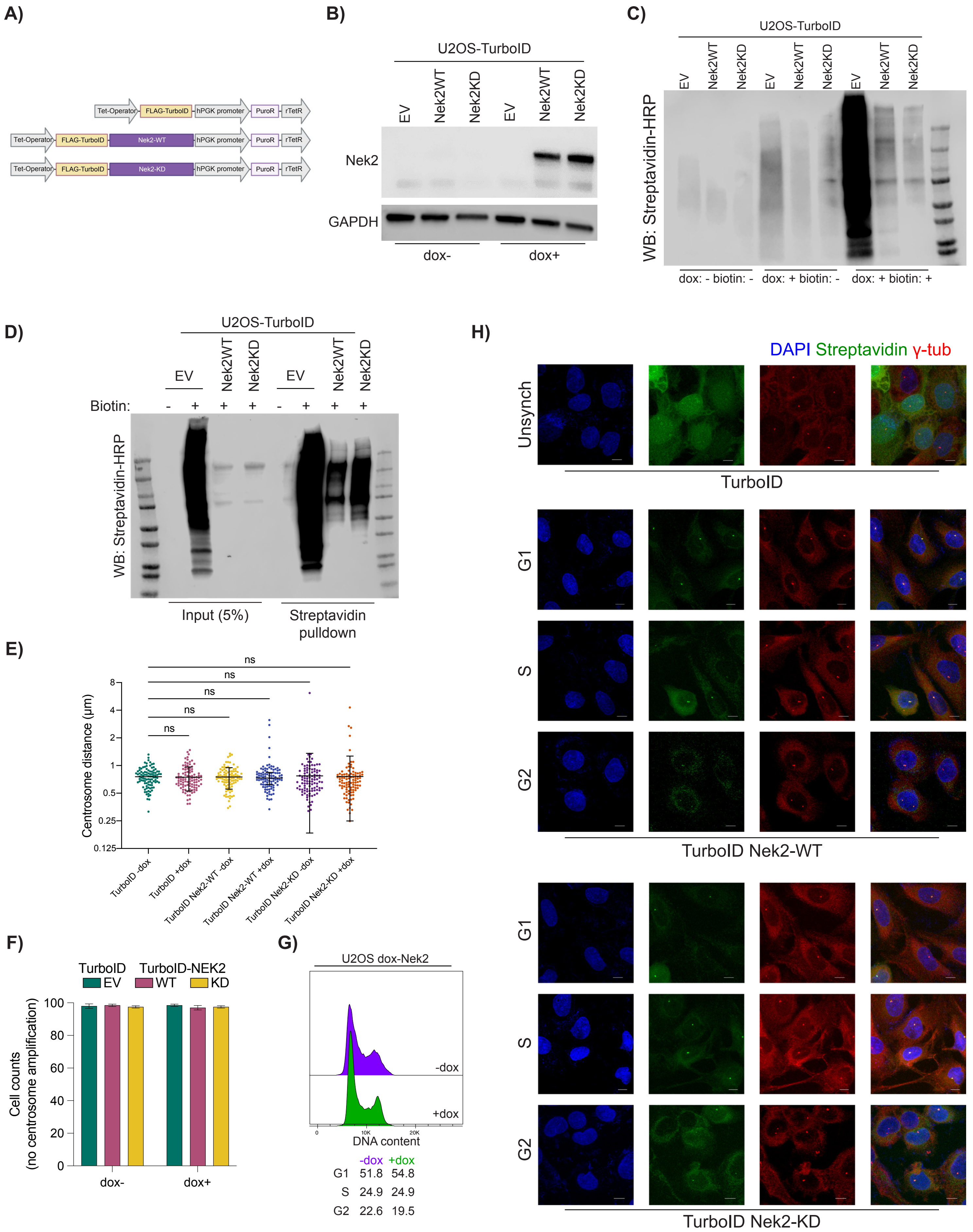
A)
Tet-Operator
FLAG-TurboID
hPGK promoter
PuroR
rTetR
Nek2-WT
Nek2-KD
B)
U2OS-TurboID
EV
Nek2WT
Nek2KD
Nek2
GAPDH
dox-
dox+
C)
U2OS-TurboID
WB: Streptavidin-HRP
dox: - biotin: -
dox: + biotin: -
dox: + biotin: +
D)
U2OS-TurboID
Biotin:
WB: Streptavidin-HRP
Input (5%)
Streptavidin pulldown
E)
ns
Centrosome distance (μm)
TurboID -dox
TurboID +dox
TurboID Nek2-WT -dox
TurboID Nek2-WT +dox
TurboID Nek2-KD -dox
TurboID Nek2-KD +dox
F)
TurboID
TurboID-NEK2
EV
WT
KD
Cell counts
(no centrosome amplification)
dox-
dox+
G)
U2OS dox-Nek2
-dox
+dox
DNA content
-dox +dox
G1 51.8 54.8
S 24.9 24.9
G2 22.6 19.5
H)
DAPI Streptavidin γ-tub
Unsynch
TurboID
G1
S
G2
TurboID Nek2-WT
TurboID Nek2-KD

**Supplementary Figure 1: Related to Figure 1.**

(A) Doxycycline (dox)-inducible FLAG-TurboID vectors used in this study. (B) Dox-induced expression of TurboID-NEK2-WT and TurboID-NEK2-KD constructs in U2OS cells. (C) Biotinylation patterns of TurboID groups in untreated (dox-), dox+biotin-, and dox+biotin+ conditions. (D) Enrichment of biotinylated proteins by streptavidin pulldown. (E–G) NEK2 overexpression in U2OS cells does not affect (E) interphase centrosome distances, (F) centrosome amplification, or (G) cell-cycle distribution. (H) Centrosomal biotinylation across cell-cycle phases in TurboID-NEK2-WT and TurboID-NEK2-KD cells.

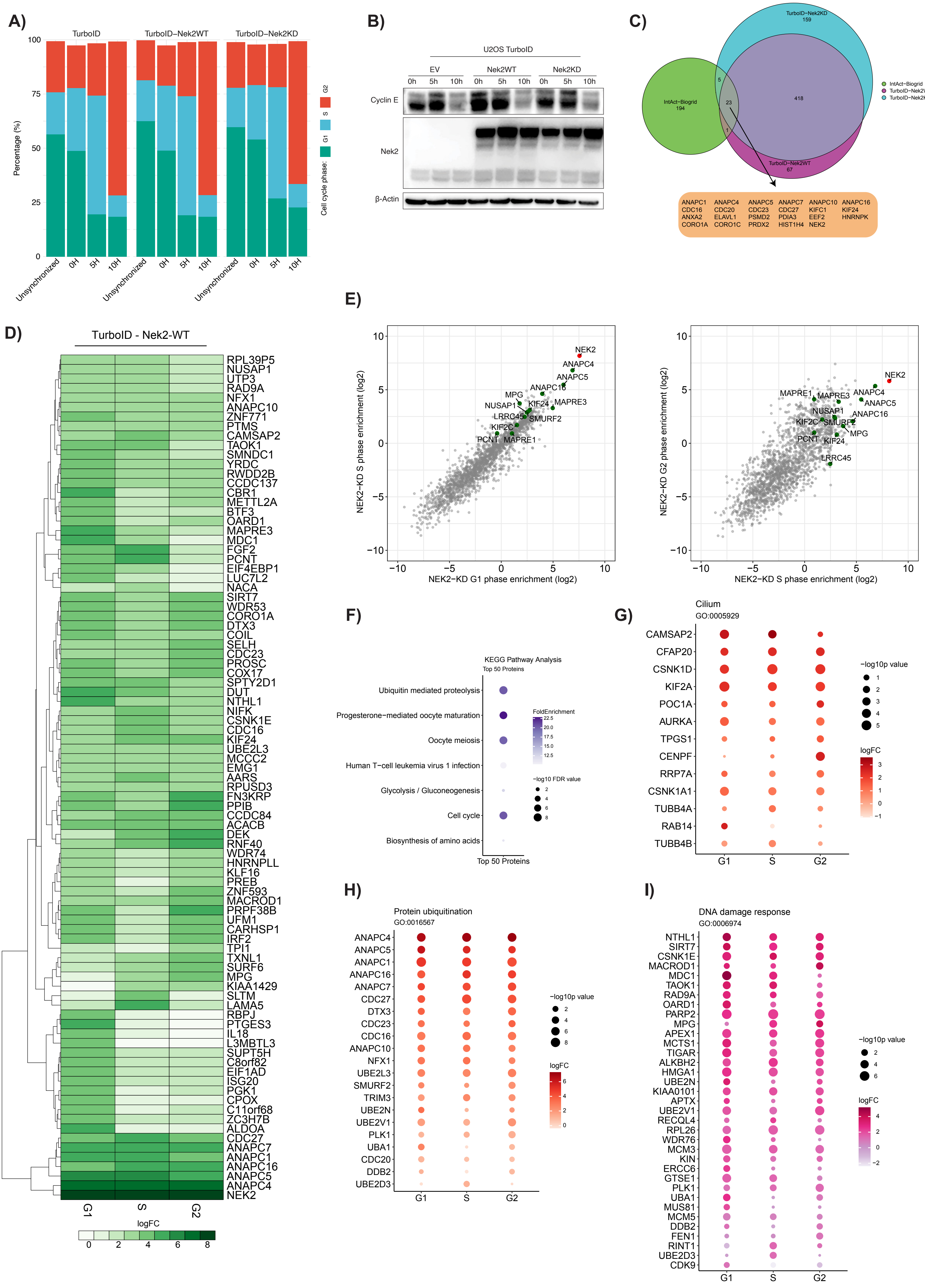
A)
TurboID
TurboID−Nek2WT
TurboID−Nek2KD
Percentage (%)
100
75
50
25
0
Unsynchronized
0H
5H
10H
Cell cycle phase:
G1
S
G2
B)
U2OS TurboID
EV
Nek2WT
Nek2KD
0h
5h
10h
Cyclin E
Nek2
β-Actin
C)
TurboID−Nek2KD
159
IntAct−Biogrid
194
TurboID−Nek2WT
67
5
23
418
1
IntAct−Biogrid
TurboID−Nek2WT
TurboID−Nek2KD
ANAPC1 ANAPC4 ANAPC5 ANAPC7 ANAPC10 ANAPC16
CDC16 CDC20 CDC23 CDC27 KIFC1 KIF24
ANXA2 ELAVL1 PSMD2 PDIA3 EEF2 HNRNPK
CORO1A CORO1C PRDX2 HIST1H4 NEK2
D)
TurboID - Nek2-WT
RPL39P5
NUSAP1
UTP3
RAD9A
NFX1
ANAPC10
ZNF771
PTMS
CAMSAP2
TAOK1
SMNDC1
YRDC
RWDD2B
CCDC137
CBR1
METTL2A
BTF3
OARD1
MAPRE3
MDC1
FGF2
PCNT
EIF4EBP1
LUC7L2
NACA
SIRT7
WDR53
CORO1A
DTX3
COIL
SELH
CDC23
PROSC
COX17
SPTY2D1
DUT
NTHL1
NIFK
CSNK1E
CDC16
KIF24
UBE2L3
MCCC2
EMG1
AARS
RPUSD3
FN3KRP
PPIB
CCDC84
ACACB
DEK
RNF40
WDR74
HNRNPLL
KLF16
PREB
ZNF593
MACROD1
PRPF38B
UFM1
CARHSP1
IRF2
TPI1
TXNL1
SURF6
MPG
KIAA1429
SLTM
LAMA5
RBPJ
PTGES3
IL18
L3MBTL3
SUPT5H
C8orf82
EIF1AD
ISG20
PGK1
CPOX
C11orf68
ZC3H7B
ALDOA
CDC27
ANAPC7
ANAPC1
ANAPC16
ANAPC5
ANAPC4
NEK2
G1
S
G2
logFC
0
2
4
6
8
E)
NEK2−KD S phase enrichment (log2)
NEK2−KD G1 phase enrichment (log2)
NEK2
ANAPC4
ANAPC5
ANAPC16
MPG
NUSAP1
KIF24
MAPRE3
LRRC45
SMURF2
KIF2C
PCNT
MAPRE1
NEK2−KD G2 phase enrichment (log2)
NEK2−KD S phase enrichment (log2)
NEK2
MAPRE1
MAPRE3
ANAPC4
ANAPC5
NUSAP1
ANAPC16
KIF2C
SMURF2
MPG
PCNT
KIF24
LRRC45
10
5
0
−5
−10
F)
KEGG Pathway Analysis
Top 50 Proteins
Ubiquitin mediated proteolysis
Progesterone−mediated oocyte maturation
Oocyte meiosis
Human T−cell leukemia virus 1 infection
Glycolysis / Gluconeogenesis
Cell cycle
Biosynthesis of amino acids
Top 50 Proteins
FoldEnrichment
22.5
20.0
17.5
15.0
12.5
−log10 FDR value
2
4
6
8
G)
Cilium
GO:0005929
CAMSAP2
CFAP20
CSNK1D
KIF2A
POC1A
AURKA
TPGS1
CENPF
RRP7A
CSNK1A1
TUBB4A
RAB14
TUBB4B
G1
S
G2
−log10p value
1
2
3
4
5
logFC
3
2
1
0
−1
H)
Protein ubiquitination
GO:0016567
ANAPC4
ANAPC5
ANAPC1
ANAPC16
ANAPC7
CDC27
DTX3
CDC23
CDC16
ANAPC10
NFX1
UBE2L3
SMURF2
TRIM3
UBE2N
UBE2V1
PLK1
UBA1
CDC20
DDB2
UBE2D3
G1
S
G2
−log10p value
2
4
6
8
logFC
6
4
2
0
I)
DNA damage response
GO:0006974
NTHL1
SIRT7
CSNK1E
MACROD1
MDC1
TAOK1
RAD9A
OARD1
PARP2
MPG
APEX1
MCTS1
TIGAR
ALKBH2
HMGA1
UBE2N
KIAA0101
APTX
UBE2V1
RECQL4
RPL26
WDR76
MCM3
KIN
ERCC6
GTSE1
PLK1
UBA1
MUS81
MCM5
DDB2
FEN1
RINT1
UBE2D3
CDK9
G1
S
G2
−log10p value
2
4
6
logFC
4
2
0
−2

**Supplementary Figure 2: Related to Figure 2.** (A) Cell cycle distribution of synchronized proximity interaction experiments. (B) Western blot showing reduction of Cyclin E during progression into the G2/M phase. (C) Venn diagram illustrating the overlap between NEK2-WT and NEK2-KD hits with previously reported NEK2 interactors. (D) Heatmap of enrichment logFC values for the top 50 NEK2-WT interactome hits across cell cycle phases. (E) Distribution of NEK2-interacting proteins detected in NEK2-KD proximity labeling experiments. Left: G1–S comparison; Right: S–G2 comparison. (F) KEGG pathway analysis of the top 50 interacting proteins. (G–I) Enrichment of proteins in the NEK2 interactome across selected GO terms in TurboID–NEK2-WT samples at different cell cycle phases: (G) cilium, (H) protein ubiquitination, and (I) DNA damage response.

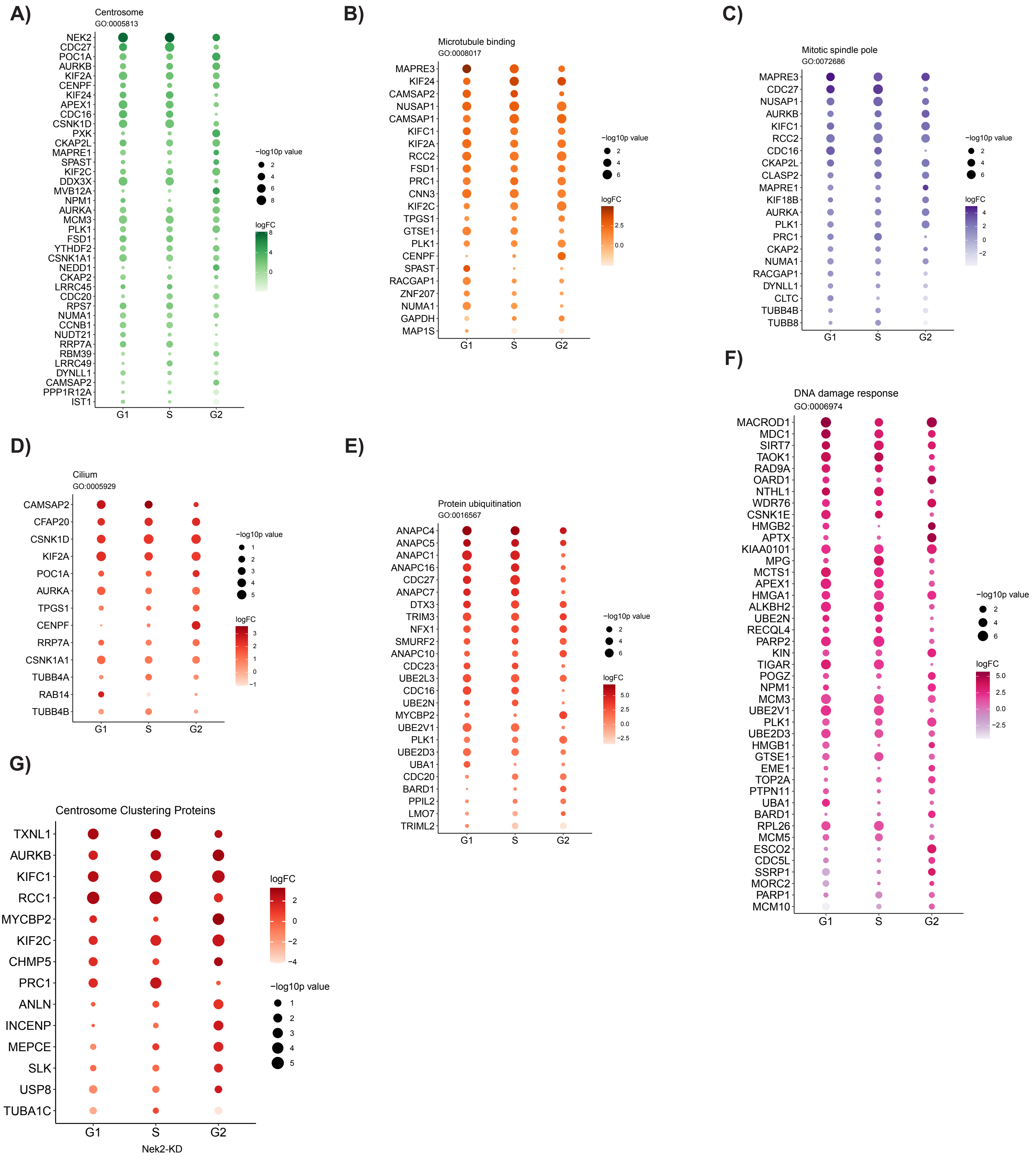


**Supplementary Figure 3: Related to Figure 2.** Enrichment of proteins in the NEK2 interactome across selected GO terms in TurboID–NEK2-KD samples at different cell cycle phases. (A) Centrosome, (B) Microtubule-binding proteins, (C) Mitotic spindle pole–associated proteins, (D) Cilium, (E) Protein ubiquitination, (F) DNA damage response, and (G) Centrosome clustering proteins.

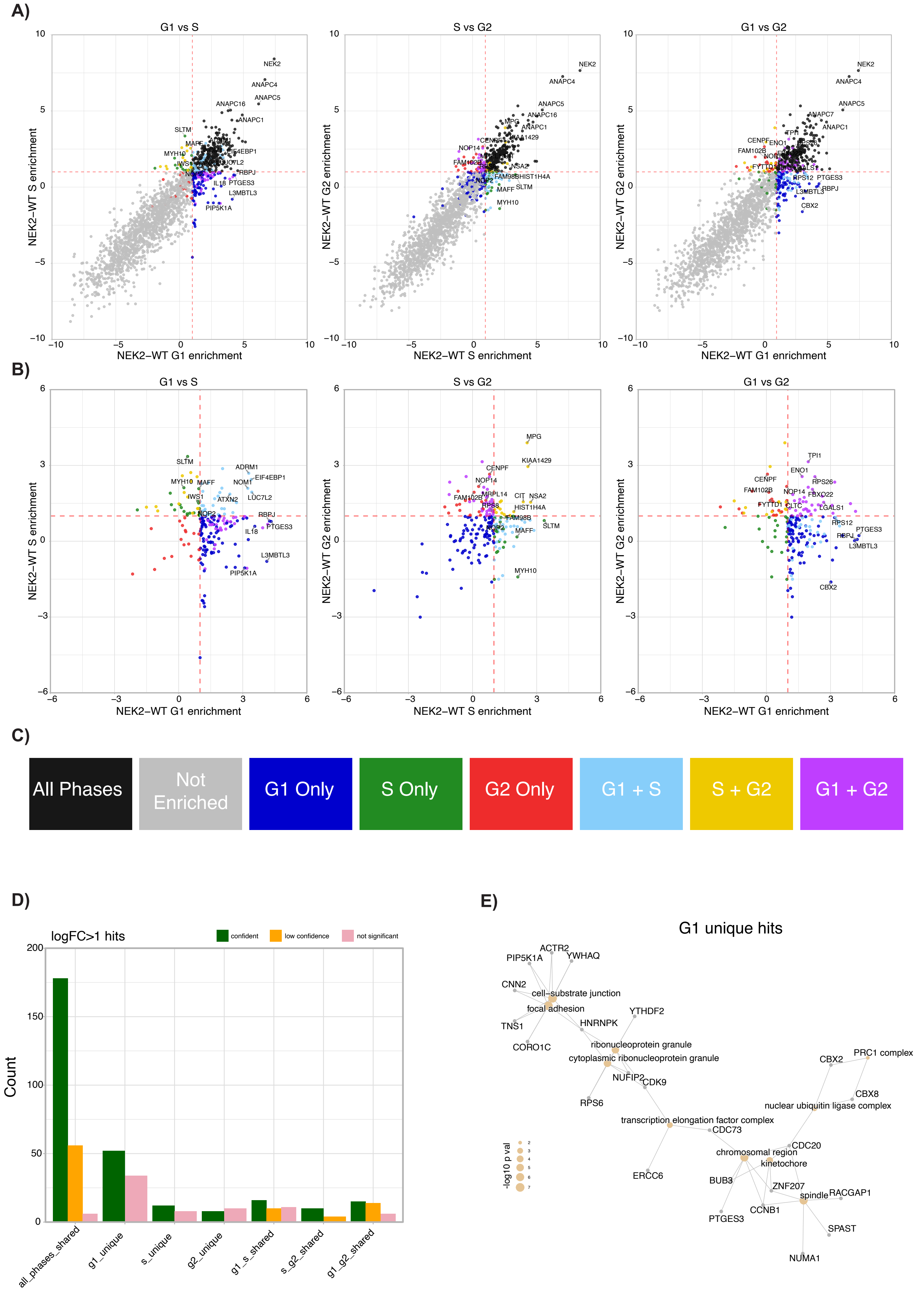
A)
G1 vs S
S vs G2
G1 vs G2
NEK2–WT S enrichment
NEK2–WT G2 enrichment
NEK2–WT G1 enrichment
NEK2–WT S enrichment
NEK2–WT G1 enrichment
NEK2
ANAPC4
ANAPC5
ANAPC16
ANAPC1
ANAPC7
SLTM
MAFF
MYH10
EIF4EBP1
LUC7L2
RBPJ
IL18
PTGES3
L3MBTL3
PIP5K1A
MPG
KIAA1429
CENPF
NOP14
FAM102B
NSA2
FAM98B
HIST1H4A
MYH10
TPI1
ENO1
RPS12
CBX2
B)
G1 vs S
S vs G2
G1 vs G2
SLTM
ADRM1
EIF4EBP1
NOM1
MYH10
MAFF
IWS1
ATXN2
LUC7L2
NOP2
RBPJ
IL18
PTGES3
L3MBTL3
PIP5K1A
MPG
KIAA1429
CENPF
NOP14
MRPL14
FAM102B
RPS8
CIT
NSA2
HIST1H4A
FAM98B
MAFF
SLTM
MYH10
TPI1
ENO1
RPS26
FBXO22
FYTTD1
CLTC
LGALS1
RPS12
RBPJ
PTGES3
L3MBTL3
CBX2
C)
All Phases
Not Enriched
G1 Only
S Only
G2 Only
G1 + S
S + G2
G1 + G2
D)
logFC>1 hits
confident
low confidence
not significant
Count
all_phases_shared
g1_unique
s_unique
g2_unique
g1_s_shared
s_g2_shared
g1_g2_shared
E)
G1 unique hits
ACTR2
PIP5K1A
YWHAQ
CNN2
cell−substrate junction
focal adhesion
YTHDF2
TNS1
HNRNPK
CORO1C
ribonucleoprotein granule
cytoplasmic ribonucleoprotein granule
NUFIP2
CDK9
RPS6
transcription elongation factor complex
CDC73
ERCC6
CBX2
PRC1 complex
CBX8
nuclear ubiquitin ligase complex
CDC20
chromosomal region
kinetochore
BUB3
ZNF207
spindle
RACGAP1
CCNB1
PTGES3
SPAST
NUMA1
-log10 p val

**Supplementary Figure 4: Related to Figure 3.** (A) All proteins enriched in the NEK2 interactome, shown with distinct enrichment patterns in comparative plots. Left: G1–S comparison; Middle: S–G2 comparison; Right: G1–G2 comparison. (B) Enriched protein distributions after excluding non-enriched proteins and proteins detected in all phases. (C) Color legend used in scatter plots in (A) and (B). (D) Counts of confident, low-confidence, and non-significant enriched hits across categories. (E) Gene–concept networks of G1-unique hits.

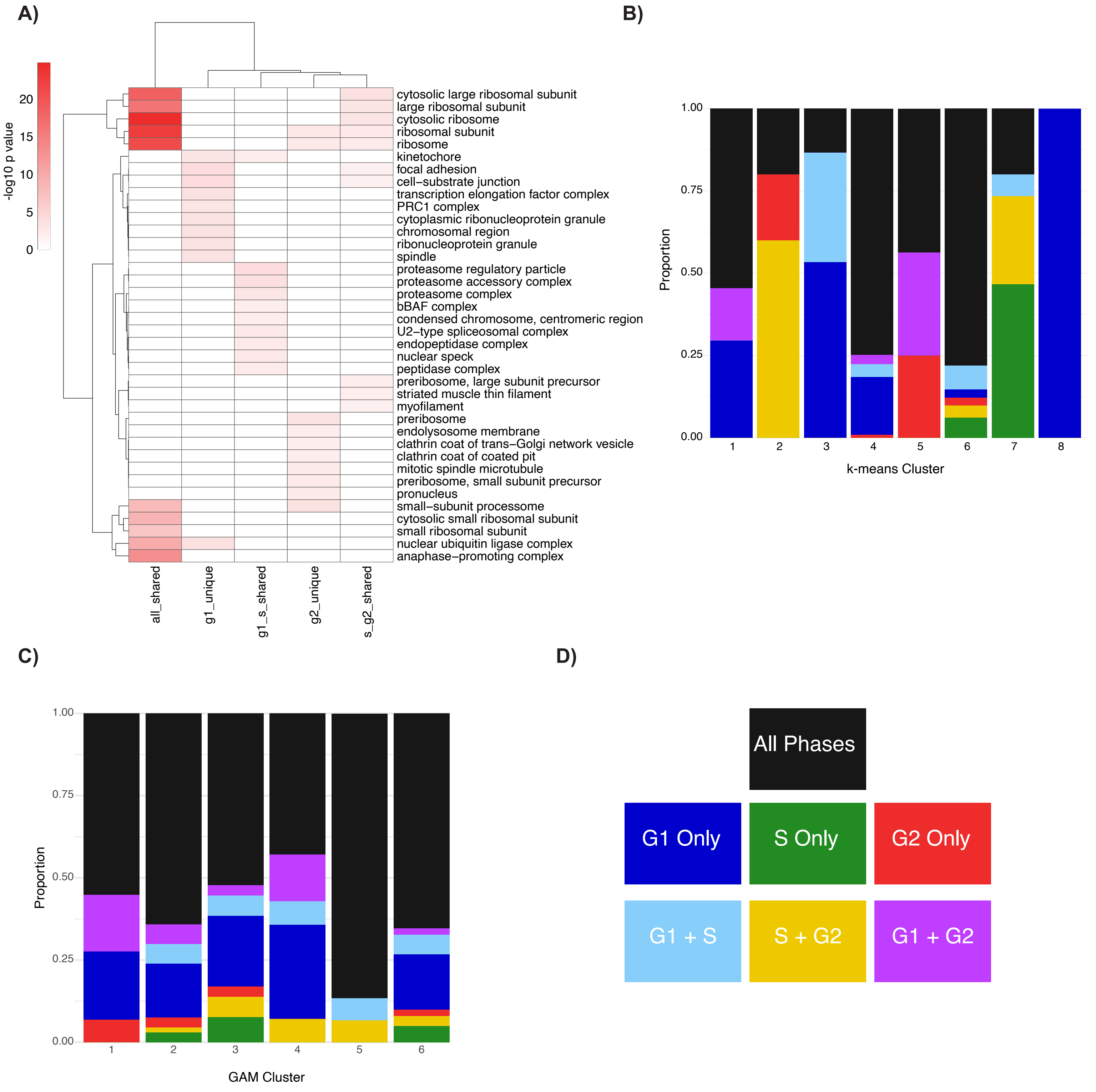


**Supplementary Figure 5: Related to Figure 3.** (A) Heatmap of significant GO-CC terms across categories of NEK2 interactome–enriched proteins. (B) Distribution of cell cycle–dependent enrichment categories across k-means clusters. (C) Distribution of cell cycle–dependent enrichment categories across GAM clusters. (D) Color legend used in the bar plots shown in (B) and (C).

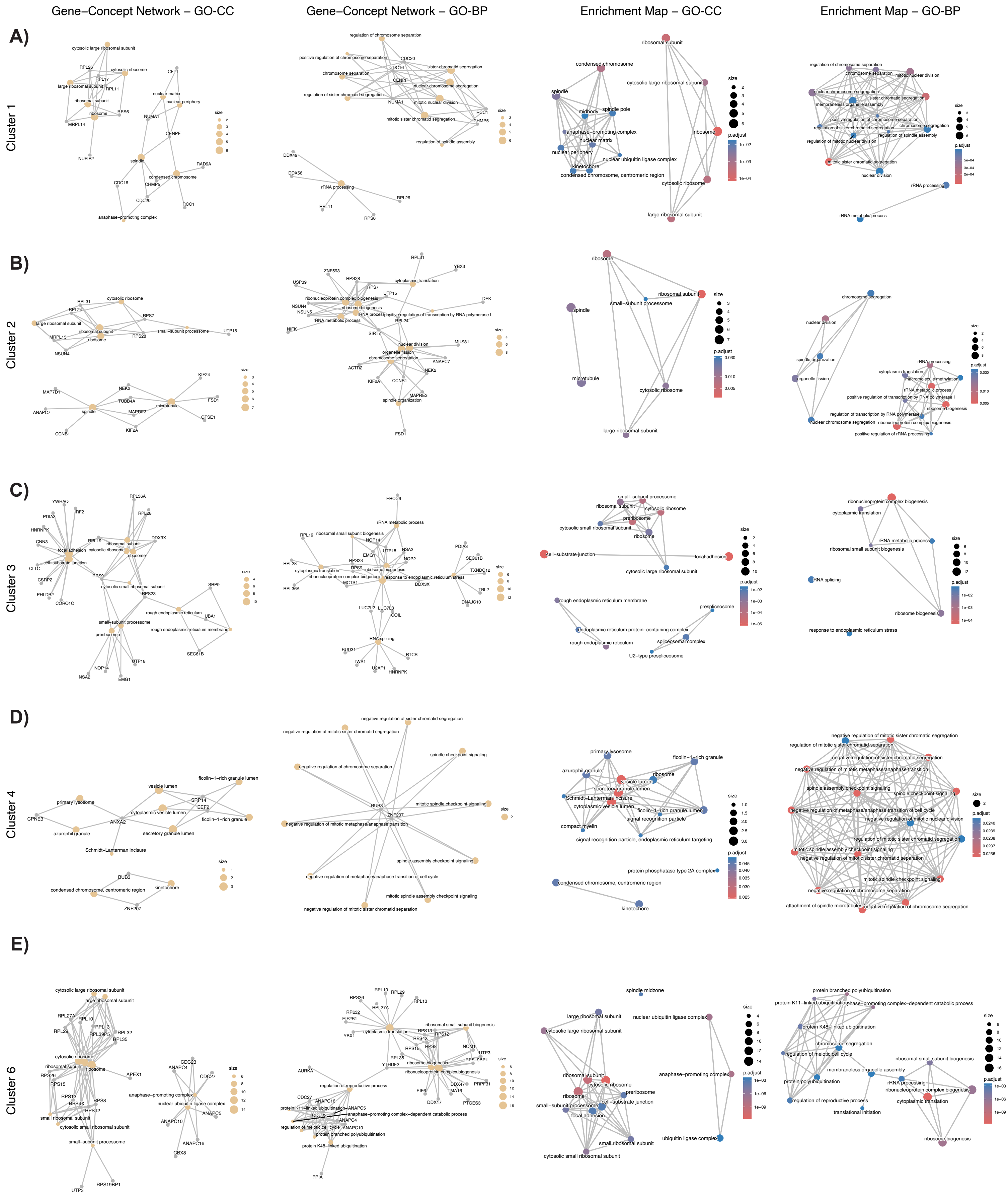

Gene−Concept Network − GO-CC
Gene−Concept Network − GO-BP
Enrichment Map − GO-CC
Enrichment Map − GO-BP
A)
Cluster 1
B)
Cluster 2
C)
Cluster 3
D)
Cluster 4
E)
Cluster 6

**Supplementary Figure 6: Related to Figure 3.** GO-CC and GO-BP enrichment results of different GAM clusters. (A–E) correspond to clusters 1, 2, 3, 4, and 6, respectively. For each cluster: Left, gene–concept network of GO-CC terms; mid-left, gene–concept network of GO-BP terms; mid-right, enrichment map of GO-CC terms; right, enrichment map of GO-BP terms.

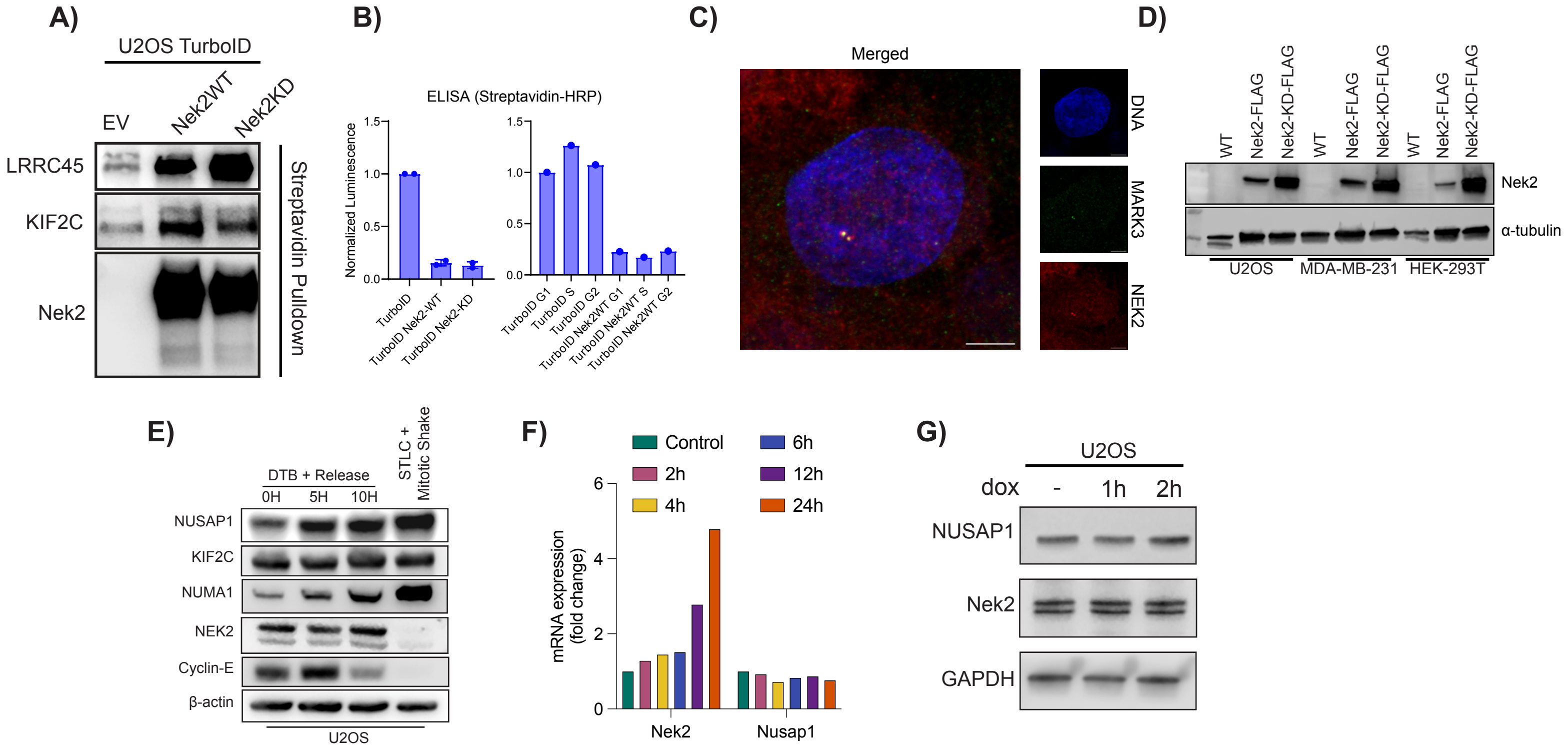


**Supplementary Figure 7: Related to Figure 4 and 5.** (A) Enrichment of LRRC45 and KIF2C in TurboID–NEK2-WT and TurboID–NEK2-KD expressing cells. (B) ELISA results from unsynchronized and synchronized cell lines used for validation western blot experiments (C) Co-localization of MARK3 and NEK2 at centrosomes. (D) Generation of FLAG-tagged NEK2 overexpressing cell lines for co-IP experiments. (E) Cell cycle dependent regulation of endogenous NUSAP1, KIF2C, NUMA1, and NEK2 protein levels. (F) NEK2 and NUSAP1 mRNA expression levels in U2OS-dox-NEK2 cells at indicated time points following doxycycline induction. (G) NUSAP1 and NEK2 levels in dox treated U2OS cells.

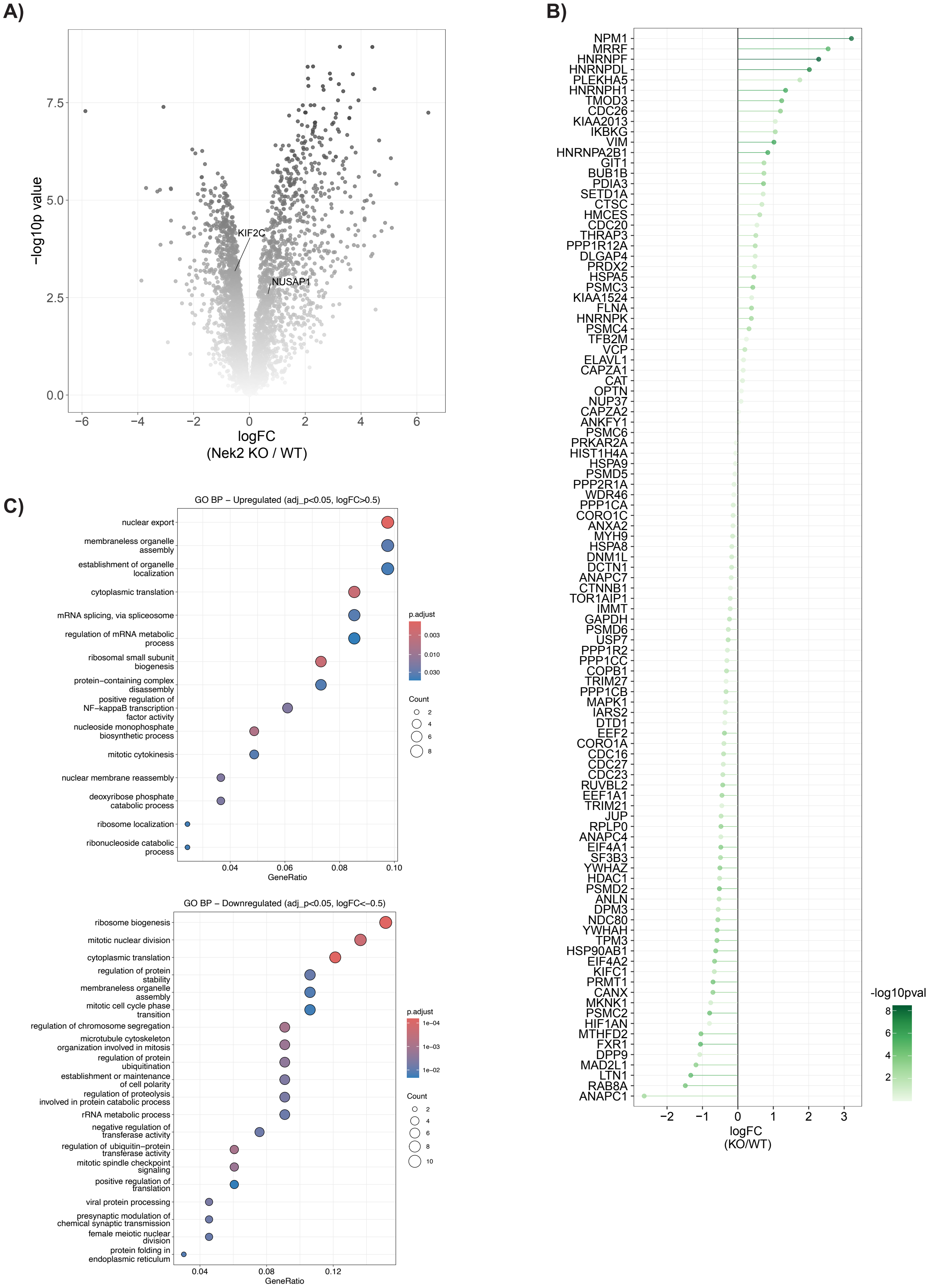


**Supplementary Figure 7: Related to Figure 6.** (A) Global proteomic changes in NEK2-KO cells compared with U2OS-WT cells. (B) Effect of NEK2 knockout on the expression of previously reported NEK2 interaction partners. (C) GO-BP enrichment analysis of up- and down-regulated NEK2-interacting proteins identified in the NEK2 knockout global proteomics dataset.
