## Supplementary material for "Proximity labeling reveals cell cycle–specific NEK2 interactions and a regulatory axis controlling NUSAP1 stability": Uncut Western Blots

1C)

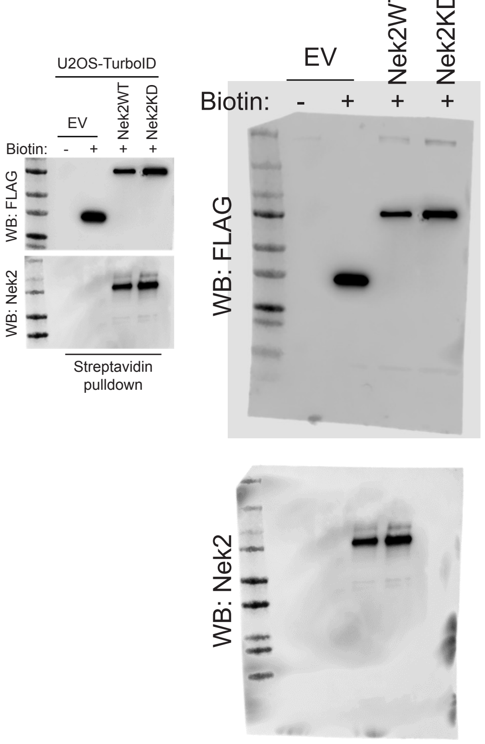


S1B)

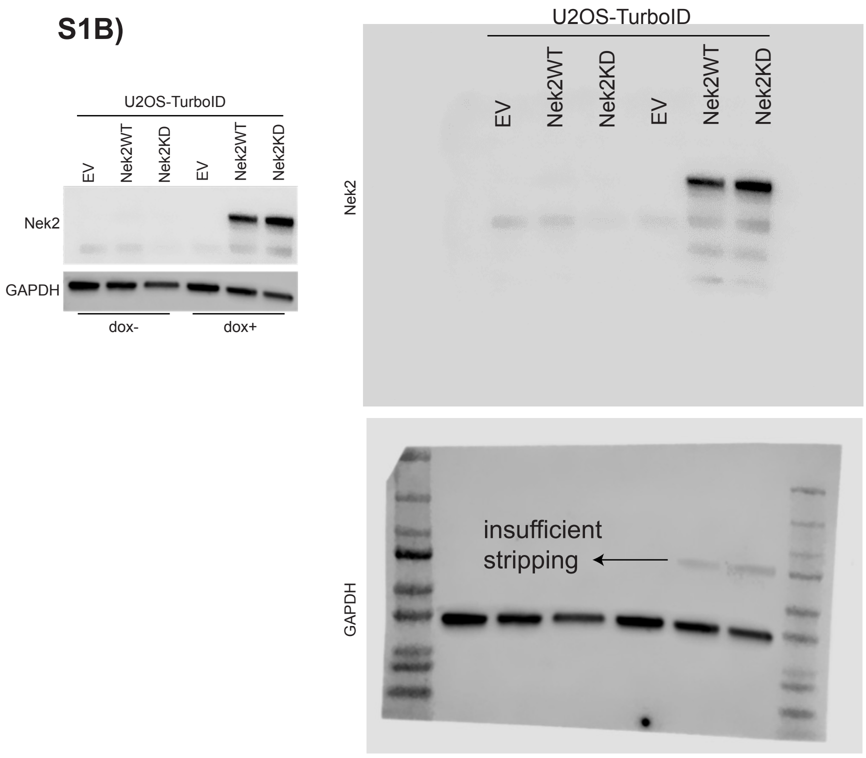


S1C

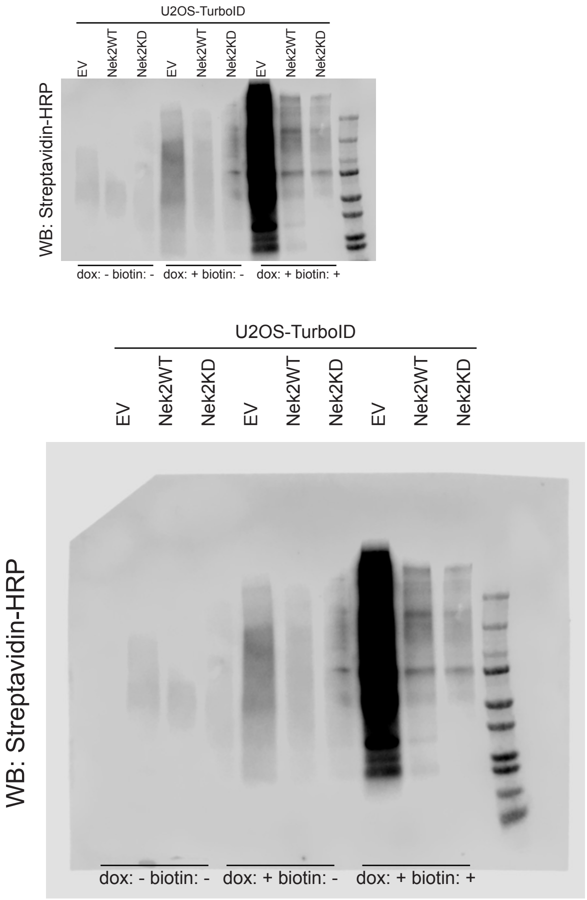


S1D)

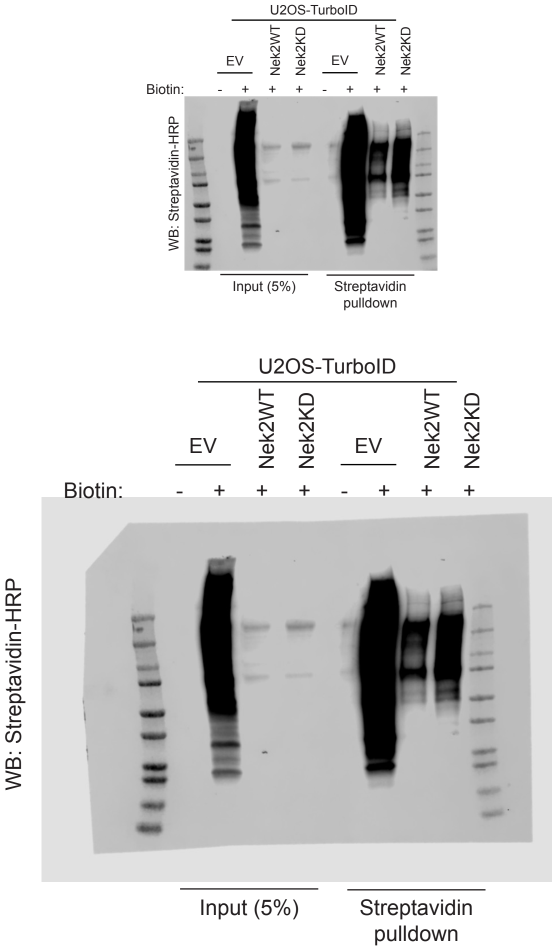

**S2B)**

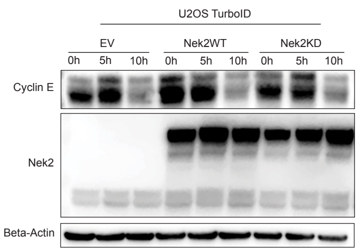


Cyclin E

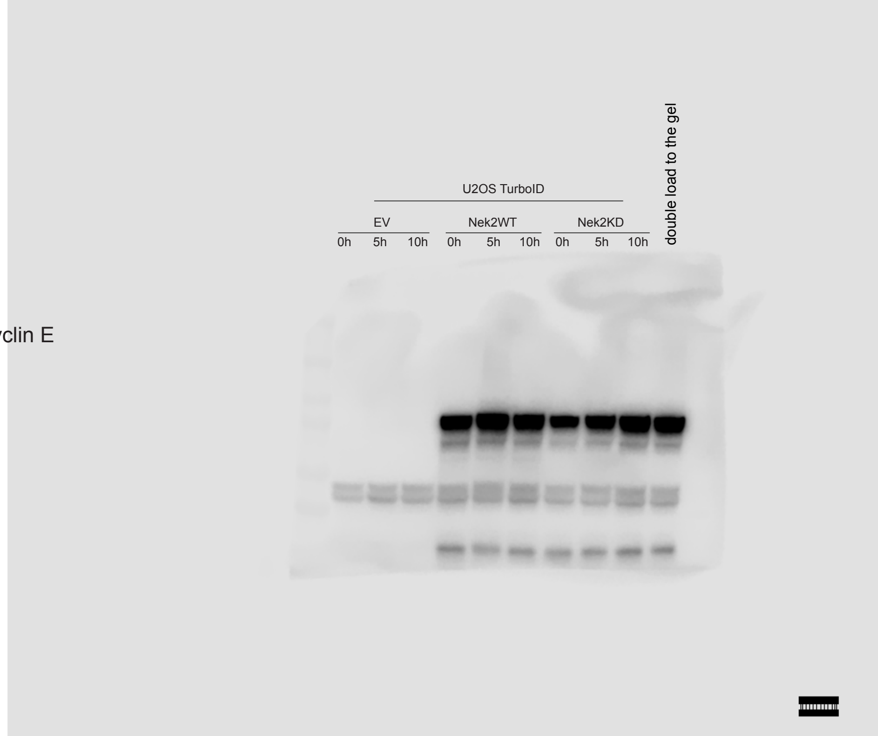


Nek2

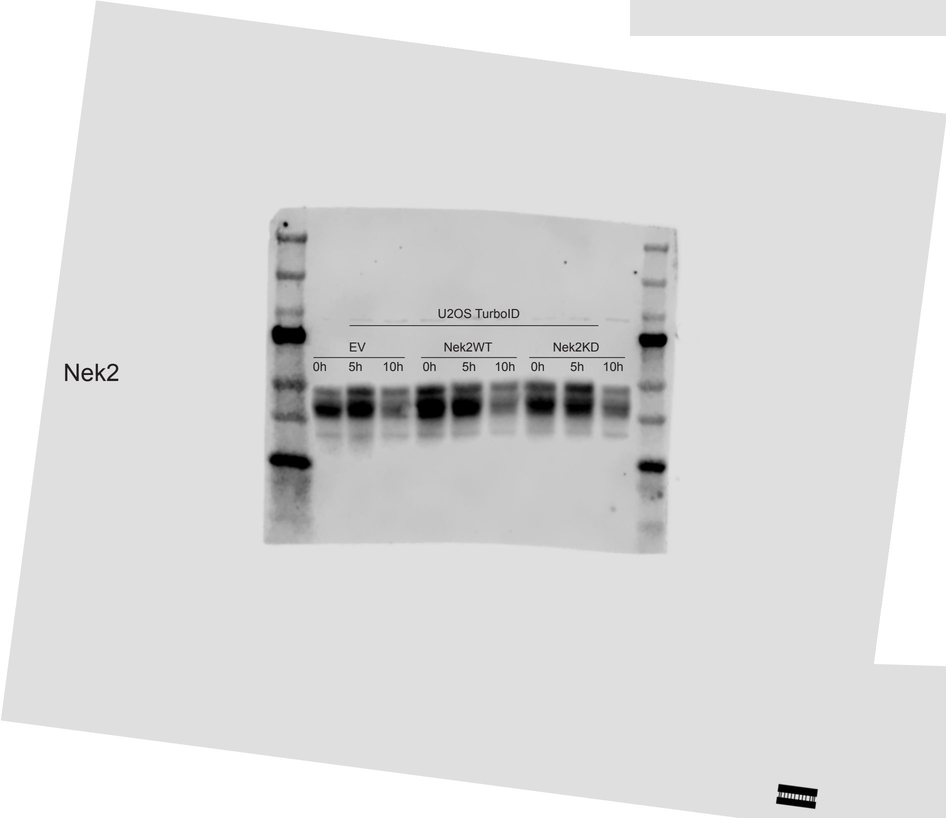


Beta-Actin

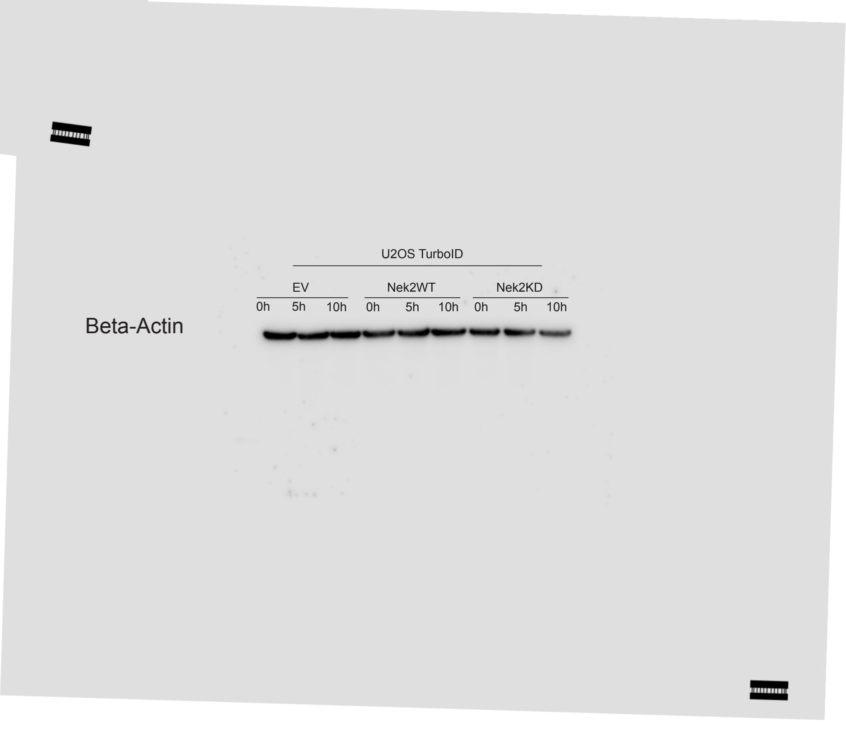

4A

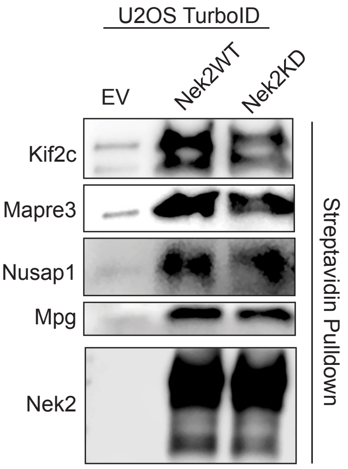
U2OS TurboID
EV
Nek2WT
Nek2KD
Kif2c
Mapre3
Nusap1
Mpg
Nek2
Streptavidin Pulldown

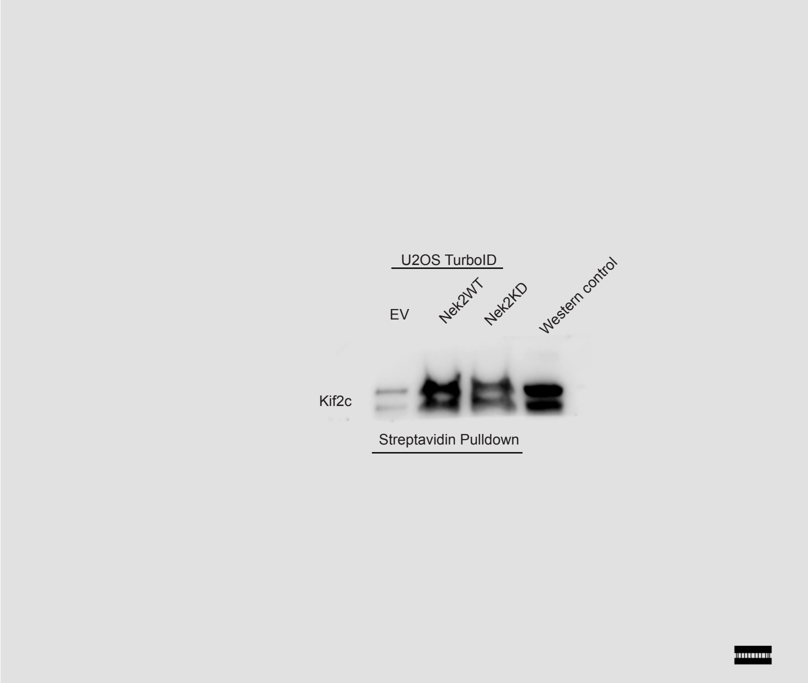
U2OS TurboID
EV
Nek2WT
Nek2KD
Western control
Kif2c
Streptavidin Pulldown

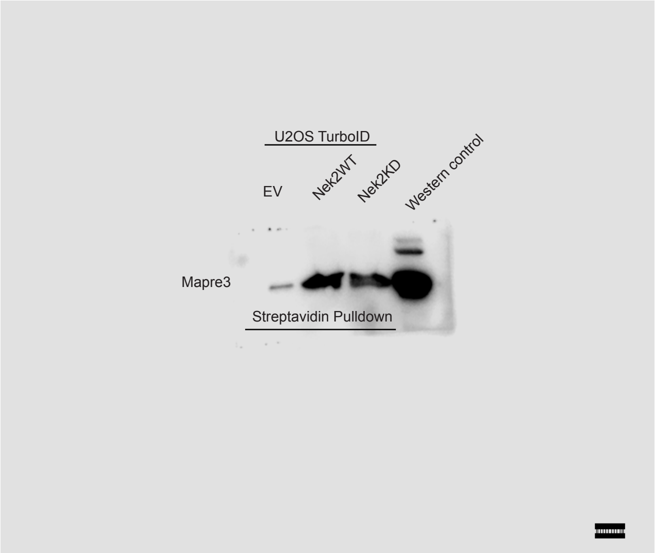
U2OS TurboID
EV
Nek2WT
Nek2KD
Western control
Mapre3
Streptavidin Pulldown

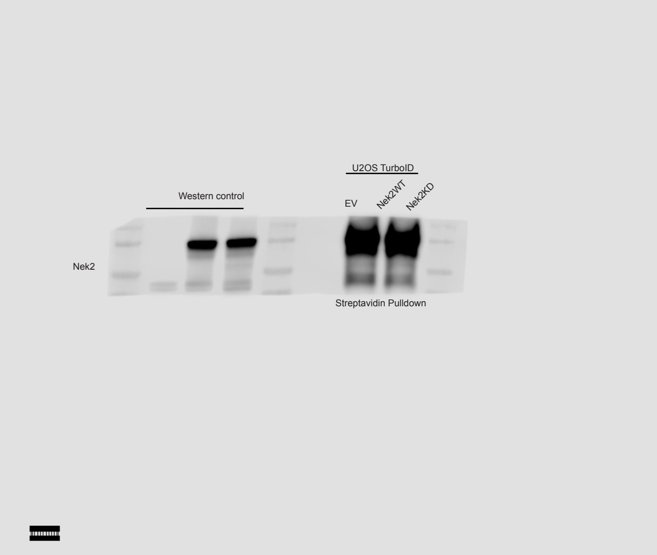
Western control
U2OS TurboID
EV
Nek2WT
Nek2KD
Nek2
Streptavidin Pulldown

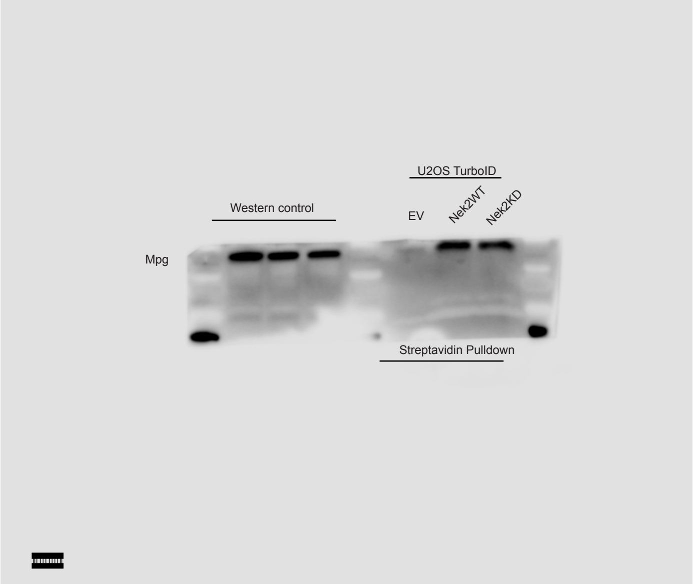
Western control
U2OS TurboID
EV
Nek2WT
Nek2KD
Mpg
Streptavidin Pulldown

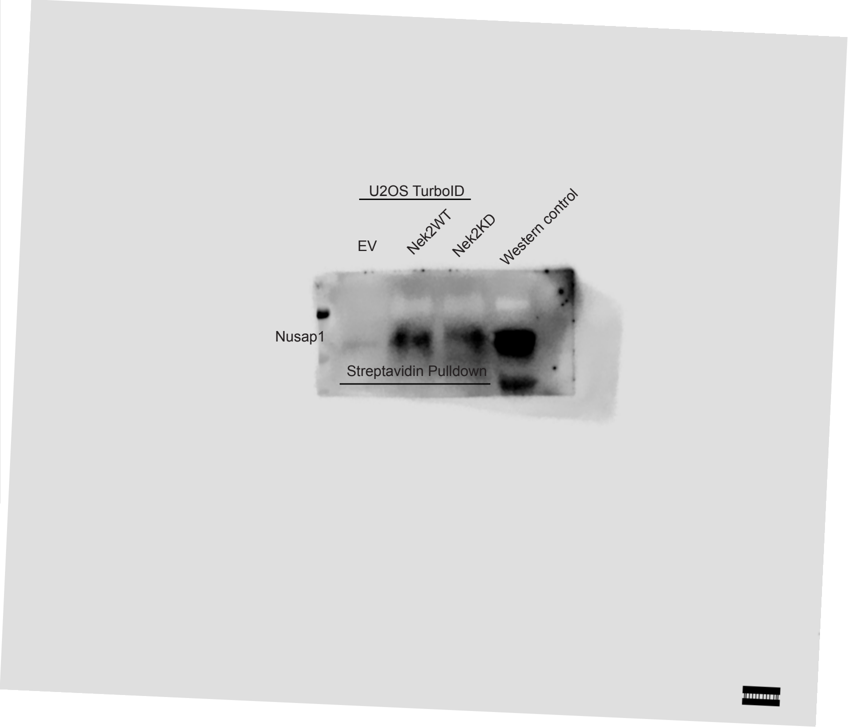
U2OS TurboID
EV
Nek2WT
Nek2KD
Western control
Nusap1
Streptavidin Pulldown

**4B**

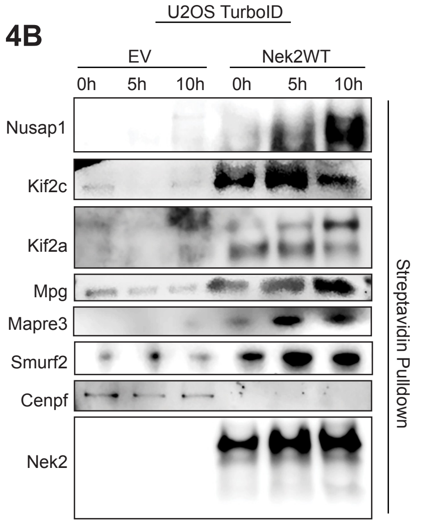
U2OS TurboID
EV
Nek2WT
0h
5h
10h
0h
5h
10h
Nusap1
Kif2c
Kif2a
Mpg
Mapre3
Smurf2
Cenpf
Nek2
Streptavidin Pulldown

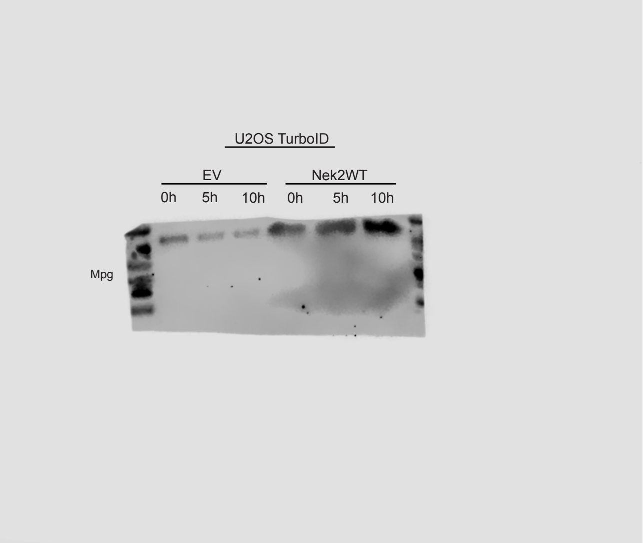
U2OS TurboID
EV
Nek2WT
0h
5h
10h
0h
5h
10h
Mpg

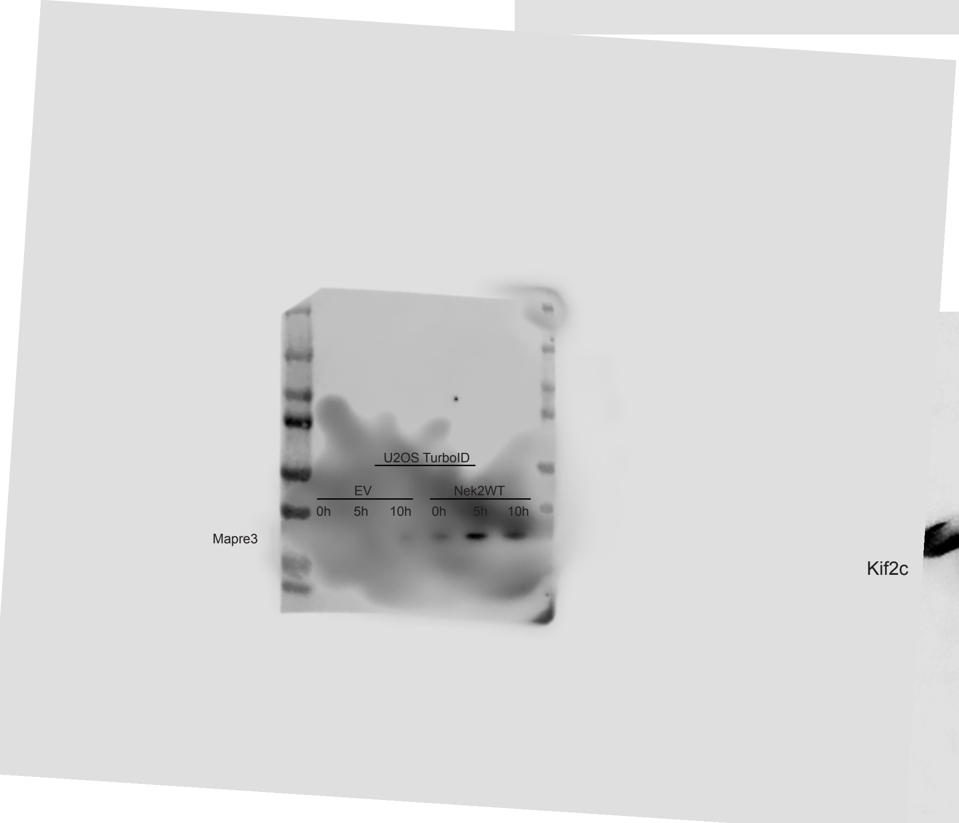
U2OS TurboID
EV
Nek2WT
0h
5h
10h
0h
5h
10h
Mapre3

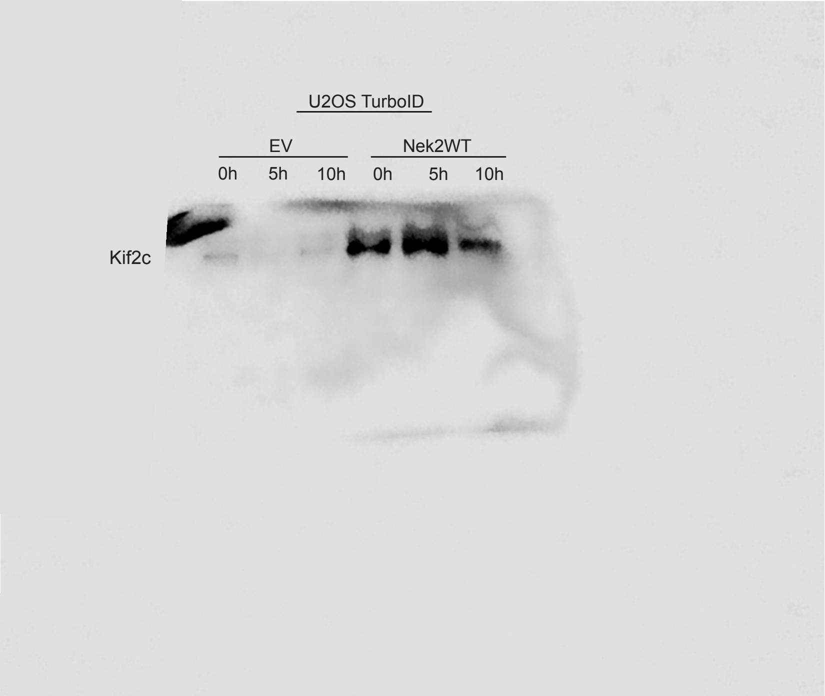
U2OS TurboID
EV
Nek2WT
0h
5h
10h
0h
5h
10h
Kif2c

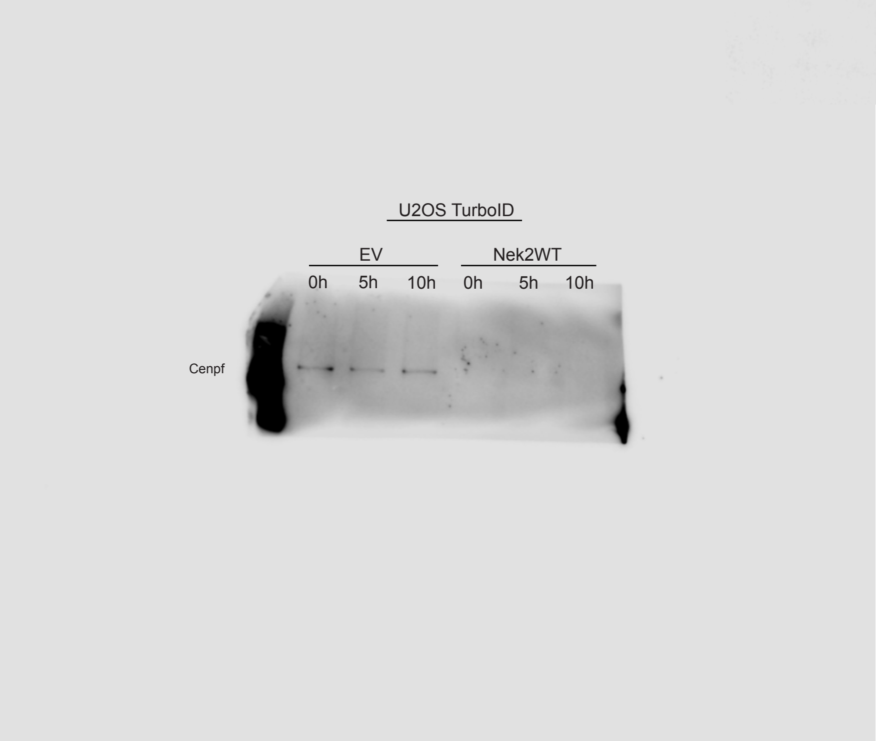
U2OS TurboID
EV
Nek2WT
0h
5h
10h
0h
5h
10h
Cenpf

4B)
(continued)

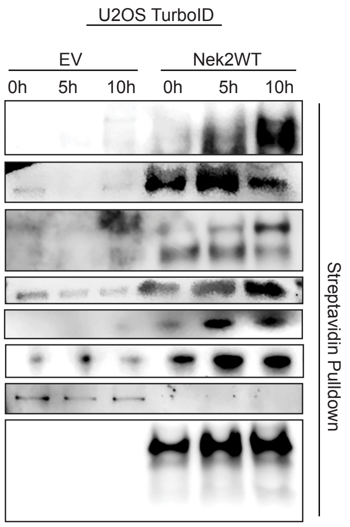


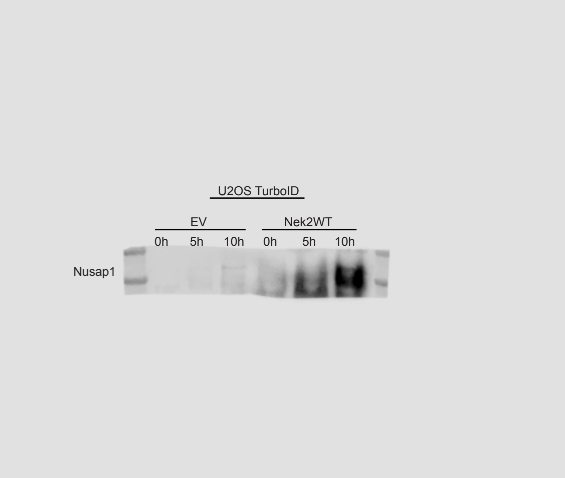


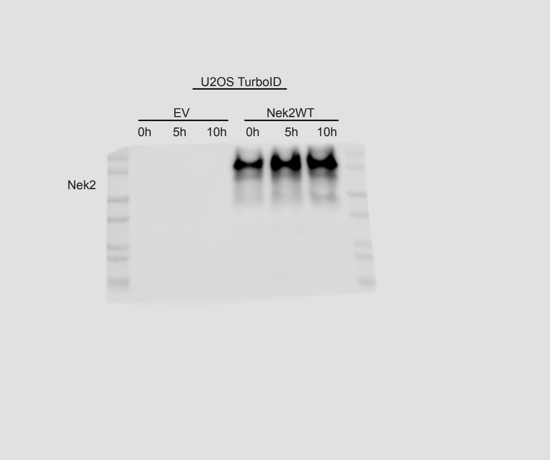


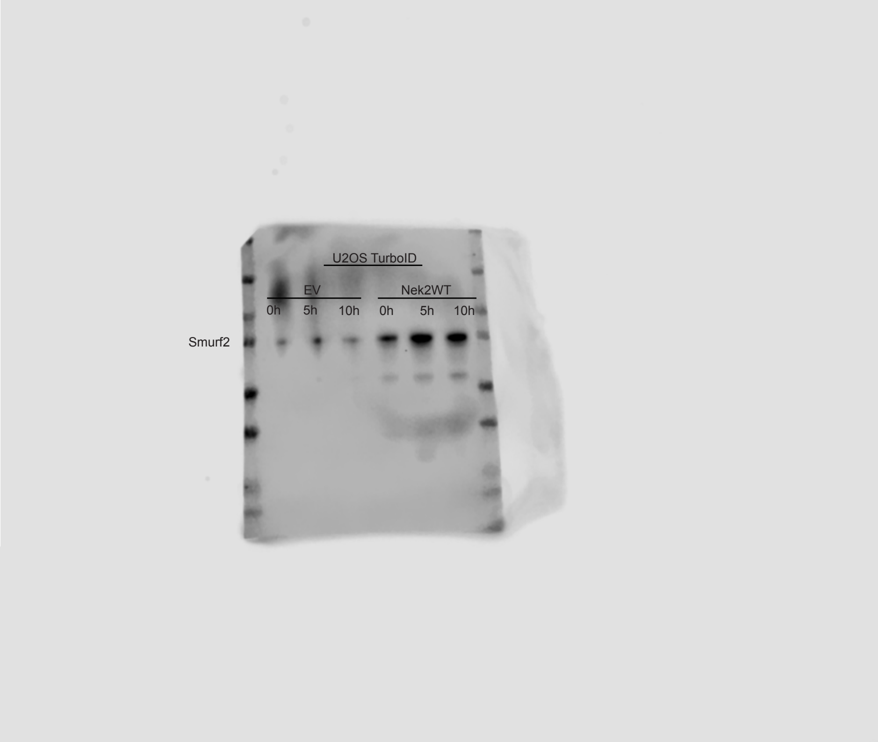


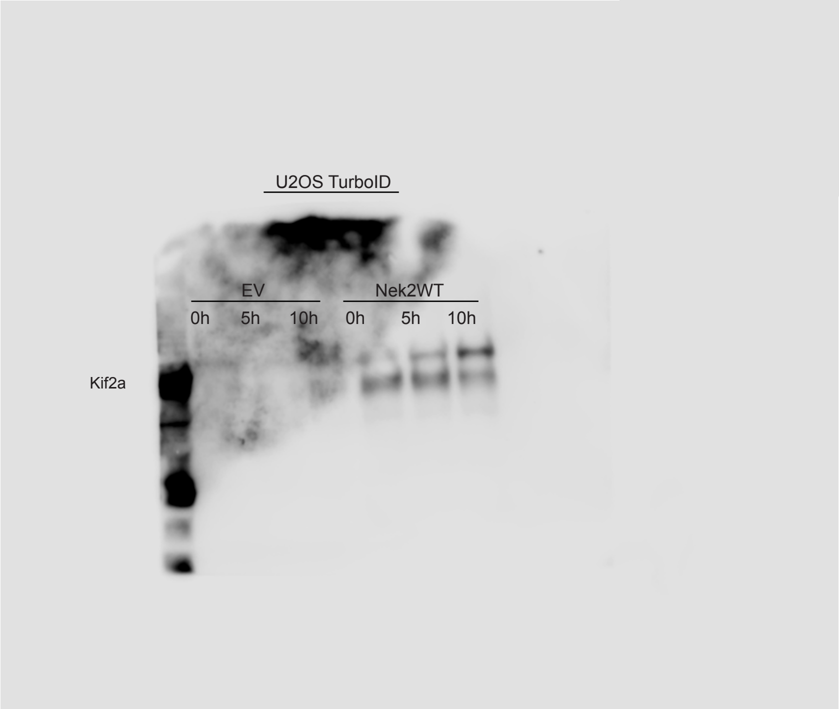

**4D**

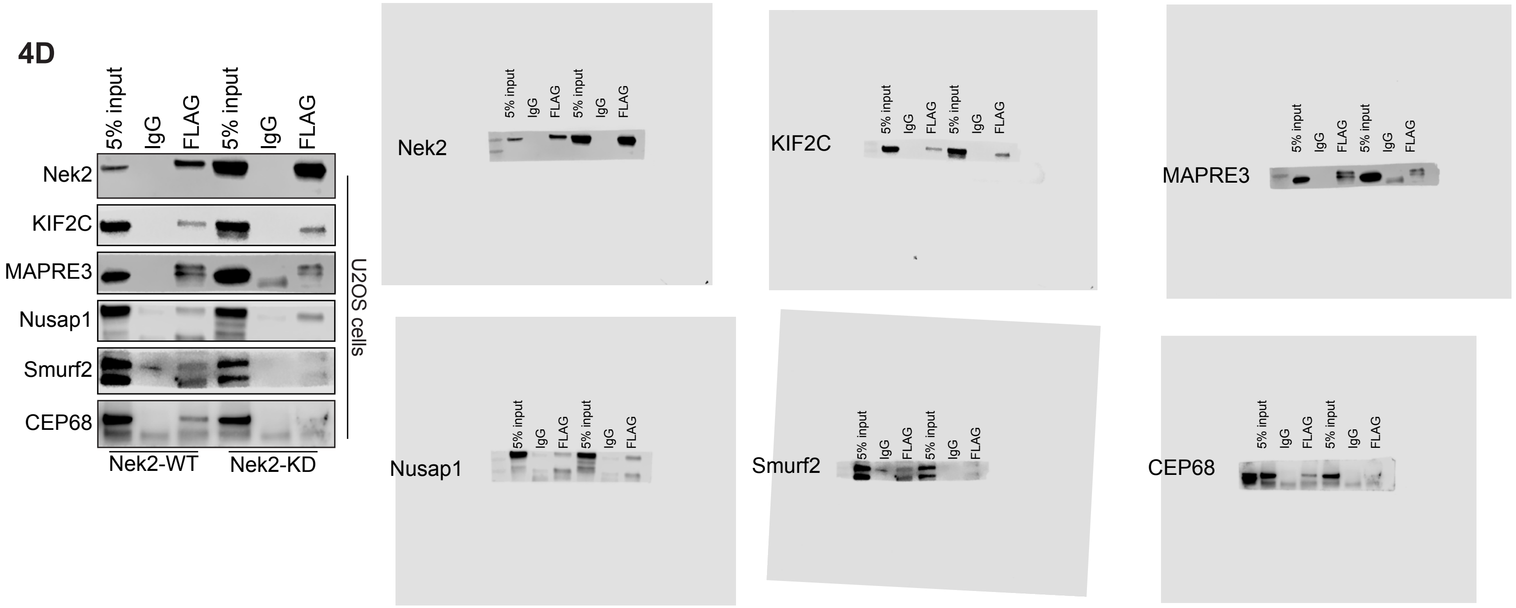
5% input
IgG
FLAG
5% input
IgG
FLAG
Nek2
KIF2C
MAPRE3
Nusap1
Smurf2
CEP68
U2OS cells
Nek2-WT
Nek2-KD

**4E**

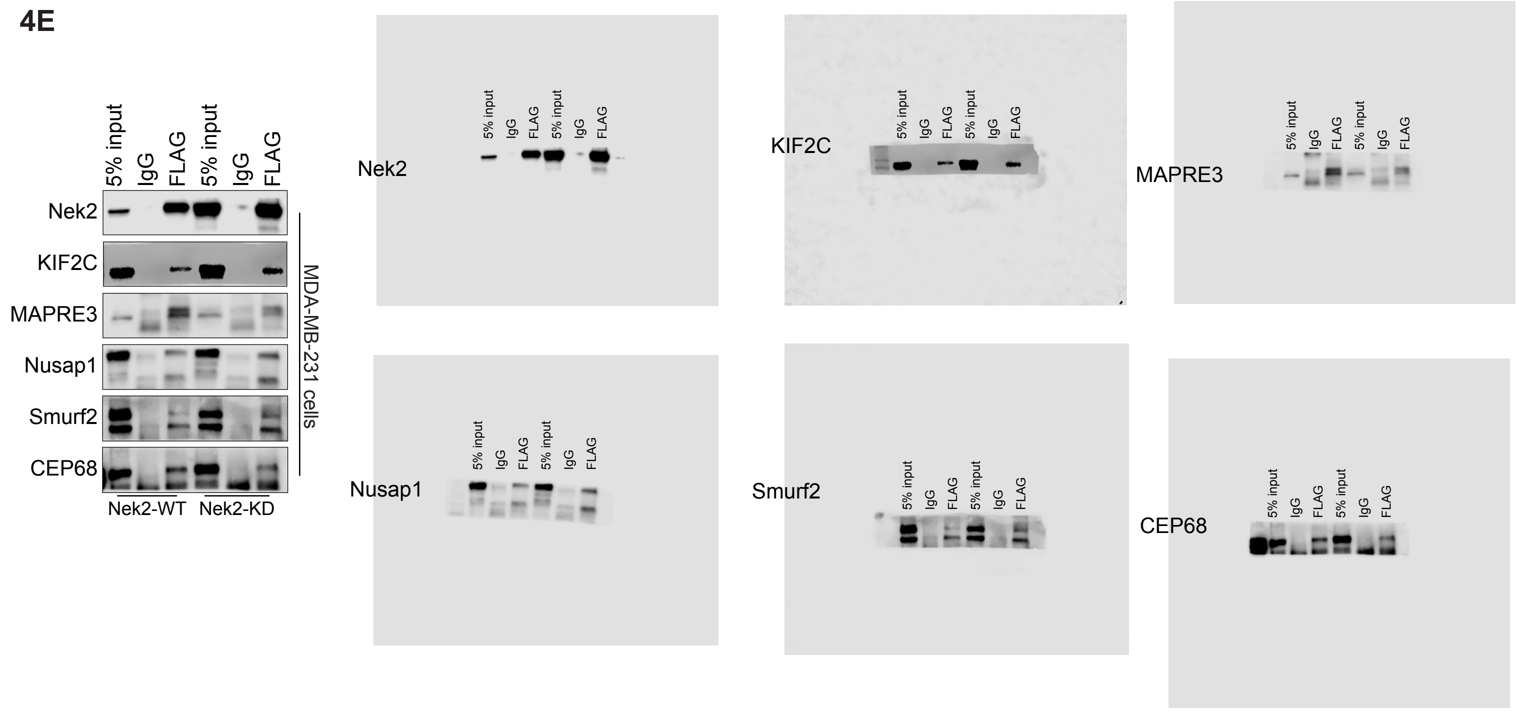
5% input
IgG
FLAG
5% input
IgG
FLAG
Nek2
KIF2C
MAPRE3
Nusap1
Smurf2
CEP68
MDA-MB-231 cells
Nek2-WT
Nek2-KD

**4F**

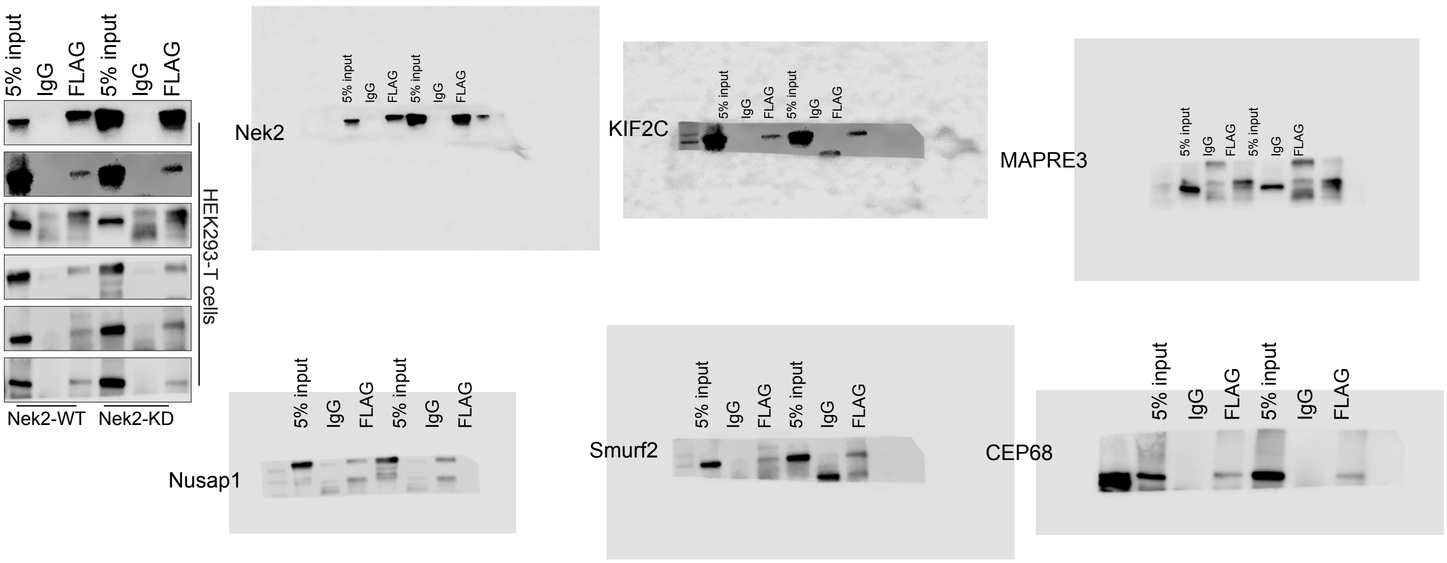
5% input
IgG
FLAG
5% input
IgG
FLAG
Nek2
KIF2C
MAPRE3
Nusap1
Smurf2
CEP68
HEK293-T cells
Nek2-WT
Nek2-KD

**S7A)**

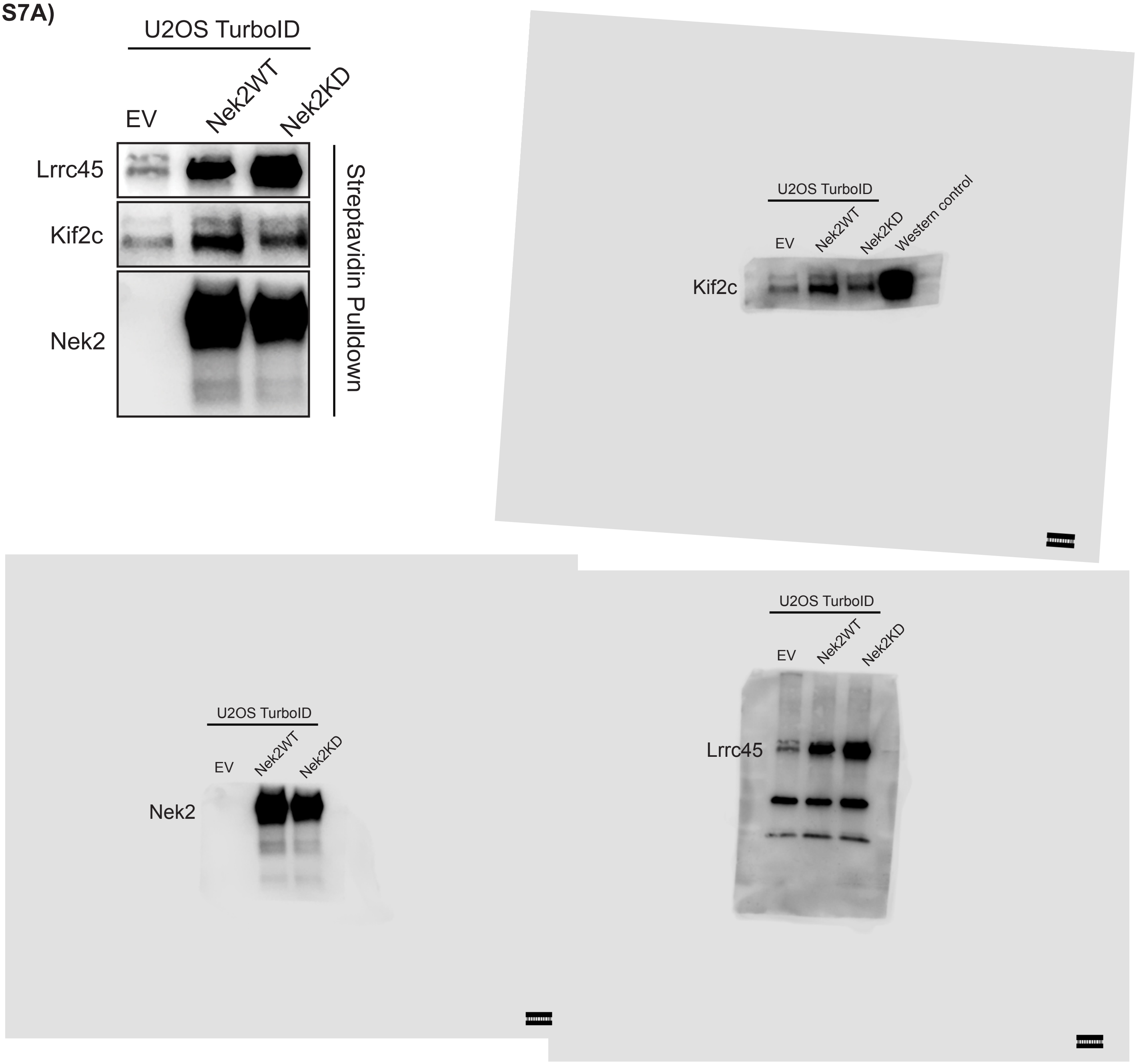
U2OS TurboID
EV
Nek2WT
Nek2KD
Lrrc45
Kif2c
Nek2
Streptavidin Pulldown
U2OS TurboID
EV
Nek2WT
Nek2KD
Western control
Kif2c
U2OS TurboID
EV
Nek2WT
Nek2KD
Nek2
U2OS TurboID
EV
Nek2WT
Nek2KD
Lrrc45

**S7D)**

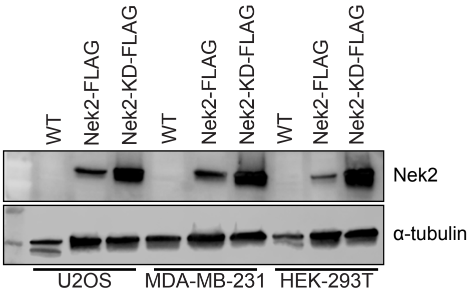
WT
Nek2-FLAG
Nek2-KD-FLAG
WT
Nek2-FLAG
Nek2-KD-FLAG
WT
Nek2-FLAG
Nek2-KD-FLAG
Nek2
α-tubulin
U2OS
MDA-MB-231
HEK-293T

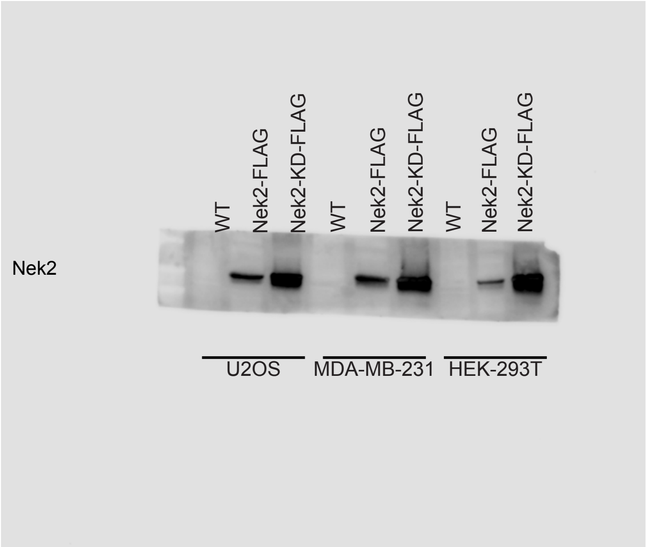
WT
Nek2-FLAG
Nek2-KD-FLAG
WT
Nek2-FLAG
Nek2-KD-FLAG
WT
Nek2-FLAG
Nek2-KD-FLAG
Nek2
U2OS
MDA-MB-231
HEK-293T

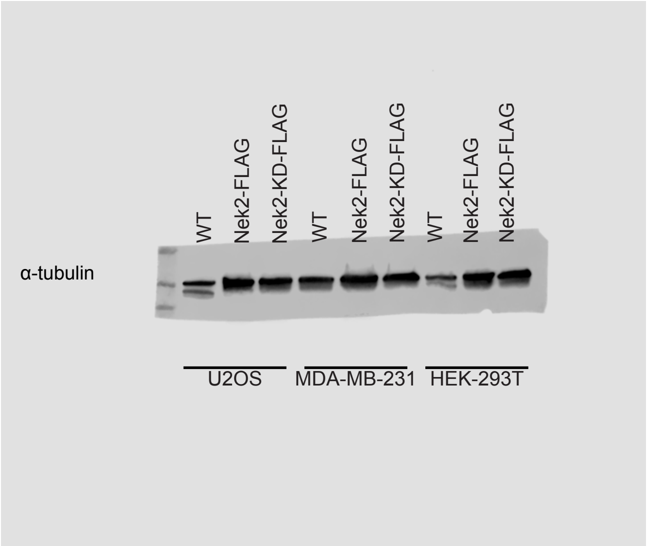
WT
Nek2-FLAG
Nek2-KD-FLAG
WT
Nek2-FLAG
Nek2-KD-FLAG
WT
Nek2-FLAG
Nek2-KD-FLAG
α-tubulin
U2OS
MDA-MB-231
HEK-293T

**S7E)**

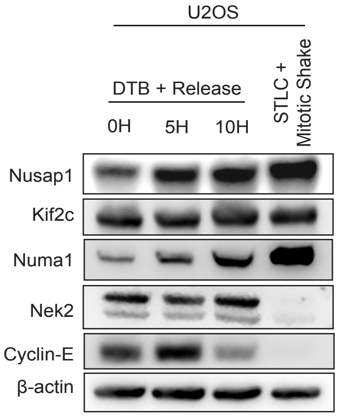
U2OS
DTB + Release
0H
5H
10H
STLC + Mitotic Shake
Nusap1
Kif2c
Numa1
Nek2
Cyclin-E
β-actin

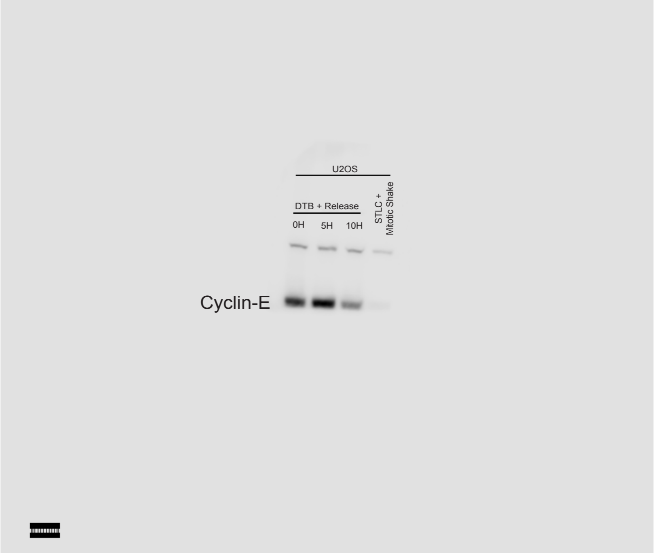
U2OS
DTB + Release
0H
5H
10H
STLC + Mitotic Shake
Cyclin-E

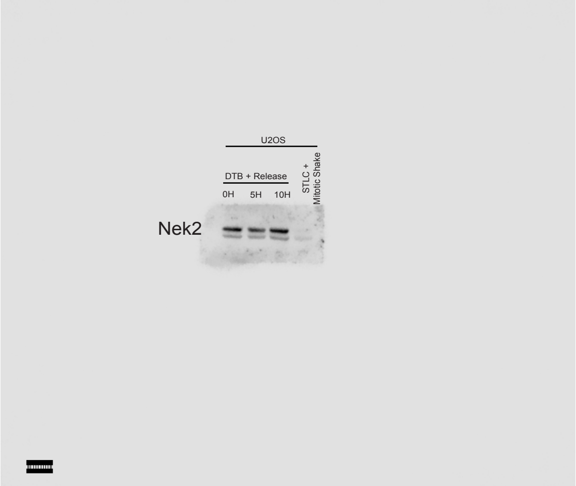
U2OS
DTB + Release
0H
5H
10H
STLC + Mitotic Shake
Nek2

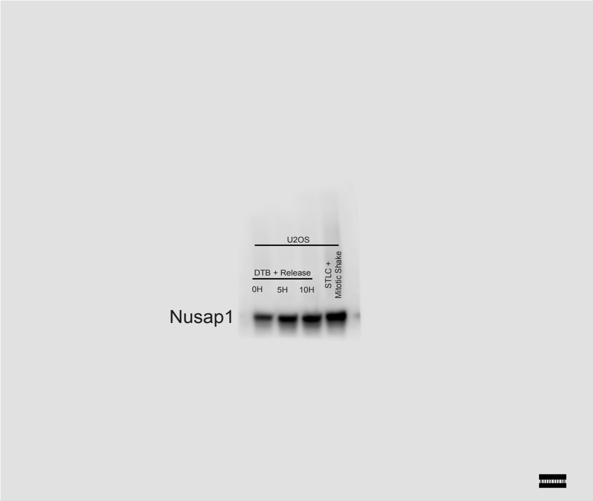
U2OS
DTB + Release
0H
5H
10H
STLC + Mitotic Shake
Nusap1

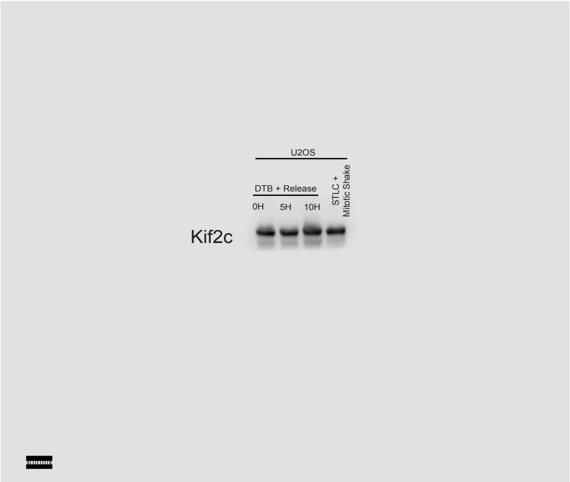
U2OS
DTB + Release
0H
5H
10H
STLC + Mitotic Shake
Kif2c

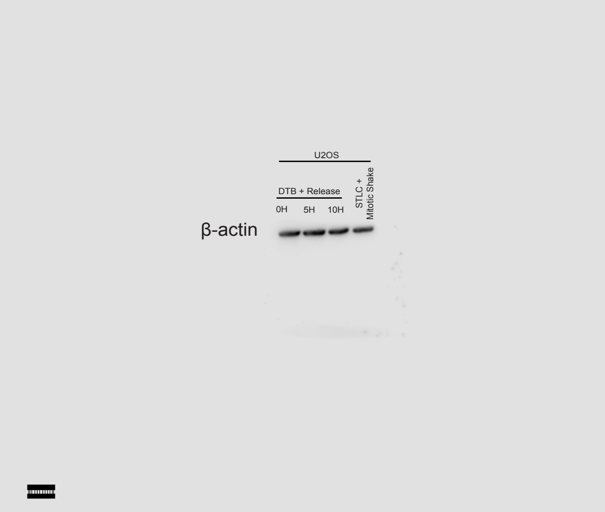
U2OS
DTB + Release
0H
5H
10H
STLC + Mitotic Shake
β-actin

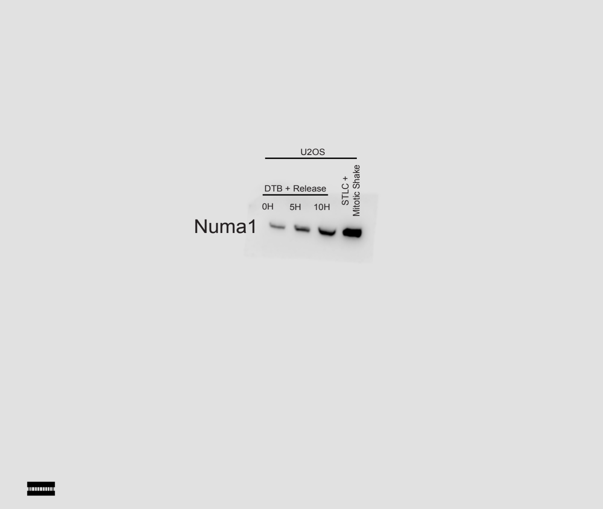
U2OS
DTB + Release
0H
5H
10H
STLC + Mitotic Shake
Numa1

5A)

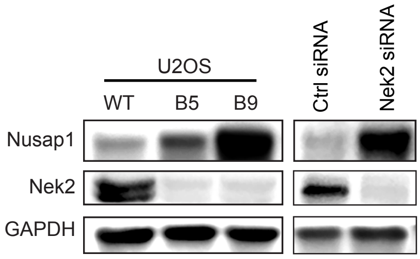
U2OS
WT
B5
B9
Ctrl siRNA
Nek2 siRNA
Nusap1
Nek2
GAPDH

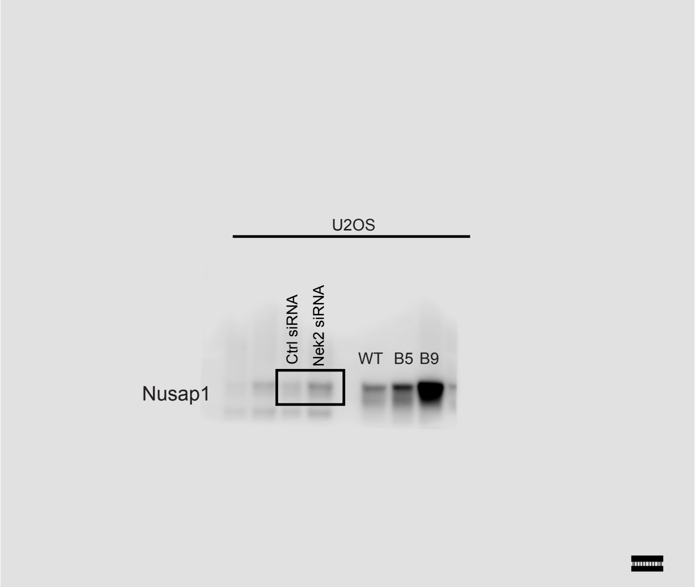
U2OS
Ctrl siRNA
Nek2 siRNA
WT
B5
B9
Nusap1

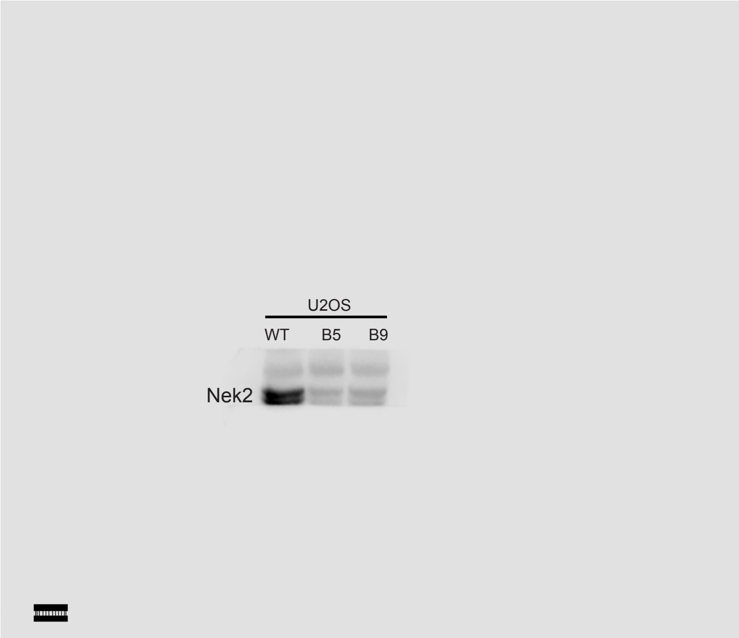
U2OS
WT
B5
B9
Nek2

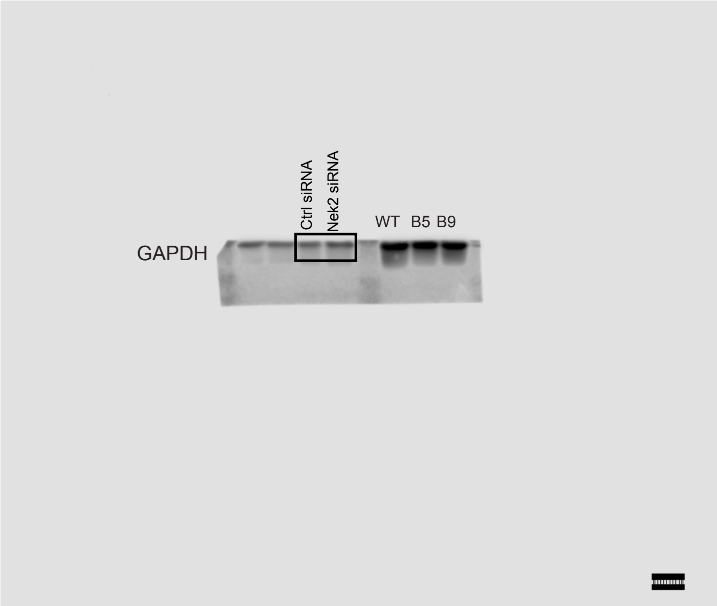
Ctrl siRNA
Nek2 siRNA
WT
B5
B9
GAPDH

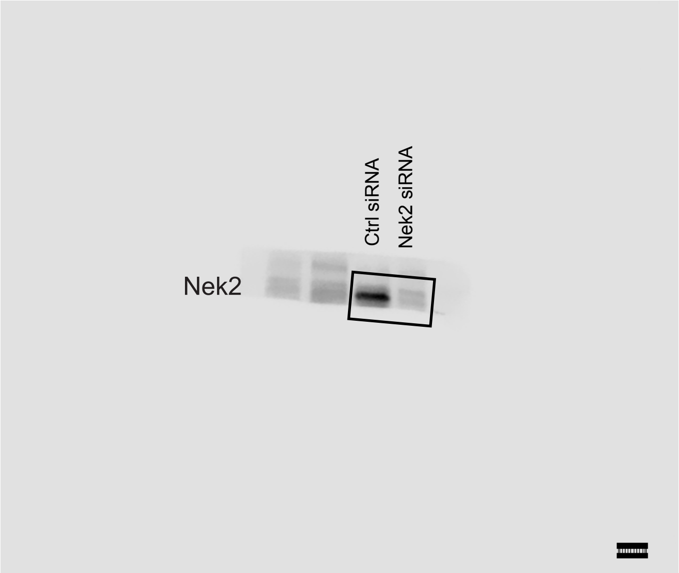
Ctrl siRNA
Nek2 siRNA
Nek2

**5B)**

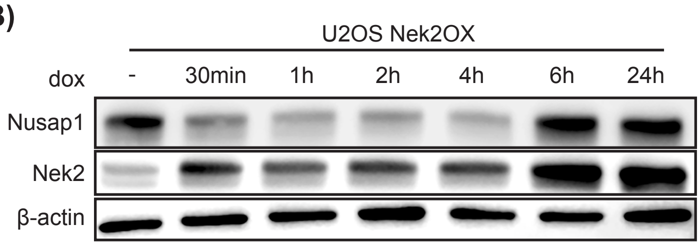
U2OS Nek2OX
dox
-
30min
1h
2h
4h
6h
24h
Nusap1
Nek2
β-actin

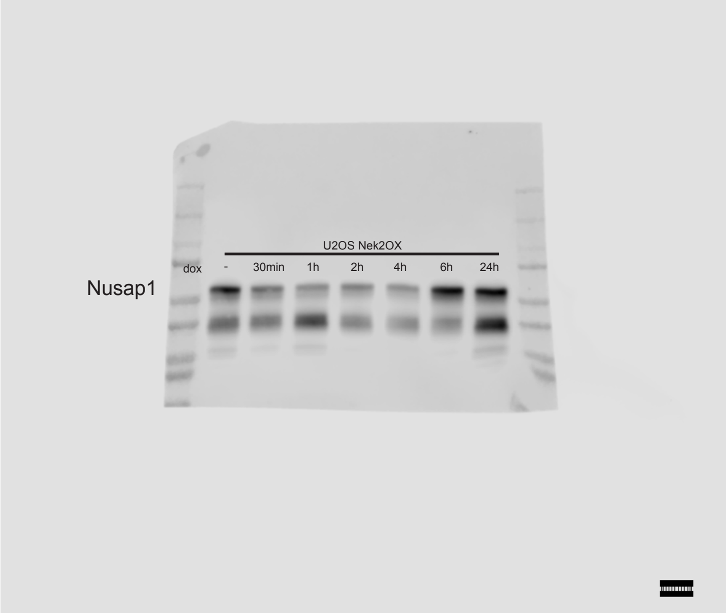
U2OS Nek2OX
dox
-
30min
1h
2h
4h
6h
24h
Nusap1

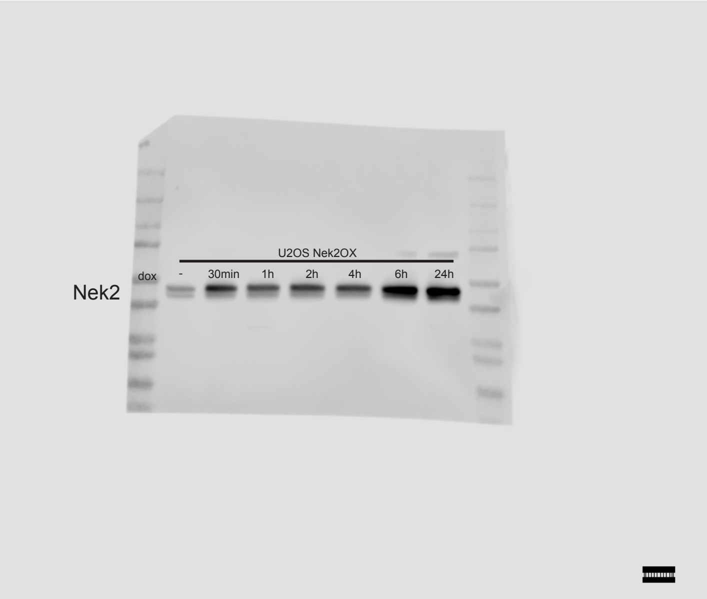
U2OS Nek2OX
dox
-
30min
1h
2h
4h
6h
24h
Nek2

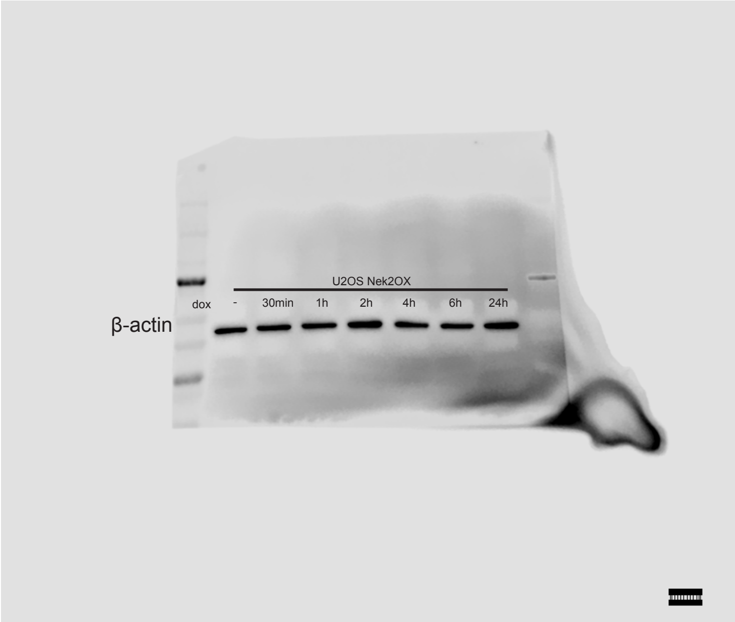
U2OS Nek2OX
dox
-
30min
1h
2h
4h
6h
24h
β-actin

**5C)**

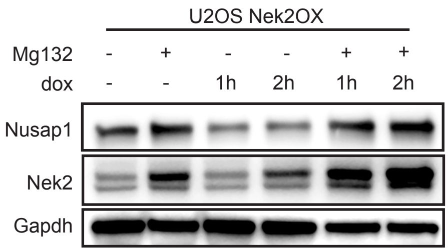

U2OS Nek2OX
Mg132 - + - - + +
dox - - 1h 2h 1h 2h
Nusap1
Nek2
Gapdh


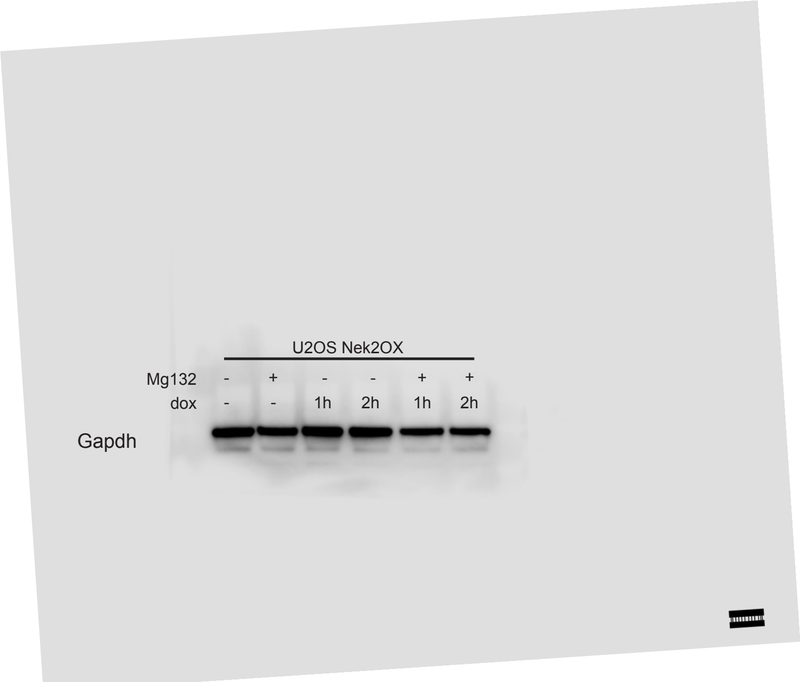

U2OS Nek2OX
Mg132 - + - - + +
dox - - 1h 2h 1h 2h
Gapdh


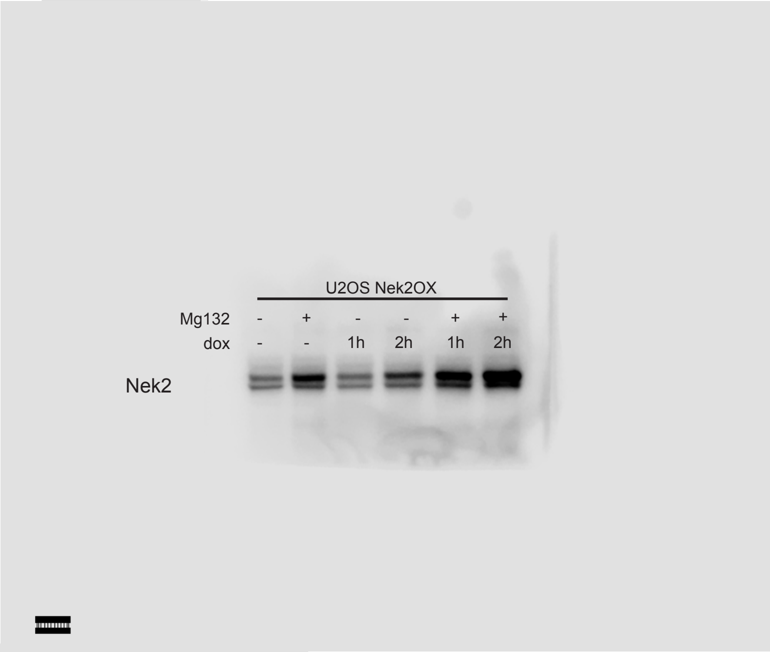

U2OS Nek2OX
Mg132 - + - - + +
dox - - 1h 2h 1h 2h
Nek2


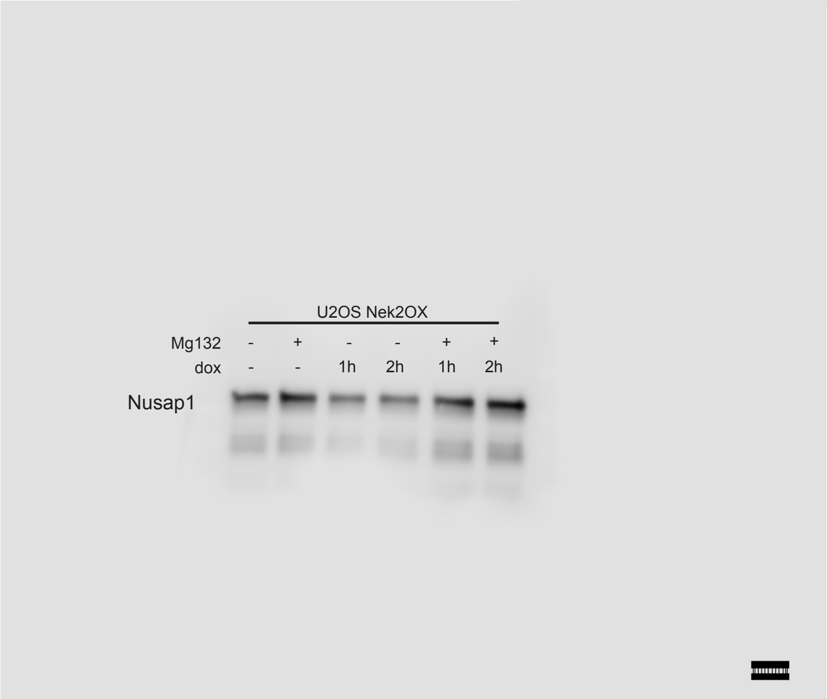

U2OS Nek2OX
Mg132 - + - - + +
dox - - 1h 2h 1h 2h
Nusap1

**5E)**

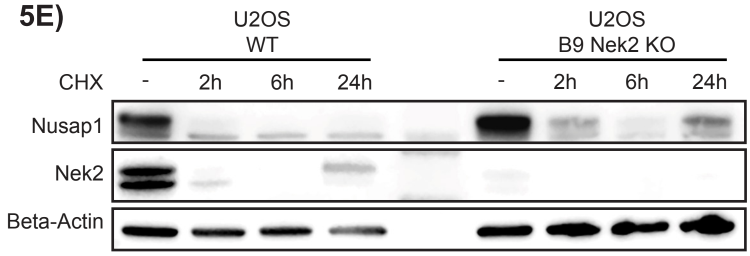
U2OS
WT
U2OS
B9 Nek2 KO
CHX
-
2h
6h
24h
-
2h
6h
24h
Nusap1
Nek2
Beta-Actin

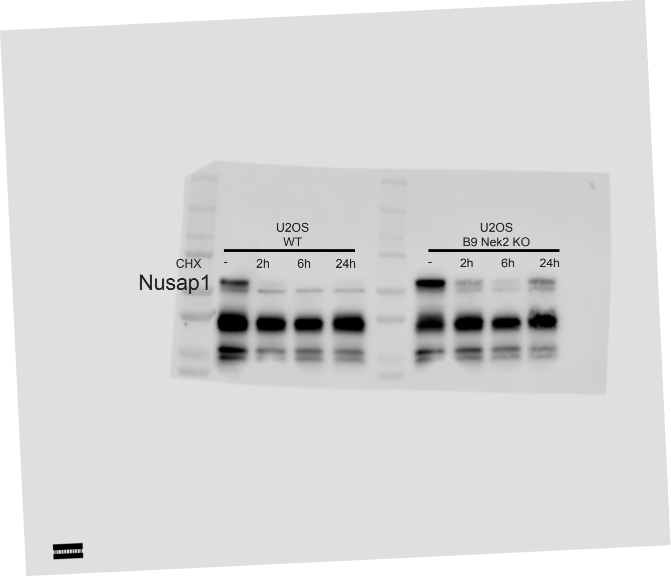
U2OS
WT
U2OS
B9 Nek2 KO
CHX
-
2h
6h
24h
-
2h
6h
24h
Nusap1

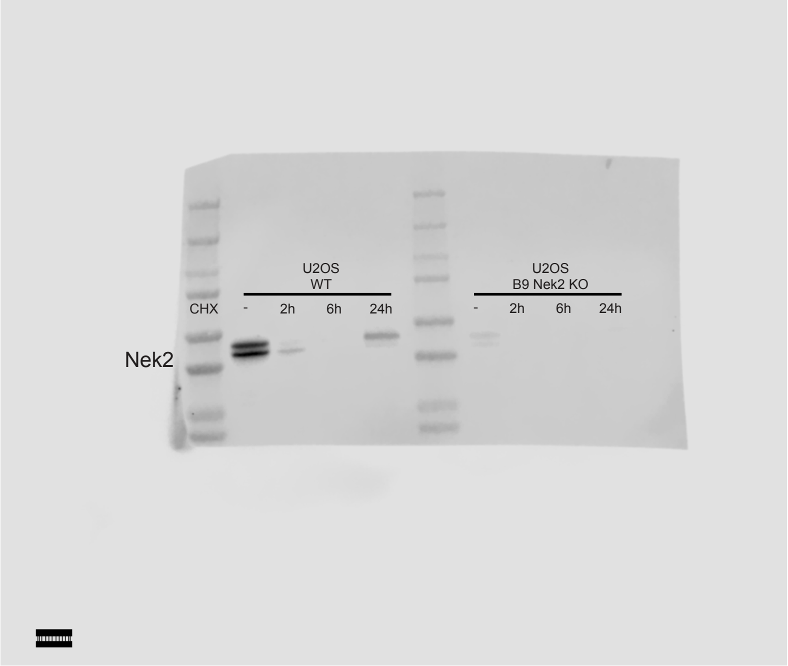
U2OS
WT
U2OS
B9 Nek2 KO
CHX
-
2h
6h
24h
-
2h
6h
24h
Nek2

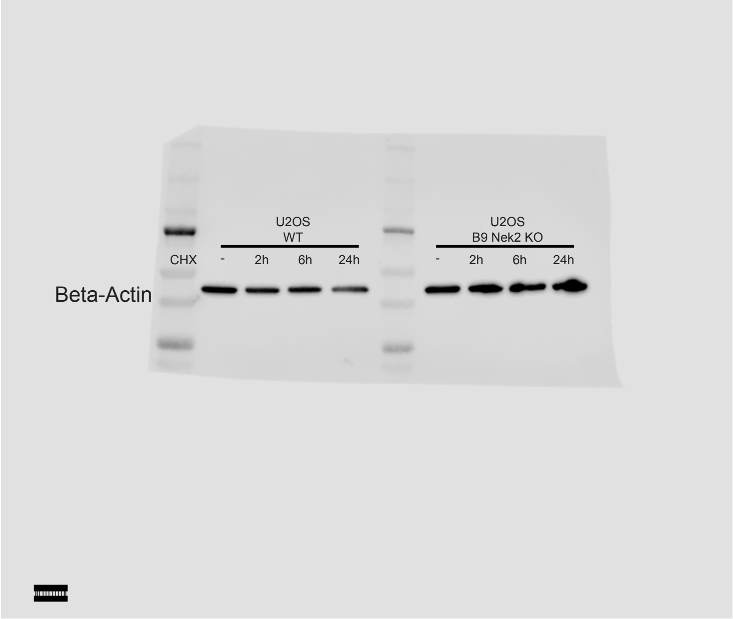
U2OS
WT
U2OS
B9 Nek2 KO
CHX
-
2h
6h
24h
-
2h
6h
24h
Beta-Actin

**5G)**

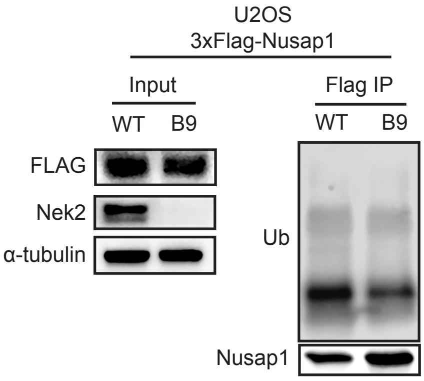
U2OS
3xFlag-Nusap1
Input
Flag IP
WT
B9
WT
B9
FLAG
Nek2
α-tubulin
Ub
Nusap1

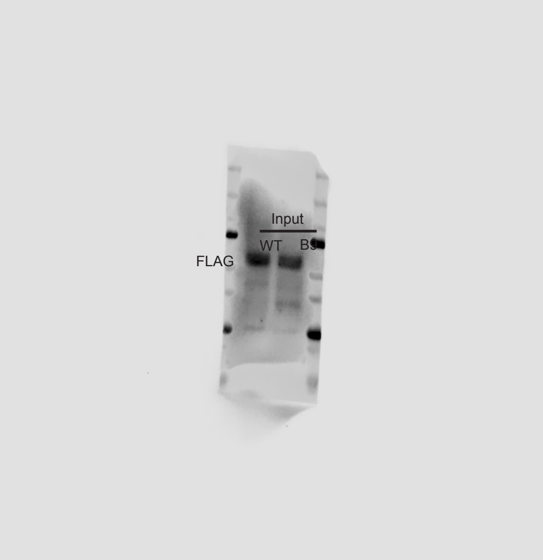
Input
WT
FLAG

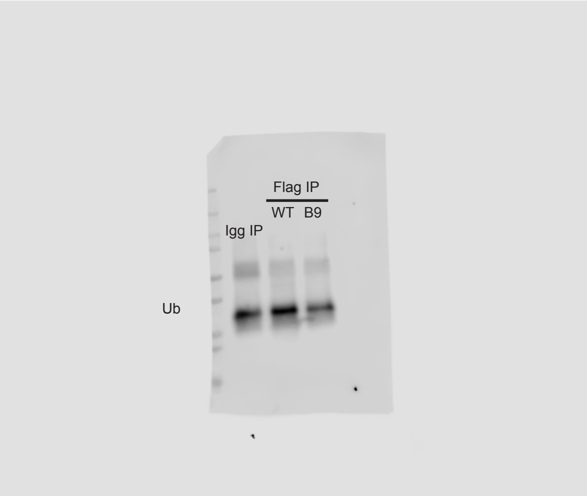
Flag IP
WT
B9
Igg IP
Ub

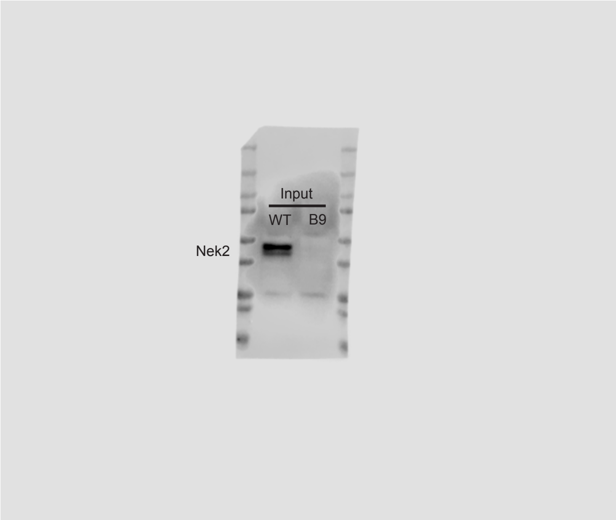
Input
WT
B9
Nek2

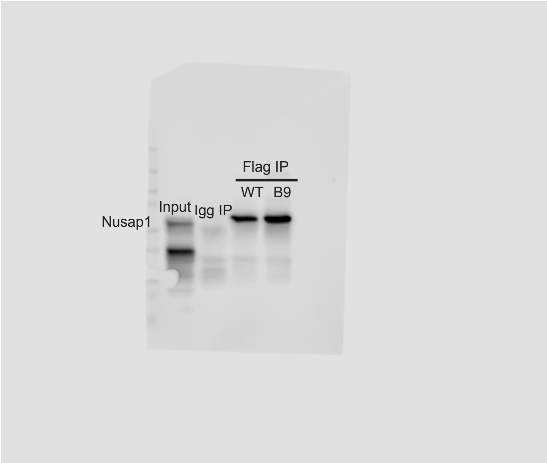
Flag IP
WT
B9
Input
Igg IP
Nusap1

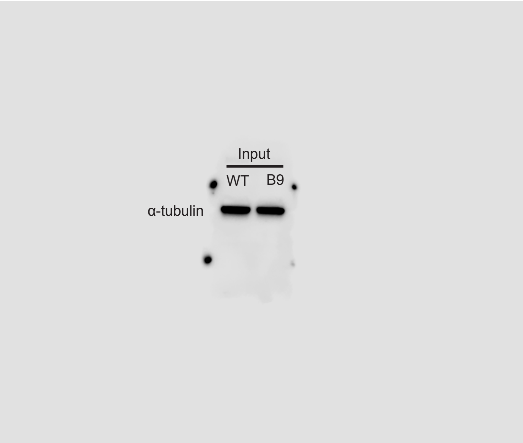
Input
WT
B9
α-tubulin

6C)

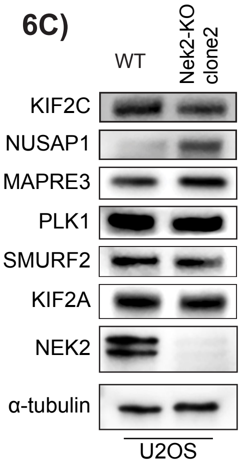
WT
Nek2-KO clone2
KIF2C
NUSAP1
MAPRE3
PLK1
SMURF2
KIF2A
NEK2
α-tubulin
U2OS

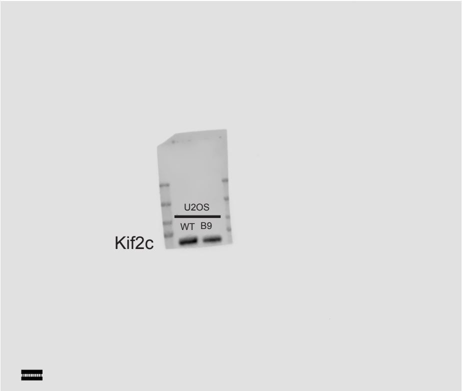
U2OS
WT B9
Kif2c

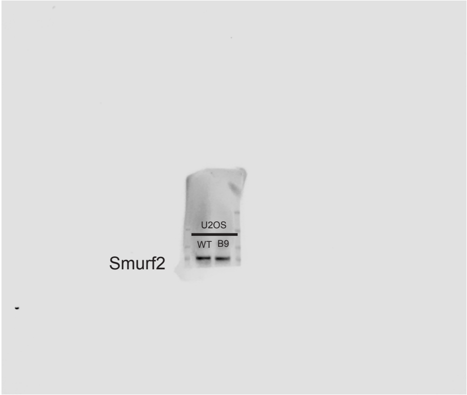
U2OS
WT B9
Smurf2

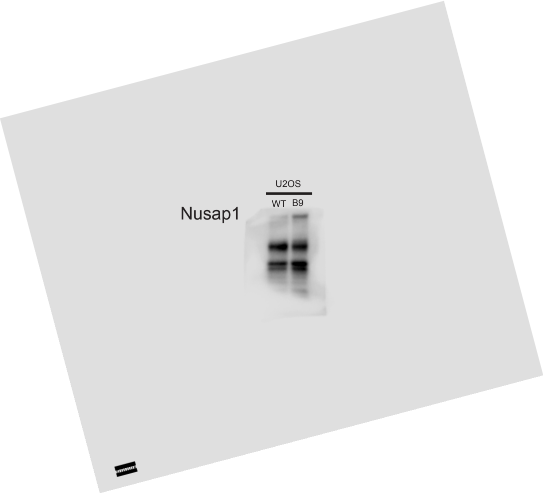
U2OS
WT B9
Nusap1

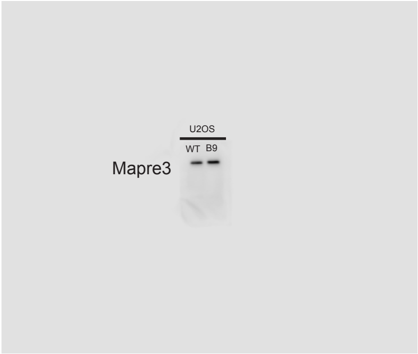
U2OS
WT B9
Mapre3

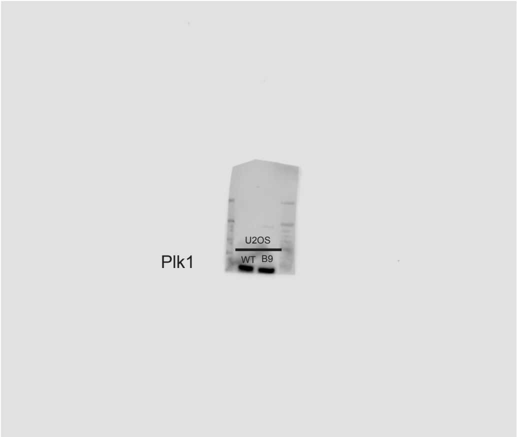
U2OS
WT B9
Plk1

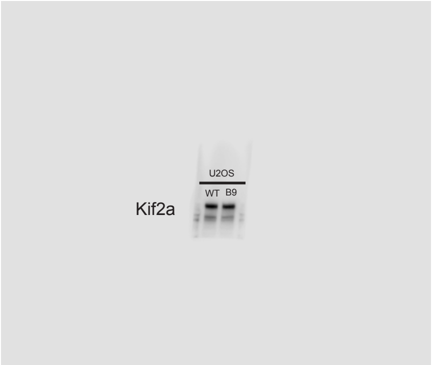
U2OS
WT B9
Kif2a

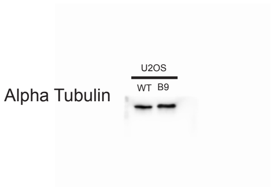
U2OS
WT B9
Alpha Tubulin

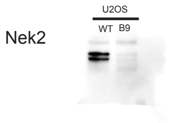
U2OS
WT B9
Nek2

**6C)**

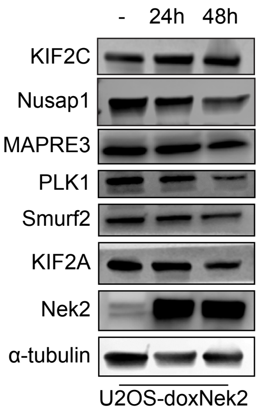

- 24h 48h
KIF2C
Nusap1
MAPRE3
PLK1
Smurf2
KIF2A
Nek2
α-tubulin
U2OS-doxNek2


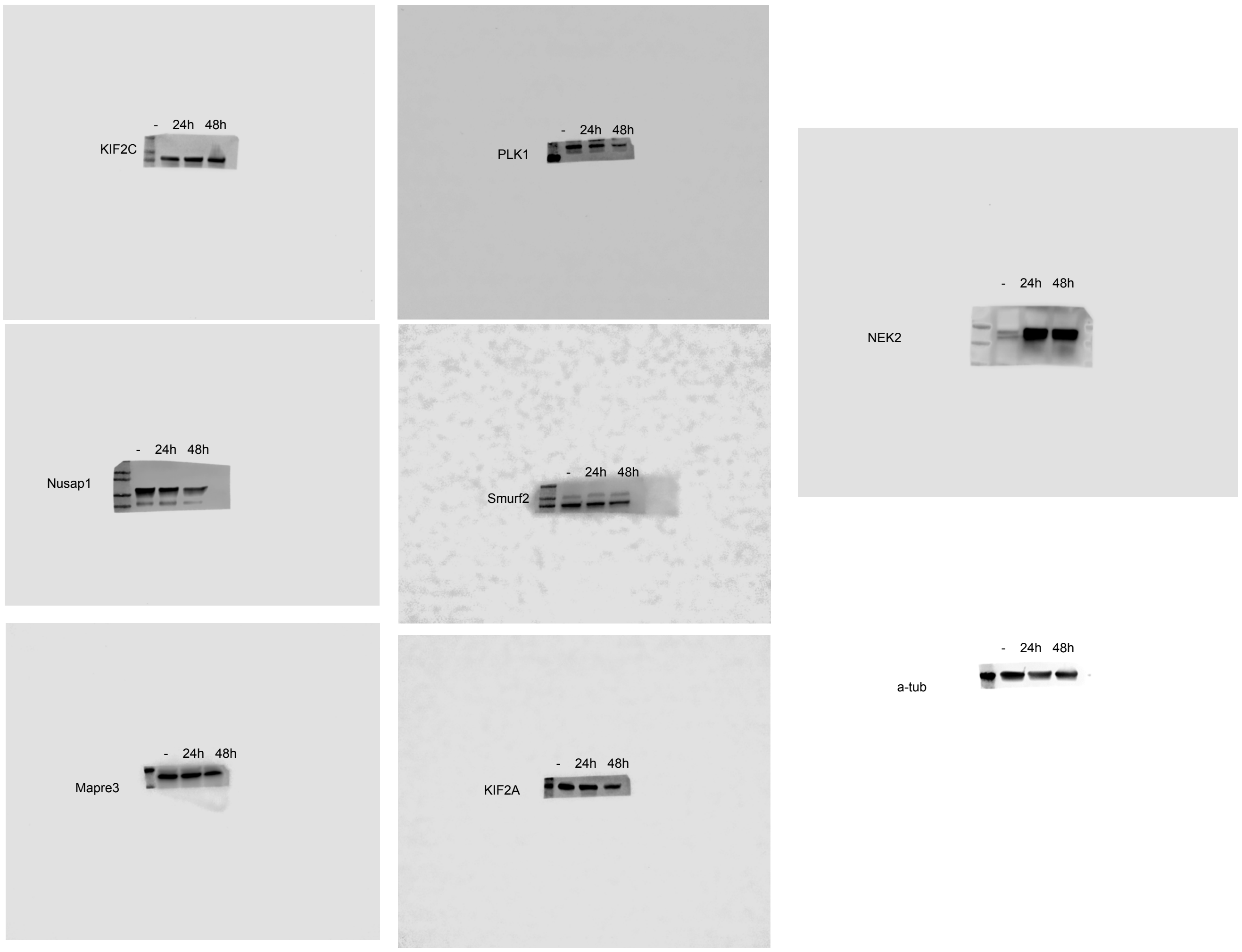

- 24h 48h
KIF2C
- 24h 48h
PLK1
- 24h 48h
NEK2
- 24h 48h
Nusap1
- 24h 48h
Smurf2
- 24h 48h
Mapre3
- 24h 48h
KIF2A
- 24h 48h
a-tub
